# Euchromatin forms condensed domains with short active regions on the surface

**DOI:** 10.1101/2025.08.05.668703

**Authors:** Joseph M. Paggi, Bin Zhang

## Abstract

Technological advances in chromatin structure characterization have continually refined our understanding of transcriptional regulation in eukaryotic systems. Despite these developments, achieving nucleosome-resolution structural characterization remains a significant challenge. As a result, it remains unclear what structural features distinguish active enhancers and promoters and how these elements are organized. To address this, we developed a simulation framework that leverages high-resolution Region-Capture Micro-C (RCMC) contact maps to infer conformational ensembles of megabase-scale chromatin segments at nucleosome resolution. A key component of this framework is a balancing strategy tailored for Micro-C data, which identifies per-nucleosome variation in contact density, in contrast to existing methods that assume uniform contact density. Our model accurately reproduces contact frequencies observed in RCMC data, pairwise spatial distances measured via chromatin tracing, and local structural motifs observed in imaging studies. The high spatial and genomic resolution of the inferred structures reveal a striking departure from the classical view of euchromatin as uniformly open. Instead, euchromatin generally folds into compact domains, consistent with the “packing domains” observed in imaging studies. These domains are further organized into disordered, insulated clusters of approximately 50 nucleosomes, resembling nucleosome clutches. Notably, kilobase-scale regions surrounding active promoters and enhancers often protrude from these condensed domains, becoming highly accessible. This spatial arrangement effectively compartmentalizes regulatory elements from the surrounding chromatin, facilitating protein binding and promoting enhancer–promoter communication. The distinct structural features of euchromatin revealed here offer new insights into enhancer regulation and may help explain their enigmatic behavior.

## Introduction

The textbook view of genome organization posits that transcriptionally inactive heterochromatin adopts closed conformations, whereas active euchromatin adopts open ones. In this model, transcriptional machinery can access only euchromatin, leading to selective gene expression. However, recent evidence challenges this dichotomy, suggesting that euchromatin also contains largely condensed domains [1–4]. Accurately quantifying the compactness of euchromatin and resolving its three-dimensional (3D) organization—particularly at the nucleosome level near promoters—could yield critical insights into the mechanisms of transcriptional regulation. Achieving this, however, demands both high spatial and genomic resolution—an ongoing technological challenge.

Advances in imaging have substantially deepened our understanding of chromatin organization. Techniques such as super-resolution microscopy [5] and electron microscopy [1, 6–10] have reached single-nucleosome resolution and uncovered structural features ranging from packing domains [1, 11, 12] to nanodomains [13] and clutches [5, 14]. However, these methods generally have limited ability to identify the specific genomic loci being imaged and to trace the chain of nucleosomes, leaving critical questions unanswered. In particular, the hierarchical relationships among structural motifs—e.g., how clutches (tens of nucleosomes) aggregate into packing domains (thousands of nucleosomes)—and how these features depend on genomic context remain elusive. It also remains unclear whether condensed domains identified through imaging correspond to active or inactive chromatin states. Chromatin tracing methods [12, 15, 16] can target specific genomic segments, but lack the resolution required to resolve single-nucleosome features and address these questions.

Chromatin conformation capture and related methods offer complementary advantages by capturing spatial proximity between DNA segments in situ [17–21]. Micro-C, in particular, can achieve high genomic resolution [22–25]. Nonetheless, interpreting contact maps remains challenging due to the confounding effects of population averaging and the inherently indirect relationship between contact frequency and 3D structure. Computational approaches [26–42], such as maximum entropy inversion (MEI) [43–47], have made progress in addressing these challenges by reconstructing 3D conformational ensembles consistent with average contact maps. However, the limited sequencing depth typical of standard Micro-C experiments continues to constrain the resolution of the inferred structures.

Region-capture Micro-C (RCMC) mitigates this limitation by concentrating sequencing depth on selected genomic regions [48–50]. This approach dramatically enhances resolution and has enabled the generation of high-quality, nucleosome-resolution contact maps. These maps have already revealed previously unknown structural motifs, such as micro-compartments, and support modeling efforts aimed at reconstructing nucleosome-resolution 3D structures.

In this study, we apply MEI to RCMC data to reconstruct nucleosome-resolution conformational ensembles. To enable this, we developed a balancing strategy, “neighbor balancing,” which recovers variations in contact density. Our simulations accurately reproduces both RCMC contact maps and pairwise distances from chromatin tracing. The simulated conformations contain structural motifs consistent with packing domains (or TAD-like domains) [1, 12] and clutches [5]. In this way, our results provide a coherent, multiscale view of chromatin architecture—from single nucleosomes to megabase domains—bridging structural motifs observed across diverse techniques. Notably, our analysis reveals a surprising similarity in the overall compaction of euchromatin and heterochromatin. At the same time, we identify euchromatin-specific structural features, including an organization in which compact chromatin domains are folded internally, exposing regulatory regions on the surface. We discuss the functional implications of this architecture in the context of enhancer–promoter communication and transcriptional regulation.

## Results

### Neighbor balancing recovers biological variation in contact density

Iterative correction (ICE) is a widely used balancing technique for Hi-C and micro-C data [51]. The role of balancing is to account for biases in read coverage in different genomic windows, including digestibility by restriction enzymes or MNase and PCR amplification biases. Without balancing, genomic regions with intrinsically higher coverage would have inflated contact frequencies.

ICE balancing and other popular approaches [52] remove variations in coverage by enforcing that each genomic window is assigned the same contact density: the sum of its contact frequencies with all other windows. There is a natural relationship between the contact density and the physical density of the system (Figure 1a). If we assume a model in which a micro-C contact is likely if two nucleosomes are within some “capture radius” of each other and unlikely otherwise [53], then the ensemble average physical density is equal to the contact density divided by the volume enclosed by the capture radius. In this way, the assumption of constant contact density is equivalent to an assumption of constant average physical density.

**Figure 1:**
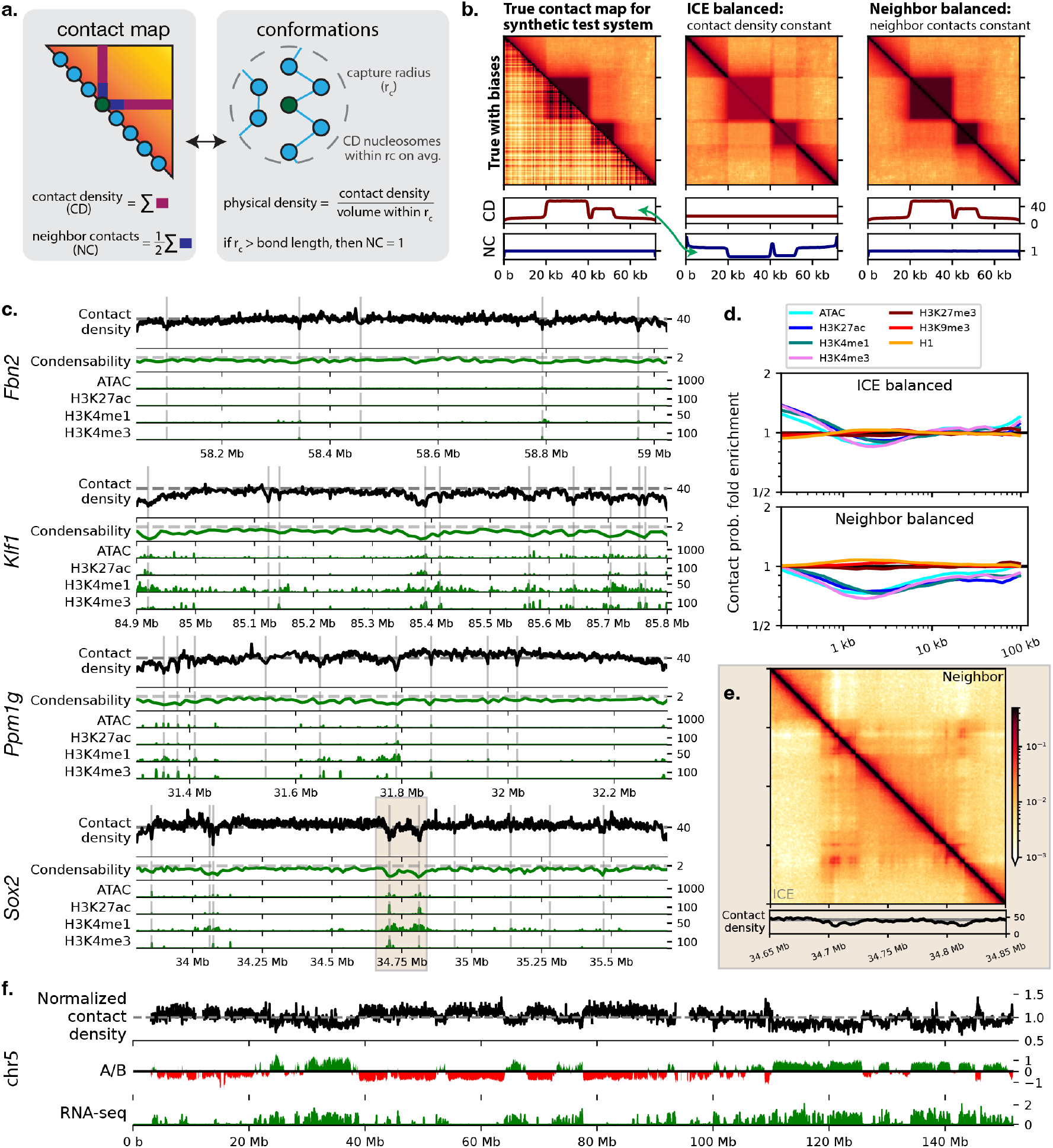
Neighbor balancing recovers variation in contact density. (a) Illustration of the relationship between contact maps and the underlying conformational ensemble. (b) Contact maps for a synthetic test system composed of two densely packed regions connected by an extended linker. The left panel shows the true contact map on the upper-triangle and the true contact map corrupted by multiplicative biases drawn from a log-normal distribution in the lower-triangle. The per-nucleosome contact density and neighbor contacts for the true map are shown at the bottom. The center and right panels show the ICE and neighbor balanced maps, respectively, computed starting from the corrupted contact map. (c) Comparison of contact densities inferred using neighbor balancing for RCMC data from Goel et al. [48] to condensability scores [54], ATAC-seq coverage, and several active histone marks. The sources for all epigenetics data are provided in Supplementary Table 2. (d) Enrichment of contact probabilities for windows in the top 5% of the indicated epigenetic feature over windows in the bottom 50%. (e) Comparison of ICE balanced and neighbor balanced contact maps for the interaction between the *Sox2* promoter and one of its enhancers. (f) Comparison of contact densities for the entirety of chr5 computed using genome-wide micro-C data from Hsieh et al. [25] to A and B compartment scores and RNA-seq coverage (log10(1 + reads per genomic content)). All genome-wide data are at 25.6 kb resolution (see Supplementary Information).

When applied to systems that do not have a uniform physical density, ICE balancing erases the differences, artificially inflating contact frequencies for low density regions and decreasing them for high density regions. This likely results in problems in real systems because some regions of the genome, like acetylated regions, are believed to be more open than others [54, 55]. We demonstrate this effect using a synthetic system consisting of two dense domains separated by an extended linker (Figure 1b). After simulating this system and producing an ICE balanced contact map, the contact frequencies inside the dense domains are too low while those for the linker regions are too high, relative to the ground truth.

We propose “neighbor balancing” to recover variation in contact density. Unlike ICE, which assumes uniform contact density across all nucleosomes, our approach is based on the assumption that neighboring nucleosomes are consistently in contact. The intuition behind our approach is that, if neighboring nucleosomes are generally close enough that they have a high probability of being ligated in micro-C, then the contact frequencies between neighbors should all be the same. This assumption is supported by our analysis in the next section.

To enforce uniform contact frequency between neighboring nucleosomes, we re-balance the ICE-balanced contact map based on local interaction patterns. Specifically, for each nucleosome, we compute the average of its interaction frequency with its two neighbors then divide each row and column by the square root of this average (see Methods). In our synthetic example, we observe that in the ICE balanced contact map the neighbor contact frequencies are not constant and instead are inversely proportional to the true contact density (Figure 1b, blue lines). By normalizing against these neighbor interactions, our method effectively restores true variations in contact density. On the synthetic example system, neighbor balancing correctly reproduces the true contact densities.

We applied neighbor balancing to RCMC data for five genomic regions in mouse embryonic stem cells (mESCs) at nucleosome resolution (200 bps) [48]. Most nucleosomes have approximately the same contact density, except for short regions with as little as half the average density (Figure 1c, Fig. S3). These dilute positions are highly correlated with condensability scores [54], ATAC-seq, H3K27ac, H3K4me3, and other marks associated with active chromatin (Figure 1c, Fig. S3).

ICE balancing predicts that nucleosomes with active marks have high contact probabilities with nucleosomes within 1 kb, suggesting that they are locally compact, whereas neighbor balancing predicts that these regions are expanded at all scales (Figure 1d). The trend observed with neighbor balancing is in line with a variety of data suggesting that such regions are more extended and form fewer contacts [56–58]. Conversely, inactive marks, including H3K9me3, H3K27me3, and histone H1, are positively correlated with contact density although this trend is much weaker (Figure 1d). Since regions around promoters and enhancers tend to have lower contact density, the contact frequencies between enhancers and promoters are reduced, however, they remain visible in the contact map (Figure 1e, Fig. S4).

Neighbor balancing is not specific to RCMC and can be applied to typical, genome-wide micro-C (see Supplementary Information and Methods). We applied neighbor balancing to compute contact densities using genome wide micro-C in mESCs at a resolution of 25.6 kb [25]. The contact density is relatively consistent: nearly all positions fall within 50% of the average (Figure 1f). We observed similar correlations between contact densities and epigenetic marks as in the RCMC data (Fig. S5e) and that contact densities are inversely correlated with A/B compartment scores and RNA-seq coverage (Figure 1f). Neighbor balancing suggests that B compartment regions are roughly 20% denser than A compartment regions [59].

### Nucleosome-resolution chromatin modeling bridges contact maps with chromatin tracing

While the RCMC contact maps and contact densities from neighbor balancing begin to suggest what distinguishes active from inactive chromatin, they do not provide the complete picture. We therefore used maximum entropy inversion (MEI) to recover a 3D conformational ensemble consistent with the contact maps (Figure 2). In MEI, one begins with a baseline model and then adds auxiliary energy terms to make the system recapitulate experimental data (see Methods). Thereby, to apply MEI, we needed to define the baseline model and a way to compute a simulated micro-C contact map from a set of simulation frames.

**Figure 2:**
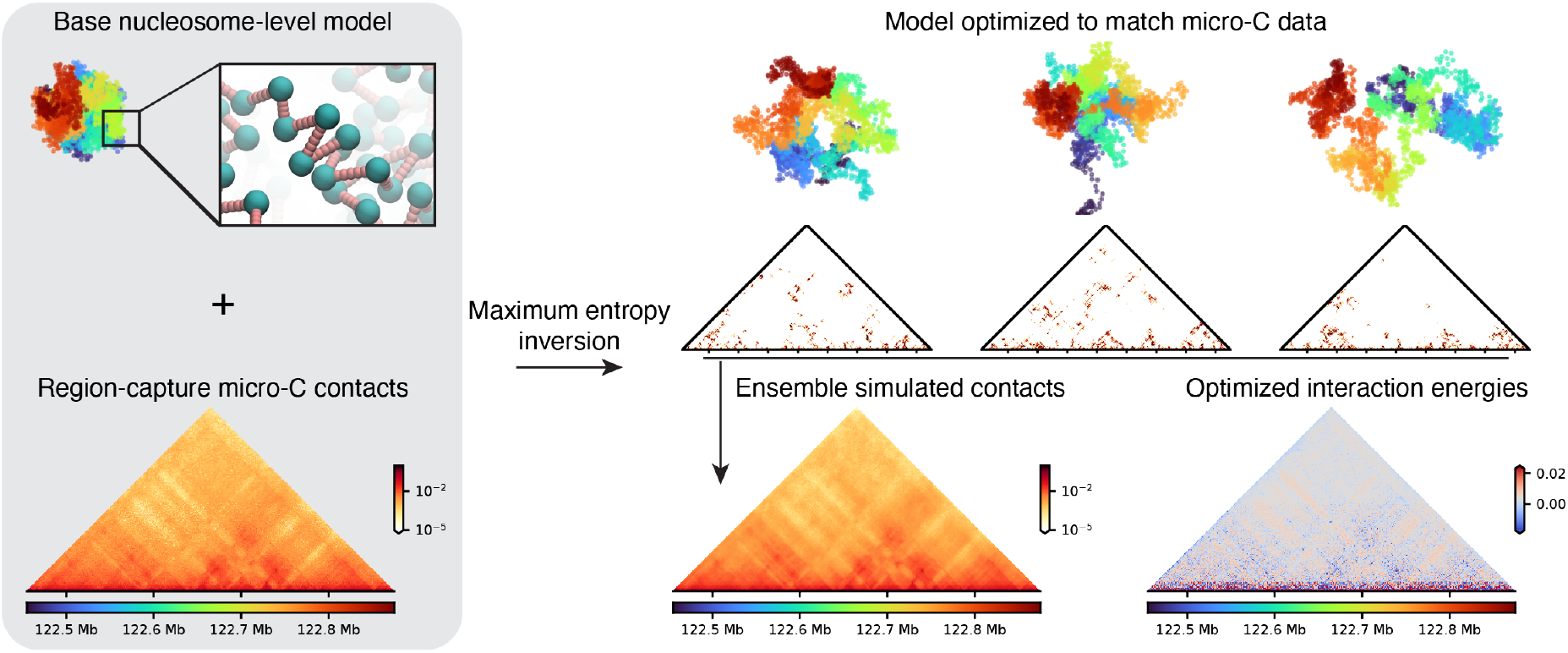
Data-driven modeling of chromatin at nucleosome resolution. In maximum entropy inversion (MEI), we begin with a base model of the system and an experimentally determined contact map. In the base model, all nucleosomes interact uniformly, and thereby the features in the contact map are not reproduced. MEI updates the nucleosome interaction potentials so that the simulations closely recapitulate the experimental contact map. This example uses RCMC data from the *Nanog* region [48].

Selecting a baseline model posed two competing requirements. Using a nucleosome resolution model appeared to be important to accurately capture the behavior of the short dilute regions detected by neighbor balancing (Figure 1c). The differences in contact density are diminished if viewed at coarser resolution (Fig. S5d), and we reasoned that the dilute regions could have different local structures. At the same time, it was essential to simulate at least megabase-scale systems to capture complete topologically associating domains (TADs). Relatively large genomic windows are required to include the majority of contacts; on average to include 75% of a nucleosome’s contacts requires simulating a 1 MB region (Fig. S5a,b). To meet these two criteria, we used a highly simplified nucleosome-level model, in which nucleosomes are positioned uniformly along the DNA and interact uniformly with an isotropic potential (see Methods).

Computing a simulated contact map requires specifying the likelihood of two nucleosomes forming a micro-C contact as a function of their distance. We call this function the “contact indicator function” and it’s 50% point the “capture radius”. The choice of contact indicator function strongly influences the overall scale of the optimized system: smaller capture radii lead to more compact systems and larger capture radii lead to more extended systems. To choose a contact indicator function and generally validate our approach, we compare our simulations to chromatin conformations observed using Optical Reconstruction of Chromatin (ORCA) [12]. ORCA uses fluorescent probes to sequentially image regions along the chromatid, providing 3D coordinates of specific genomic regions in thousands cells. ORCA data is available for two of the regions for which we have RCMC data: *Nanog* and *Sox2* [60, 61]. The *Nanog* data has a 140 kb region tiled at 15 kb resolution and the *Sox2* data has a 1.6 MB region tiled at 30 kb resolution.

Using a capture radius of 42 nm (two bond lengths in our model), the overall scaling of pairwise distances as a function of genomic separation in our simulations closely matches the ORCA experiments, after accounting for measurement uncertainty (Figure 3a,c, see Supplementary information). Moreover, the average pairwise distances between specific genomic regions are also well matched (r=0.89 for *Nanog*, r=0.92 for *Sox2*, Figure 3b,d). This correlation is greater than what is expected purely due to genomic separation, as illustrated by distance striated correlation coefficients (Figure 3b,d). The only substantial disagreement between our simulated ensemble and the chromatin tracing data is that the simulations underestimate median distances at short genomic separations (Figure 3a). We suspect this occurs because our simulations assume full nucleosome occupancy, whereas in reality at least some nucleosomes are missing, which would locally expand the system. Moreover, the full distributions of per-frame pairwise distances in simulation closely matches the per-cell distributions from ORCA (Figure 3e, Fig. S7). Our simulation protocol contains no free parameters controlling the relative values of specific pairwise distances, so the close correspondence with ORCA is validation of the simulation approach.

**Figure 3:**
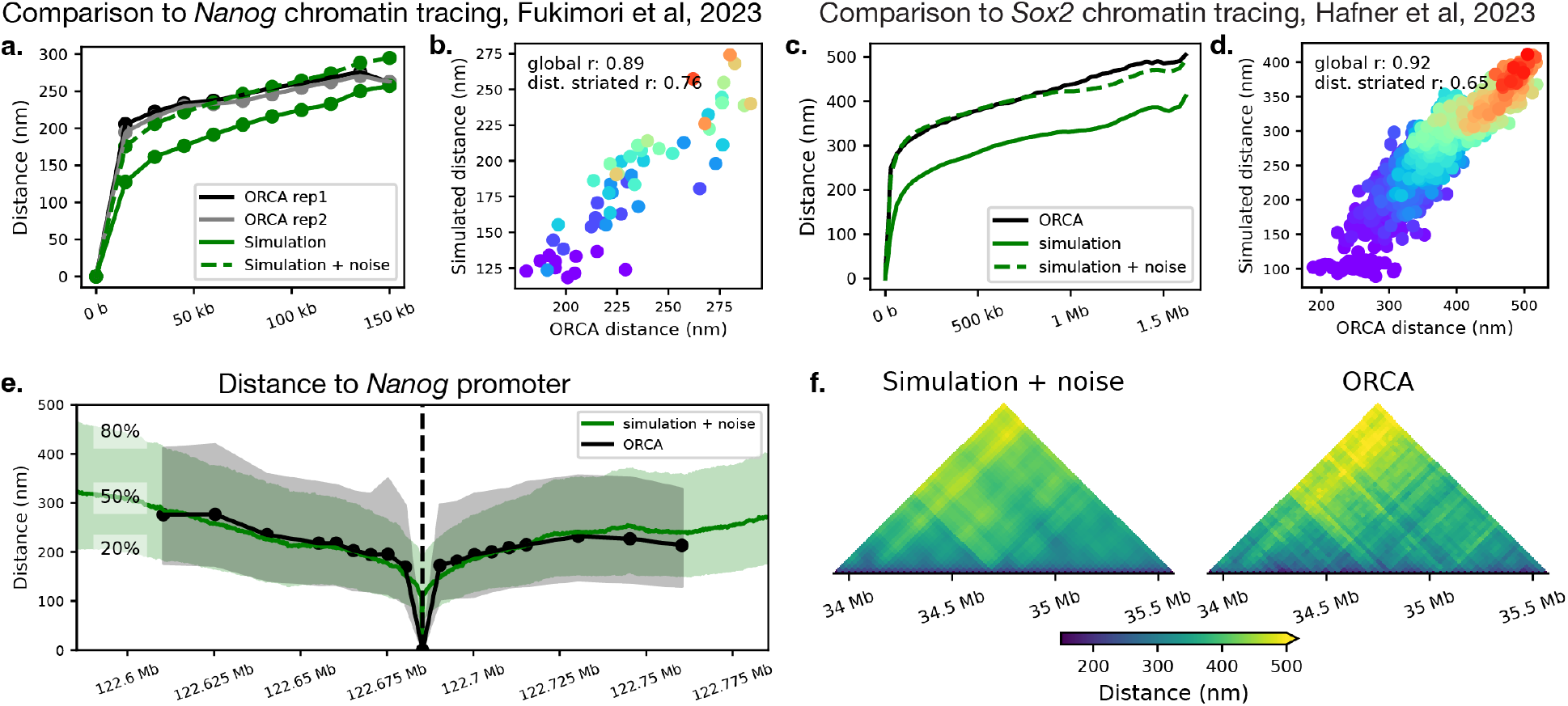
Simulations reproduce pairwise distance distributions from chromatin tracing. (a) Comparison of the median pairwise distances as a function of genomic separation from simulations of the *Nanog* locus to chromatin tracing data from Fujimori et al. [60]. The solid green line is the raw simulation output, and the dashed green line is the simulation output corrupted by measurement noise estimated using control experiments (see Supplementary Information). Importantly, the capture radius used in the simulations was selected to match this distance scaling trend, whereas the remaining results are validation. (b) Comparison of individual pairwise distances. Each point is colored by its genomic separation to highlight that the simulation predictions are better correlated with the chromatin tracing data than can be explained by only genomic separation. The distance striated correlation coefficient is the average of Pearson correlations computed separately for each genomic separation. (c, d) Equivalent to panels a and b but comparing simulations for the *Sox2* locus to chromatin tracing data from Hafner et al. [61]. (e) Distributions of distances to the *Nanog* promoter from simulation and chromatin tracing. The solid lines show the median value, while the shaded region encloses the 20 to 80th percentile. (f) Comparison of median distances from simulation and ORCA for the *Sox2* locus.

Together, these results show that our simulations recapitulate state-of-the-art experimental measurements of contacts frequencies from RCMC (Fig. S6) and pairwise distance distributions from ORCA (Figure 3, Fig. S7).

### Both heterochromatin and euchromatin are largely comprised of condensed domains

We ran simulations for five approximately 1 MB regions using RCMC data from Goel et al. [48]. The five regions mostly fall into the A-compartment based on eigenvector analysis of genome-wide micro-C data, except for *Fbn2*, which is mostly B-compartment (Figure 4a-c, Fig. S8). Visual analysis of our simulations reveals that the systems are comprised of condensed domains punctuated by kilobase-scale protrusions adopting extended structures (Figure 4d-g).

**Figure 4:**
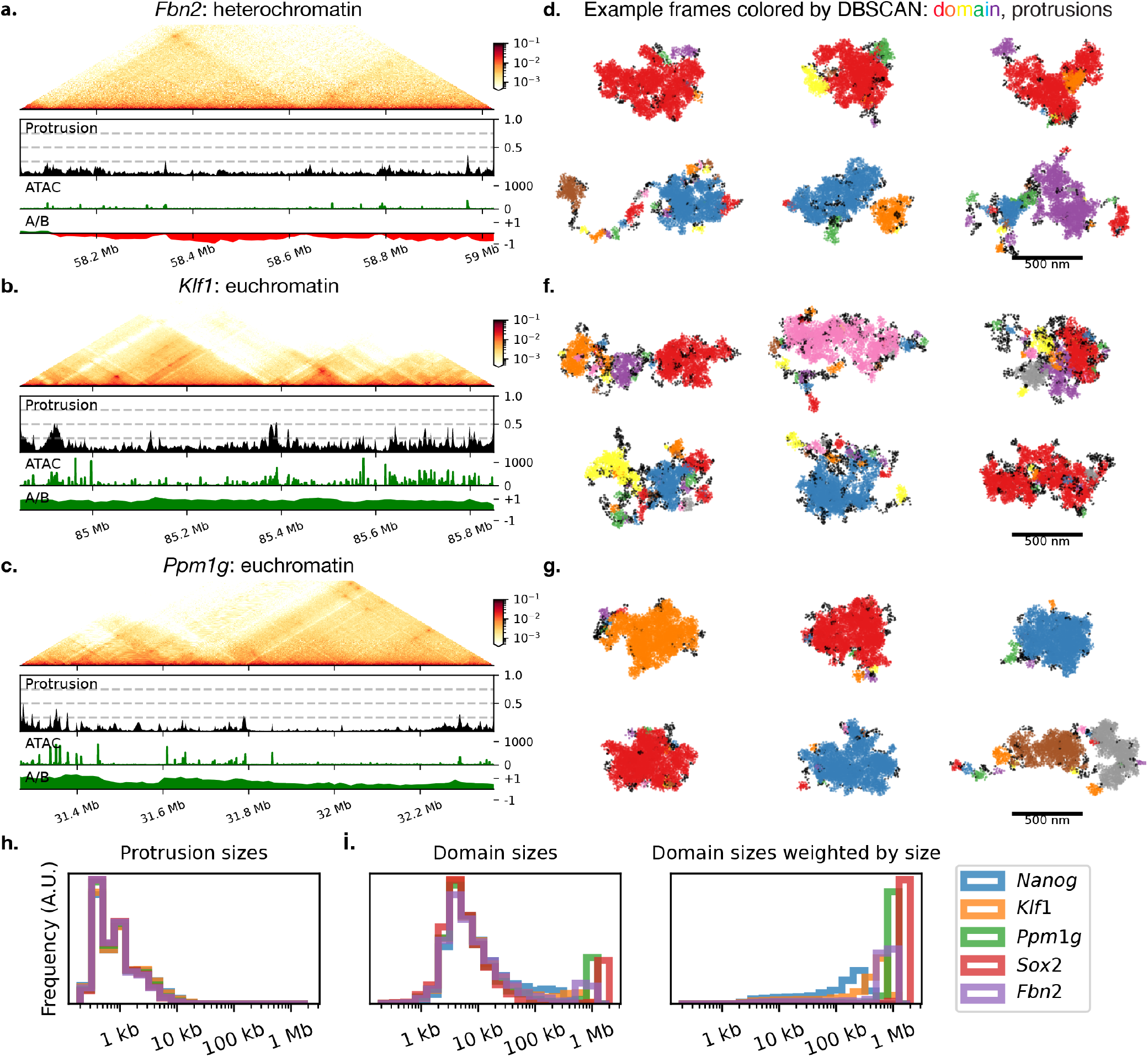
Both heterochromatin and euchromatin are largely comprised of condensed domains. (a, b, c) Contact maps from RCMC used in MEI, the probability of each position forming a protrusion, i.e. not in a domain, ATAC-seq coverage, and A/B compartment profile computed using genome-wide micro-C data [25]. (d, f, g) Randomly selected simulation frames for each of the three regions. Packing domains are shown in rainbow colors and protrusions are shown in black. (h) Protrusions are generally about a kilobase long. (i) Packing domains most commonly contain about 10 kb of genomic content (left), however, some packing domains are closer to 1 MB in size. Plotting packing domain sizes weighted by size (right) reveals that most nucleosomes are positioned in large packing domains.

To quantify these structures, we used density-based clustering with noise (DBSCAN) [62] (see Supplementary Information). DBSCAN groups densely packed points into clusters and assigns points in sparse regions as noise points. This leads to a natural interpretation of the clusters being “condensed domains” and the noise points being exposed “protrusions”. For both a heterochromatin region (*Fbn2*) and largely euchromatin regions (e.g. *Klf1* and *Ppm1g*), the majority of nucleosomes occur in domains with only *∼* 5 % of the *Fbn2, Sox2*, and *Ppm1g* regions and *∼* 15 % of the *Klf1* and *Nanog* regions forming protrusions (Table S1).

The domains identified by DBSCAN appear to correspond to experimentally observed packing (or TAD-like) domains [1, 12]. Consistent with experimental observations, the domains contain between a few kilobases to a megabase of DNA content and have diameters ranging from tens to hundreds of nanometers (Figure 4). Moreover, as has been observed in experiments [12, 13, 63] and previous simulation studies [64], the positioning of the domains in simulation varies greatly between single conformations. For example, the contact map for the *Fbn2* region suggests the presence of three topologically associating domains (TADs) (Fig. S10). In some frames, the entire region forms a single domain, while in others a single TAD is split into multiple packing domains. The packing domains are generally but not always formed by contiguous regions, as observed in chromatin tracing experiments [12].

The packing domains have nucleosome concentrations of approximately 300 µM (Fig S18). However, the precise value ranges depending on the approach used, ranging from about 250 µM using probe excluded volume to nearly 500 µM using the maximum local concentration (see Supplementary Information). The nucleosome concentration, as well as the scaling of radius of gyration with size (Figure 5c), shows only minor variations between regions, despite their different epigenetic states (Fig. S9). We will discuss this perhaps unexpected result in more detail later. This concentration is largely consistent with in vitro [65] and in vivo [4] measurements. Additionally, this packing domain density would allow the whole genome to fit comfortably in the nucleus with over half of the volume mostly devoid of chromatin [1]: 6 GB of DNA in a nucleus of 10 µm nucleus would require a nucleosome concentration of about 100 µM if uniformly spaced.

**Figure 5:**
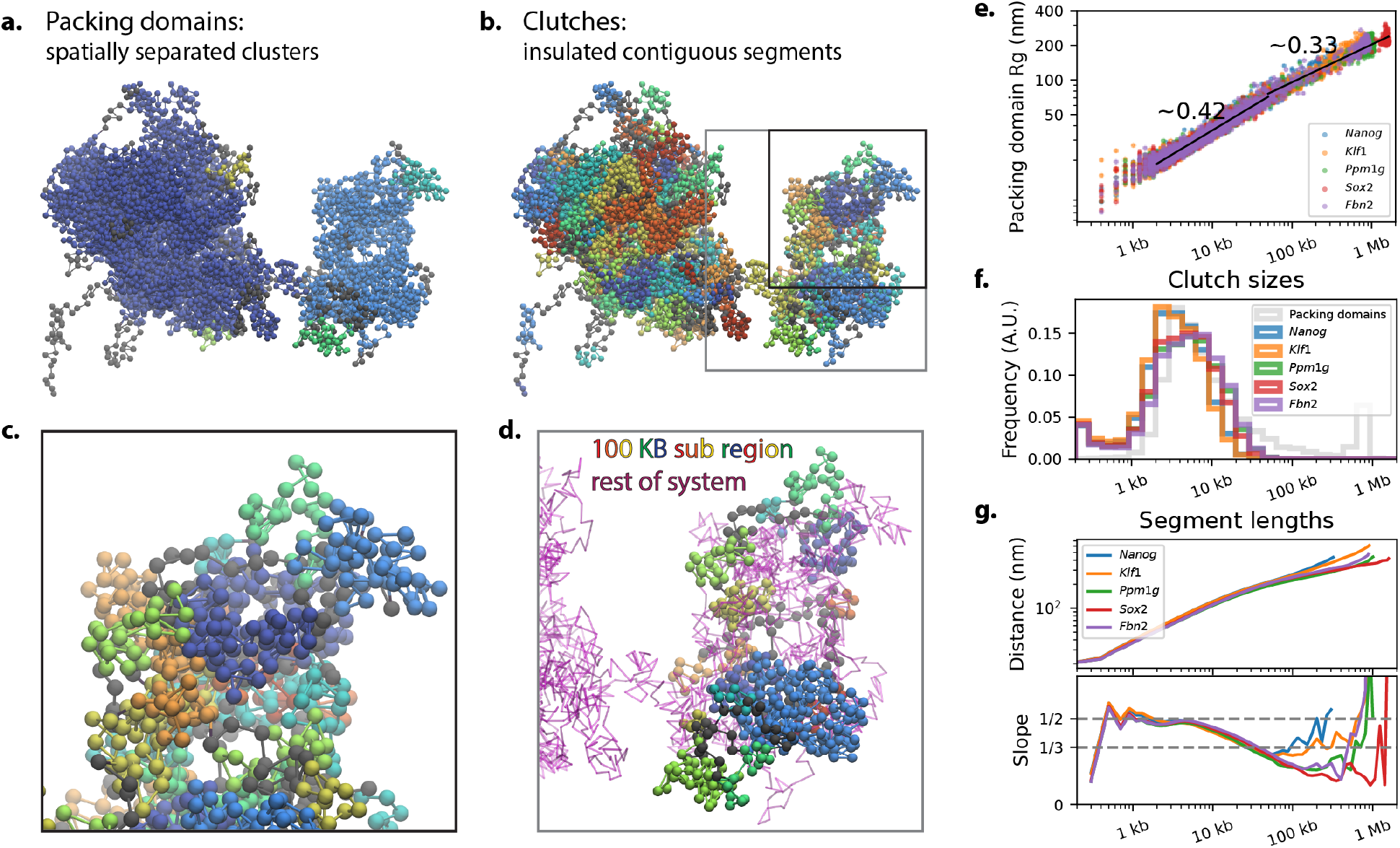
Packing domains are composed of clutches. (a) Illustration of the packing domains in a representative simulation frame. (b) Decomposition of the same simulation frame into clutches. (c) Zooming into the representative frame shows that clutches are generally unmixed, but contact each other on their surfaces to form packing domains. (d) A zoom-in highlighting the conformation of a 100 kb segment of the system. Modeling this region on its own would neglect a significant fraction of nucleosome interactions. (e) Relationship between the radius of gyration (Rg) and size of packing domains. Packing domains have similar Rg between regions, despite them having different epigenetic state. The scaling exponents are shown for domains smaller and larger than 50 KB. Small packing domains scale with a larger exponent because they are often formed by a single clutches and inherit their fiber-like character. (f) Clutches are about 10 kb in size, consistent with the peak in packing domain sizes. (g) Average segment lengths over each region. All regions initially display similar scaling, but at large separations gene rich regions, e.g. *Klf1*, are more extended.

### Packing domains are loose aggregations of clutches

Various imaging experiments suggest that packing domains contain internal structure. For instance, Szabo et al. [13] observed “chromatin nanodomains” corresponding to dense blobs inside of packing domains and Ricci et al. [5] observed “clutches” corresponding to clustered groups of tens of nucleosomes. Additionally, heterogeneous fiber-like structures have been observed in vitro [66] and in vivo using cryo-ET [7–9]. However, without the ability to trace the chromatin strand, it has proven hard to precisely characterize these structures and relate them to higher-order chromatin structure.

Indeed, packing domains in our simulations are far from homogeneous droplets. Especially for domains larger than 10 kb, packing domains contain dense regions interspersed with unfilled cavities (Figure 5a). Visual inspection shows that the dense regions largely correspond to contiguous chromatin segments (Figure 5b). This suggests that the chromatin chain decomposes into compact, contiguous segments, which then loosely aggregate to form packing domains. We refer to these contiguous segments as “contiguous nucleosome clutches” (or “clutches” for short) because they appear to correspond roughly to the small, dense structures observed in super-resolution images by Ricci et al. [5].

We define clutches using insulation scores calculated from single-frame contact maps. Positions with fewer than half the median number of cross contacts are defined as clutch boundaries, and the segments between such boundaries are defined as clutches (Supplementary Information). By construction clutches are insulated from flanking regions, however, we find that the clutches are also generally not mixed with other regions (Figure 5c, Fig. S14c). This definition differs from the original definition of nucleosome clutches as clustered nucleosome localizations in super-resolution images [5]. Under our definition, the clutches are explicitly defined as contiguous segments, which is enabled by the ability to trace the chromatin chain in our simulations. There are likely cases where multiple contiguous clutches are pressed against each other and thereby appear as a single cluster in imaging. Conversely, large or irregularly shaped contiguous clutches may be separated into multiple clusters or not fit in the field of view when imaged.

The precise choice of parameters used to define clutches, and thereby their precise sizes and boundary probabilities, is somewhat arbitrary, as hierarchically folded structure exists across a broad range of scales (Fig. S12). However, there are several factors supporting that clutches, as defined here, are an important layer of chromatin organization. First, these are the largest contiguous segments that are mostly unmixed (Fig. S14e). Second, their sizes coincide with a change in the end-to-end distance scaling exponent from 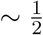 to 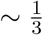 (Figure 5g). Finally, their size distribution overlaps the distribution of packing domain sizes, suggesting that clutches represent an underlying feature that can either occur alone as small packing domains or group together into large packing domains (Figure 5f, Figure 6a).

**Figure 6:**
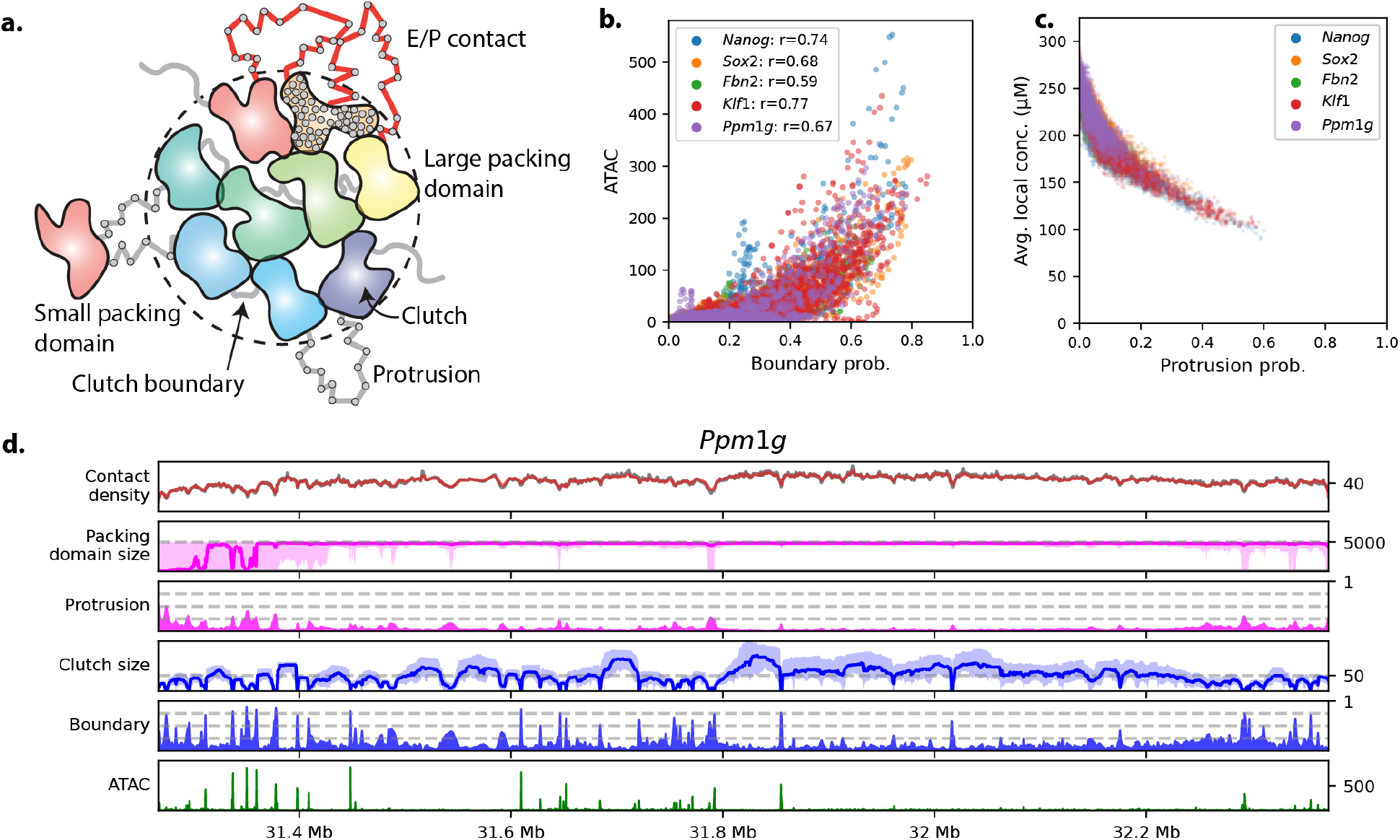
Protrusions, clutch boundaries, contact density, and ATAC-seq coverage are closely related. (a) Cartoon depicting the relationship between enhancers and promoters, clutches, and packing domains. (b) The probability of a boundary between clutches is correlated with ATAC-seq coverage. (c) The probability of a position occurring as a protrusion explains much of the variation in average local nucleosome concentration, defined as the concentration within a 40 nm sphere of each nucleosome. (d) Contact density, protrusion probability, clutch boundary probability, domain sizes, and ATAC-seq coverage for the *Ppm1g* region. For the domain size traces, the solid line shows the median value and the shaded region encloses the 25th to 75th percentile. The reference contact density is shown in red and the simulated contact frequency is shown in gray.

Clutches on average contain about 10 KB (50 nucleosomes) of genomic content (Figure 5d). This makes them similar in size to the experimentally observed clutches [5, 14]. The clutches are smaller in the vicinity of active regulatory elements and larger elsewhere (Figure 6d, Fig. S15a), consistent with experimentally observed changes in clutch size upon hyper-acetylation [67] and the preferential association of small clutches with nascent RNA [68]. Meanwhile, the chromatin nanodomains observed by Szabo et al. [13] are generally larger and likely represent aggregations of clutches. Clutches have an elongated fiber-like character with their radius of gyration scaling with genomic content with an exponent of 0.46 (Fig. S15b), reminiscent of the heterogeneous fibers observed in cryo-ET [7–9, 66]. In this way, packing domains can be viewed as a mosaic of fiber-like clutches (Figure 6a). This underlying structure is largely obscured without access to the chain index because clutches often press against each other on their surfaces, leading to continuously high density (Figure 5b,c).

Clutch boundaries are enriched at ATAC-seq peaks (Figure 6b, Fig. S13b). The most defined boundaries are present in over 75% of frames (Figure 6d). These well-phased boundaries result in clutches that appear in the population average contact map as small squares along the diagonal (Fig. S16) [25]. However, all positions, even those not associated with any clear features in the contact map, have a non-zero probability of forming a clutch boundary (Figure 6d, Fig. S13). These stochastic boundaries do not appear in the contact map because they are positioned differently between frames. In this way, there is a similar relationship between these small squares along the diagonal and clutches as between TADs and packing domains [12].

### Packing domains expose active enhancers and promoters at their periphery

We now turn our attention to the short “protrusions” from packing domains, which by definition are highly accessible. The protrusions we observe are typically on the scale of a few kilobases, rarely exceeding 10 kb, and generally adopt extended conformations, consistent with a 10 nm fiber (Figure 4h, Figure 5a). The protrusions sometimes link two packing domains, but just as frequently exit from and return to the same packing domain.

The probability of a nucleosome occurring in a protrusion is higher at active promoters and enhancers (Figure 6d, Fig. S15a,d). The highest protrusion probabilities, e.g. at 85.4 MB in the *Klf1* region, are *∼* 50% but probabilities of *∼* 25% are more typical. The probability of forming a protrusion is enriched specifically in short regions with active epigenetics marks, not the A compartment generally. For example, using the typical eigenvector analysis (at 12 kb resolution), the entirety of the simulated *Ppm1g* region is identified as A compartment. However, the region from 31.8–32.2 mb has few ATAC peaks and has a low probability of forming a protrusion (Figure 4b). Conversely, the *Fbn2* region is largely called as B compartment, but nonetheless there is an increase in protrusion probability at the few ATAC-seq peaks present (Figure 4a). Without neighbor balancing, we see little enrichment for protrusion formation at active promoters and enhancers (Fig. S11).

There is a close connection between protrusions and the boundaries between clutches (Figure 6a). Clutch boundaries are a local property and are formed when a few kilobases of chromatin adopt an extended conformation. Meanwhile, protrusions are a global property, and their formation requires a nucleosome not to be crowded by local interactions and to extend out from packing domains. In this way, a protrusion will generally be a clutch boundary, but a clutch boundary can be buried inside of a packing domain and thereby not a protrusion. The probabilities of a position forming a protrusion or clutch boundary are strongly correlated (Figure 6d, Fig. S15a,c), suggesting that local extension generally emerges as global accessibility independent of context. However, the correlation is not perfect, and it is possible that in some cases local and global compaction are modulated separately to control gene expression.

## Discussion

We introduced a protocol to generate nucleosome-resolution conformational ensembles that reproduce contact frequencies from RCMC and pairwise distance distributions from chromatin tracing. Our approach simultaneously provides high spatial and genomic resolution, allowing us to observe the connection between genomic features and structural motifs.

### Packing domains with regulatory protrusions

Our simulations suggest that euchromatin is generally comprised of condensed domains, but short regions at active promoters and enhancers sometimes form extended, highly exposed protrusions (Figure 6a). This view is consistent with a variety of experimental data suggesting that euchromatin is generally compact [2], while providing an explanation as to how enhancers and promoters are nonetheless accessible.

The tendency for active promoters and enhancers to form exposed protrusions leaves them accessible to transcription factors, the transcriptional machinery, and each other. Compared to a case where the entirety of euchromatin exists as a 10 nm fiber, the arrangement of the majority of euchromatin into domains lowers the entropic penalty for enhancer–promoter contacts by effectively shortening the tether between them. Moreover, non-active regions being less accessible limits the search space of transcription factors, potentially introduces a means to repress nearby genes, and allows compaction of the genome into the nucleus. Together, this arrangement naturally brings enhancers and promoters together, as they tend to be positioned on the exterior of packing domains, facilitating the formation of microcompartments [48] and fine-scale A-compartments [69].

### Packing domain density is independent of chromatin state

In our simulations, the density inside of packing domains is largely consistent between regions, despite differences in epigenetic state. This perhaps surprising result requires three clarifications. First, short regions near promoters and enhancers are more likely to exist as protrusions, which are substantially less dense and more extended. In this way, euchromatin defined narrowly as active promoters and enhancers is indeed on average less dense (Figure 6c,d). Second, gene-rich regions are broken into more, smaller packing domains (Figure 4i) and overall adopt more extended conformations on large length scales (Figure 5e). As a result, while the packing domains are of approximately the same density, the overall system is in some sense less dense [70, 71]. Moreover, since the surface-area-to-volume ratio of smaller domains is higher, these gene rich regions are generally more exposed. The fraction of time a nucleosome spends on the surface, which includes both time spent as a protrusion and time on the surface of a packing domain, explains much of the variation in local density (Fig. S9e). Third, here we only considered data for embryonic stem cells, which are believed to have an overall more open chromatin structure to support pluripotency [72]. Future work is needed to assess if heterochromatin regions in differentiated cells are more dense.

Several experimental observations support that most nucleosomes are compacted into domains with a similar density and only short regions around promoters and enhancers are highly exposed [2]. Imaging studies reveal that the chromatin is arranged heterogeneously with much of the genome grouped into dense domains and a small fraction partitioned into a substantially more dilute phase [1, 4, 73]. Tagging of specific regions [74] or histone marks [55] show that active regulatory elements are enriched in these dilute regions. Probing chromatin compaction by measuring sedimentation velocities [56] and digesting DNA with varying concentrations of MNase [57] both show that there are minimal large-scale differences in compaction but short regions around active promoters and enhancers are disproportionately open.

### Mechanisms of packing and clutch formation

Our simulations provide a unified account of sub-packing-domain-scale chromatin organization. Previous studies have reported dense nucleosome clusters inside of packing domains of different sizes [5, 13, 75]. Additionally, short sections of native chromatin have been observed to form heterogeneous fiber-like structures in vitro [66] and fiber-like structures have been observed in vivo [8, 9]. Our simulations suggest that contiguous sections of chromatin collapse along the chain to form clutches with heterogeneous fiber-like structures. These clutches aggregate hierarchically to form dense nucleosome clusters at a variety of scales, as well as packing domains themselves (Figure 5, Fig. S12). The boundaries between clutches, which are comprised of locally extended chromatin segments, often propagate up to emerge as protrusions, which are globally exposed. Different experimental techniques and data analysis pipelines likely produce different views of this underlying hierarchical structure.

The ability to trace the chromatin strand in our simulations allowed us to define clutches explicitly as contiguous chromatin segments. This is important because it highlights the mechanisms driving their formation and regulation. Clutches appear to be held together particularly tightly by the physical tethers between nucleosomes, corresponding to “correlation blobs” in polymer physics [76]. The boundaries between clutches are highly enriched at ATAC-seq peaks (Figure 6b). This likely stems from the presence of nucleosome-free regions at ATAC-seq peaks, which have been shown to induce domain boundaries in both experimental and computational studies [77–79]. Additionally, the boundaries are correlated with low condensability scores [54], suggesting that decompaction could be mediated by decreased nucleosome interaction energies (Figure 1c).

Simulations using RCMC data from cohesin-depleted cells [48] suggest that clutch formation is largely independent of loop extrusion. At large genomic separations, contact frequencies decrease, segment end-to-end distances increase, and the size of packing domains decreases substantially but they remain present (Fig. S17). This increase in segment end-to-end distances and the continued presence of packing domains is consistent with chromatin tracing data [12, 61]. Meanwhile, for separations less than 10 kb, there are only minor changes to contact frequencies, segment end-to-end distances, and clutch sizes. Cohesin depletion having only a minor change on 10 kb end-to-end distances is consistent with chromatin tracing data [80]. These findings suggest that, while cohesin-mediated loop extrusion plays a significant role in forming packing domains and large-scale structure generally, clutches are mostly shaped by other factors. This is also supported by the near 1/2 scaling of segment distances on short length scales (Figure 5g). Theoretical models of loop extrusion suggest this regime occurs because on short scales the chromatin chain has time to fully relax between cohesin looping events [81]. These models also predict the decay to scaling exponents less than 1/3 at above 30 kb observed in our simulations.

## Methods

### Neighbor balancing

The neighbor balanced contact frequencies *C*^(neighbor)^ are computed from the ICE balanced contact frequencies *C* as

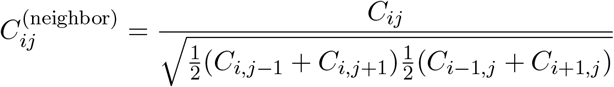

The ICE balanced contact frequencies are smoothed using a Gaussian filter with a standard deviation of 1 bin when computing the denominator of this expression. First applying ICE balancing and then applying neighbor balancing gives more robust results then applying neighbor balancing to the raw read counts.

This procedure is akin to the “square root vanilla coverage” balancing rule [82]. Note that if ICE balanced neighbor contacts were constant over a region, then this procedure would exactly update the neighbor contact frequencies to be one. In practice, the neighbor contacts are not constant and this procedure does not perfectly set the neighbor contact frequencies to one, but it does move them closer to one. While an analog of iterative correction could be used to bring the neighbor contact frequencies closer to 1, in practice, the simpler update rule is more robust.

We processed the RCMC data largely following the procedure from the original study. However, we found that several changes were necessary to facilitate using the data as nucleosome-level contact probabilities. These changes include: (1) using an estimated nucleosome center position, instead of the 5’ read end position when determining contact coordinates, (2) accounting for an overabundance of short range inward oriented junctions, which appear to be unligated artifacts, and (3) developing an augmented ICE balancing procedure that accounts for missing data due to low read coverage or low capture probe coverage. A complete description of the processing pipeline and motivation for modifications is available in the Supplementary Information.

We offer two guidelines for the practical application of neighbor balancing. First, applying neighbor balancing at a higher resolution than is supported by the sequencing depth results in noisy contact densities and should be avoided. As described in Supplementary Information, neighbor contact frequencies can be aggregated over larger windows and used to produce neighbor balanced contact maps at lower resolution. Second, contact densities should not be directly compared for micro-C datasets produced with different protocols. The micro-C procedure, especially the cross-linking profile, seems to influence the overall scale of the contact densities determined by neighbor balancing. In particular, stronger cross-linking yields lower contact densities, presumably mediated through a concomitant decrease in the capture radius. Consistent with this, Oksuz et al. [83] observed that using stronger crosslinking yields a higher fraction of short range cis contacts, consistent with a shorter capture radius.

### Maximum entropy inversion

Maximum entropy inversion is a strategy for parameterizing data-driven simulations. That is to determine the value of parameters in a system’s energy function that result in the system reproducing experimental observables. This is a good match for our problem, where the experimental data are the probabilities of each pair of nucleosomes being in contact. Maximum entropy inversion has been used previously to model chromatin conformations, but these studies have generally focused on modeling larger genomic regions at lower resolution [43, 84, 85].

Say we have experimental observations *T* = [*t*_1_, …, *t*_*n*_] where each observation corresponds to the average of some property of the system, e.g. *t*_*i*_ = *⟨f*_*i*_(*X*)*⟩* where *X* are the conformations of the system, *⟨·⟩* denotes the ensemble average, and *f*_*i*_ is a function mapping conformations of the system to a scalar. It can be shown that the least biased way to augment the system to *i*=1 match the experimental observations is to introduce auxiliary energy terms 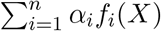. The energy coefficients *α* are derived using an iterative procedure wherein the system is simulated with the current coefficients, the observables are estimated from the simulated ensemble, and then the coefficients are updated by comparing the simulated and experimental values [43, 86].

In our work, the observables are the contact probabilities from micro-C, the functions *f*_*i*_ of interest are “contact indicator” functions modeling the probability of two nucleosomes forming a micro-C contact given their distance, and the coefficients *α*_*i*_ can be viewed as nucleosome-nucleosome interaction energies. Importantly, these interaction energies represent not just the intrinsic interaction energy between the two nucleosomes but also implicitly account for external factors, such as cohesin-mediated looping and interactions mediated by auxiliary proteins. In this way, the optimized system abstracts the molecular mechanisms into an information-theoretic energy landscape.

To initiate MEI, we must define a baseline model of the system and the contact indicator function, which we describe in detail below. For full details of the MEI procedure, see the Supplementary Information.

### Base nucleosome-level model

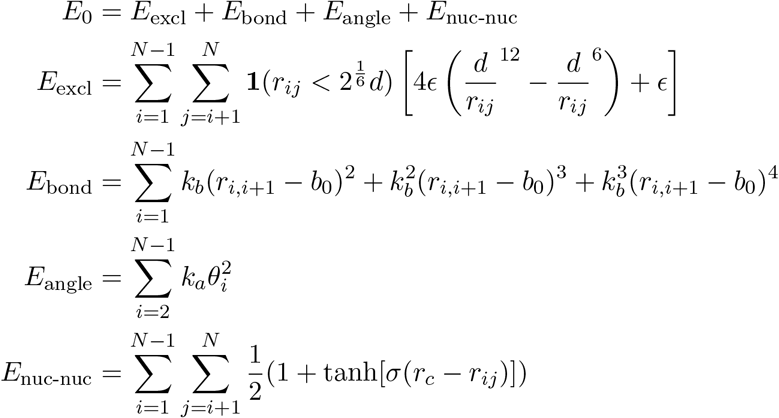

Nucleosomes are modeled as hard spheres with a diameter of *d* = 11 nm. This diameter was chosen to match the major axis of nucleosomes. We use the Lenard-Jones potential cut off at 2^1/6^*d* to only include the repulsive regime. We set *ϵ* = 1. Consecutive nucleosomes are connected by “class2” bonds with an equilibrium distance *b*_0_ = 21 nm fit to match the nucleosome-nucleosome distance observed in a cryo-EM structure of nucleosomes with a nucleosome repeat length of 197 bps plus 1 nm for the 3 additional base pairs [87]. The bond stiffness was set to 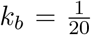, which allows fluctuations from approximately 10 to 30 nm, reflecting cases where neighboring nucleosomes are completely collapsed together to cases where nucleosomes are partially unraveled. The class2 potential was chosen to resist large deviations and favor collapse of nucleosomes together compared to large extensions.

The geometry of nucleosomes results in a preference for acute angles between three consecutive nucleosomes. It is hard to set the strength of this potential from first principles because the angle preference likely depends heavily on epigenetic modifications to the nucleosomes and the DNA-sequence. Therefore to set this parameter we used a top-down approach where we match the distribution of angles observed in cryo-ET structures of isolated chromatin [66]. We were not able to recapitulate the distribution exactly because our nucleosomes are too large to allow for very acute angles.

Nucleosomes form attractive interactions so we added short-range favorable nucleosome-nucleosome interactions [88, 89]. As for the angle energy coefficient, the nucleosome-nucleosome binding energy is perturbed by epigenetic modifications as well as salt concentrations, so it is hard to assign a priori. For the potential, we assumed the tanh functional form with *r*_*c*_ = 15 and *σ* = 8. We used an energy coefficient of -1 kT, which we justify below.

In preliminary simulations, nearly all of the structures contained knots. This is in conflict with experimental data suggesting that interphase chromatin is largely free of knots [43, 90], so we modified the system to prevent knotting. First, we introduced a string of spherical beads between each pair of bonded nucleosomes to prevent the strand from being able to cross itself. Second, we added a large bead to the end of each chain to prevent knots from entering the system through the ends. Buffer nucleosomes are added between these large beads and the core system to prevent this from biasing the ensemble. These additions leave the model kinetically trapped in an unknotted topology.

We sample from this model with Langevin dynamics using OpenMM [91]. Each monomer is assigned a mass of 1. The Langevin middle integrator [92] is used with a timestep of 0.125*τ*, a collision rate of 0.01*τ*^*−*^1, and a temperature of 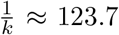 This choice of temperature results in *kT* being one, allowing us to easily specify our potential in reduced units. The simulations do not include periodic boundary conditions or any form of confinement potential.

While this is not the most accurate nucleosome-level model available, it captures the basic properties of chromatin and allows for simulations of large genomic regions.

### Nucleosome-interactions impact the optimized ensemble

Choosing the nucleosome interaction energy assumed by our base model from first principles is challenging because this quantity has proven hard to determine experimentally and depends on the conditions [88]. There is an argument that the interaction potential should be tuned to the theta condition. If we imagine the nucleus as a dense melt of chromatin, then we expect the chains to expand ideally as the interactions are effectively screened [76]. In this sense, tuning the interaction energy to the theta condition would implicitly account for the unmodeled remaining chromatin. However, imaging studies suggest the nucleus is not a dense melt of nucleosomes but separated into dense and dilute regions [1, 4]. This motivates the choice of an interaction energy stronger than that corresponding to the theta condition to implicitly account for the seemingly poor solvent.

To test the effects of this parameter, we performed MEI simulations on the *Nanog* region using base models with a range of interaction energies. We observed that stronger interactions yield systems that break into smaller, denser packing domains (Fig. S18b,f,h). However, this trend largely plateaus at or above the theta condition, which we found to be around 0.9 kT (Fig. S18a). In this regime, the conformations are visually similar (Fig. S18g) and protrusion probabilities show little change (Fig. S18i). In this way, our conclusions would not fundamentally change regardless of the specific value chosen. We selected an interaction energy of 1*kT* for our production simulations.

### Computing simulated contact maps

Given the output of a simulation, we wish to generate a simulated contact map comparable to contact maps from micro-C experiments. We assume the following model for how contact maps from micro-C arise. A large population of cells are provided as input to the micro-C experiment. In each cell, ligated junction reads are formed between nucleosomes (or other proteins) that are physically close to each other. We assume that the probability of a ligation is purely a function of the distance between nucleosomes, which we model using a “contact indicator” function *f*. The contact map is then produced by averaging the counts of junction reads across all the cells. This model leaves out details, such as that certain regions intrinsically produce more junction reads. We assume these complications are handled by ICE and neighbor balancing.

A simulation (or set thereof) gives frames *X* = *{X*^(*k*)^ *∈ R*^*m×*3^, *k* = 1..*N}* where *N* is the number of frames and *m* is the number of monomers. These simulation frames can be thought of as reflecting the chromatin conformations that might be present in a population of cells. Given a set of frames, we compute a simulated contact map *S ∈ R*^*m×m*^ by computing instantaneous contact maps *S*^(*k*)^ for each frame and then averaging them 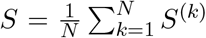 The instantaneous contact maps are computed as 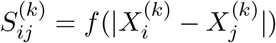

For the “contact indicator” function, which models the relative rate of observing a junction read between two nucleosomes as a function of distance, we use

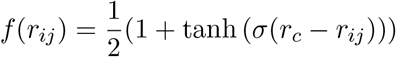

This function is one for short distances, reaches a value of 0.5 at the “capture radius” *r*_*c*_, and decays to zero for far distances. The parameter *σ* controls the rate of decay.

When comparing to micro-C contact maps, we scale these probabilities by a constant such that the average contact frequency between neighboring nucleosomes is matched between the simulated and reference contact maps.

The choice of contact indicator function has a major impact on the optimized ensemble. All else being equal, shorter capture radii result in dense, compact structures while longer capture radii result in dilute, extended conformations. This relationship is a complex function of the cross-linking, MNase digestion, and ligation protocol, and there does not appear to be a way to determine it from first principles.

We set the contact indicator function to reproduce the average pairwise distances determined by chromatin tracing data for the *Nanog* locus [60]. This required repeated optimizations with different capture radii. A capture radius of 42 nm (two bond lengths in our base model) provided a reasonable match (Figure 3a). We note that while the overall magnitude of average distances was used to determine the model and thereby cannot be considered validation. No free parameters were available to fit the shape of the distance scaling curves, the individual pairwise distances, nor the full distributions of individual pairwise distances. Close agreement of these three quantities therefore validates of our model.

## Acknowledgements

We are grateful to Advait Athreya, Anders Hansen, Alistair Boettiger, Lacramioara Bintu, and Taihei Fujimori for helpful discussions. This work was supported by the National Institutes of Health (Grant R35GM133580).

## Data and materials availability

The implementation for neighbor balancing is provided at: https://github.com/ZhangGroup-MITChemistry/neighbor-balance. Simulation trajectories and maximum entropy inversion code will be made available upon publication.

## References

[1] Ezequiel Miron, Roel Oldenkamp, Jill M. Brown, David M. S. Pinto, C. Shan Xu, Ana R. Faria, Haitham A. Shaban, James D. P. Rhodes, Cassandravictoria Innocent, Sara de Ornellas, Harald F. Hess, Veronica Buckle, and Lothar Schermelleh. Chromatin arranges in chains of mesoscale domains with nanoscale functional topography independent of cohesin. Science Advances, 6(39):eaba8811, September 2020. doi: 10.1126/sciadv.aba8811. URL https://www.science.org/doi/10.1126/sciadv.aba8811. Publisher: American Association for the Advancement of Science.

[2] Kazuhiro Maeshima, Shiori Iida, Masa A. Shimazoe, Sachiko Tamura, and Satoru Ide. Is euchromatin really open in the cell? Trends in Cell Biology, 34(1):7–17, January 2024. ISSN 1879-3088. doi: 10.1016/j.tcb.2023.05.007

[3] Tadasu Nozaki, Soya Shinkai, Satoru Ide, Koichi Higashi, Sachiko Tamura, Masa A. Shimazoe, Masaki Nakagawa, Yutaka Suzuki, Yasushi Okada, Masaki Sasai, Shuichi Onami, Ken Kurokawa, Shiori Iida, and Kazuhiro Maeshima. Condensed but liquid-like domain organization of active chromatin regions in living human cells. Science Advances, 9(14):eadf1488, April 2023. doi: 10.1126/sciadv.adf1488. URL https://www.science.org/doi/full/10.1126/sciadv.adf1488. Publisher: American Association for the Advancement of Science.

[4] Márton Gelléri, Shih-Ya Chen, Barbara Hubner, Jan Neumann, Ole Kroger, Filip Sadlo, Jorg Imhoff, Michael J. Hendzel, Marion Cremer, Thomas Cremer, Hilmar Strickfaden, and Christoph Cremer. True-to-scale DNA-density maps correlate with major accessibility differences between active and inactive chromatin. Cell Reports, 42(6):112567, June 2023. ISSN 2211-1247. doi: 10.1016/j.celrep.2023.112567. URL https://www.sciencedirect.com/science/article/pii/S2211124723005788.

[5] Maria Aurelia Ricci, Carlo Manzo, María Filomena García-Parajo, Melike Lakadamyali, and Maria Pia Cosma. Chromatin Fibers Are Formed by Heterogeneous Groups of Nucleosomes In Vivo. Cell, 160(6):1145–1158, March 2015. ISSN 0092-8674, 1097-4172. doi: 10.1016/j.cell.2015.01.054. URL https://www.cell.com/cell/abstract/S0092-8674(15)00132-4. Publisher: Elsevier.

[6] Horng D. Ou, Sébastien Phan, Thomas J. Deerinck, Andrea Thor, Mark H. Ellisman, and Clodagh C. O’Shea. ChromEMT: Visualizing 3D chromatin structure and compaction in interphase and mitotic cells. Science, 357(6349):eaag0025, July 2017. doi: 10.1126/science.aag0025

[7] Zhen Hou, Frank Nightingale, Yanan Zhu, Craig MacGregor-Chatwin, and Peijun Zhang. Structure of native chromatin fibres revealed by Cryo-ET in situ. Nature Communications, 14(1):6324, October 2023. ISSN 2041-1723. doi: 10.1038/s41467-023-42072-1. URL https://www.nature.com/articles/s41467-023-42072-1. Publisher: Nature Publishing Group.

[8] Yan Li, Haonan Zhang, Xiaomin Li, Wanyu Wu, and Ping Zhu. Cryo-ET study from in vitro to in vivo revealed a general folding mode of chromatin with two-start helical architecture. Cell Reports, 42(9):113134, September 2023. ISSN 2211-1247. doi: 10.1016/j.celrep.2023.113134. URL https://www.sciencedirect.com/science/article/pii/S2211124723011464.

[9] Jan Philipp Kreysing, Sergio Cruz-León, Johannes Betz, Carlotta Penzo, Tomáš Majtner, Markus Schreiber, Beata Turoňová, Marina Lusic, Gerhard Hummer, and Martin Beck. Molecular architecture of heterochromatin at the nuclear periphery of primary human cells, April 2025. URL https://www.biorxiv.org/content/10.1101/2025.04.09.647790v1. Pages: 2025.04.09.647790 Section: New Results.

[10] Jon Ken Chen, Tingsheng Liu, Shujun Cai, Weimei Ruan, Cai Tong Ng, Jian Shi, Uttam Surana, and Lu Gan. Nanoscale analysis of human G1 and metaphase chromatin in situ. EMBO J, 44(9):2658–2694, March 2025. doi: 10.1038/s44318-025-00407-2

[11] Wing Shun Li, Lucas M. Carter, Luay Matthew Almassalha, Ruyi Gong, Emily M. PujadasLiwag, Tiffany Kuo, Kyle L. MacQuarrie, Marcelo Carignano, Cody Dunton, Vinayak Dravid, Masato T. Kanemaki, Igal Szleifer, and Vadim Backman. Mature chromatin packing domains persist after RAD21 depletion in 3D. Science Advances, 11(4):eadp0855, January 2025. doi: 10.1126/sciadv.adp0855. URL https://www.science.org/doi/10.1126/sciadv.adp0855. Publisher: American Association for the Advancement of Science.

[12] Bogdan Bintu, Leslie J. Mateo, Jun-Han Su, Nicholas A. Sinnott-Armstrong, Mirae Parker, Seon Kinrot, Kei Yamaya, Alistair N. Boettiger, and Xiaowei Zhuang. Super-resolution chromatin tracing reveals domains and cooperative interactions in single cells. Science, 362(6413): eaau1783, October 2018. doi: 10.1126/science.aau1783. URL https://www.science.org/doi/10.1126/science.aau1783. Publisher: American Association for the Advancement of Science.

[13] Quentin Szabo, Axelle Donjon, Ivana Jerković, Giorgio L. Papadopoulos, Thierry Cheutin, Boyan Bonev, Elphege P. Nora, Benoit G. Bruneau, Frédéric Bantignies, and Giacomo Cavalli. Regulation of single-cell genome organization into TADs and chromatin nanodomains. Nature Genetics, 52(11):1151–1157, November 2020. ISSN 1546-1718. doi: 10.1038/s41588-020-00716-8. URL https://www.nature.com/articles/s41588-020-00716-8. Publisher: Nature Publishing Group.

[14] Melike Lakadamyali and Maria Pia Cosma. Visualizing the genome in high resolution challenges our textbook understanding. Nature Methods, 17(4):371–379, April 2020. ISSN 1548-7105. doi: 10.1038/s41592-020-0758-3. URL https://www.nature.com/articles/s41592-020-0758-3. Publisher: Nature Publishing Group.

[15] Brian J. Beliveau, Alistair N. Boettiger, Maier S. Avendaño, Ralf Jungmann, Ruth B. McCole, Eric F. Joyce, Caroline Kim-Kiselak, Frédéric Bantignies, Chamith Y. Fonseka, Jelena Erceg, Mohammed A. Hannan, Hien G. Hoang, David Colognori, Jeannie T. Lee, William M. Shih, Peng Yin, Xiaowei Zhuang, and Chao-ting Wu. Single-molecule super-resolution imaging of chromosomes and in situ haplotype visualization using Oligopaint FISH probes. Nature Communications, 6(1):7147, November 2015. doi: 10.1038/ncomms8147

[16] Yodai Takei, Yujing Yang, Jonathan White, Isabel N. Goronzy, Jina Yun, Meera Prasad, Lincoln J. Ombelets, Simone Schindler, Prashant Bhat, Mitchell Guttman, and Long Cai. Spatial multi-omics reveals cell-type-specific nuclear compartments. Nature, 641(8064):1037– 1047, May 2025. doi: 10.1038/s41586-025-08838-x

[17] Job Dekker, Karsten Rippe, Martijn Dekker, and Nancy Kleckner. chromosome conformation. Science (New York, N.Y.), 295(5558):1306–1311, February 2002. ISSN 1095-9203. doi: 10.1126/science.1067799

[18] Erez Lieberman-Aiden, Nynke L. Van Berkum, Louise Williams, Maxim Imakaev, Tobias Ragoczy, Agnes Telling, Ido Amit, Bryan R. Lajoie, Peter J. Sabo, Michael O. Dorschner, Richard Sandstrom, Bradley Bernstein, M. A. Bender, Mark Groudine, Andreas Gnirke, John Stamatoyannopoulos, Leonid A. Mirny, Eric S. Lander, and Job Dekker. Comprehensive mapping of long-range interactions reveals folding principles of the human genome. Science, 326(5950): 289–293, 2009. doi: 10.1126/science.1181369

[19] Longzhi Tan, Dong Xing, Chi-Han Chang, Heng Li, and X. Sunney Xie. Three-dimensional genome structures of single diploid human cells. Science, 361(6405): 924–928, 2018. doi: 10.1126/science.aat5641

[20] Robert A. Beagrie, Antonio Scialdone, Markus Schueler, Dorothee C.A. Kraemer, Mita Chotalia, Sheila Q. Xie, Mariano Barbieri, Inês De Santiago, Liron Mark Lavitas, Miguel R. Branco, James Fraser, Josée Dostie, Laurence Game, Niall Dillon, Paul A.W. Edwards, Mario Nicodemi, and Ana Pombo. Complex multi-enhancer contacts captured by genome architecture mapping. Nature, 543(7646): 519–524, 2017. doi: 10.1038/nature21411

[21] Sofia A. Quinodoz, Noah Ollikainen, Barbara Tabak, Ali Palla, Jan Marten Schmidt, Elizabeth Detmar, Mason M. Lai, Alexander A. Shishkin, Prashant Bhat, Yodai Takei, Vickie Trinh, Erik Aznauryan, Pamela Russell, Christine Cheng, Marko Jovanovic, Amy Chow, Long Cai, Patrick McDonel, Manuel Garber, and Mitchell Guttman. Higher-Order Inter-chromosomal Hubs Shape 3D Genome Organization in the Nucleus. Cell, 174(3):744–757.e24, 2018. doi: 10.1016/j.cell.2018.05.024

[22] Tsung-Han S. Hsieh, Assaf Weiner, Bryan Lajoie, Job Dekker, Nir Friedman, and Oliver J. Rando. Mapping Nucleosome Resolution Chromosome Folding in Yeast by Micro-C. Cell, 162(1):108–119, July 2015. ISSN 0092-8674. doi: 10.1016/j.cell.2015.05.048. URL https://www.sciencedirect.com/science/article/pii/S0092867415006388.

[23] Masae Ohno, Tadashi Ando, David G. Priest, Vipin Kumar, Yamato Yoshida, and Yuichi Taniguchi. Sub-nucleosomal Genome Structure Reveals Distinct Nucleosome Folding Motifs. Cell, 176(3):520–534.e25, January 2019. doi: 10.1016/j.cell.2018.12.014

[24] Nils Krietenstein, Sameer Abraham, Sergey V. Venev, Nezar Abdennur, Johan Gibcus, Tsung-han S. Hsieh, Krishna Mohan Parsi, Liyan Yang, René Maehr, Leonid A. Mirny, Job Dekker, and Oliver J. Rando. Ultrastructural Details of Mammalian Chromosome Architecture. Molecular Cell, 78(3):554–565.e7, May 2020. doi: 10.1016/j.molcel.2020.03.003

[25] Tsung-Han S. Hsieh, Claudia Cattoglio, Elena Slobodyanyuk, Anders S. Hansen, Oliver J. Rando, Robert Tjian, and Xavier Darzacq. Resolving the 3D Landscape of Transcription-Linked Mammalian Chromatin Folding. Molecular Cell, 78(3):539–553.e8, May 2020. ISSN 1097-2765. doi: 10.1016/j.molcel.2020.03.002. URL https://www.cell.com/molecular-cell/abstract/S1097-2765(20)30150-7. Publisher: Elsevier.

[26] Guang Shi and D. Thirumalai. From Hi-C Contact Map to Three-Dimensional Organization of Interphase Human Chromosomes. Physical Review X, 11(1): 11051, 2021. doi: 10.1103/PhysRevX.11.011051

[27] Sangram Kadam, Kiran Kumari, Vinoth Manivannan, Shuvadip Dutta, Mithun K. Mitra, and Ranjith Padinhateeri. Predicting scale-dependent chromatin polymer properties from systematic coarse-graining. Nature Communications, 14(1):4108, July 2023. doi: 10.1038/s41467-023-39907-2

[28] Lorenzo Boninsegna, Asli Yildirim, Guido Polles, Yuxiang Zhan, Sofia A Quinodoz, Elizabeth H Finn, Mitchell Guttman, Xianghong Jasmine Zhou, and Frank Alber. Integrative genome modeling platform reveals essentiality of rare contact events in 3D genome organizations. Nature Methods, 19(8):938–949, August 2022. doi: 10.1038/s41592-022-01527-x

[29] Johannes Nuebler, Geoffrey Fudenberg, Maxim Imakaev, Nezar Abdennur, and Leonid A. Mirny. Chromatin organization by an interplay of loop extrusion and compartmental segregation. Proceedings of the National Academy of Sciences, 115(29):E6697–E6706, July 2018. doi: 10.1073/pnas.1717730115

[30] M. Di Pierro, B. Zhang, E.L. L Aiden, P.G. G Wolynes, and J.N. N Onuchic. Transferable model for chromosome architecture. Proceedings of the National Academy of Sciences, 113 (43):12168–12173, 2016. doi: 10.1073/pnas.1613607113

[31] Michele Di Pierro, Ryan R. Cheng, Erez Lieberman Aiden, Peter G. Wolynes, and José N. Onuchic. De novo prediction of human chromosome structures: Epigenetic marking patterns encode genome architecture. Proceedings of the National Academy of Sciences, 114(46):12126–12131, 2017. doi: 10.1073/pnas.1714980114

[32] Michael Chiang, Chris A. Brackley, Catherine Naughton, Ryu-Suke Nozawa, Cleis Battaglia, Davide Marenduzzo, and Nick Gilbert. Genome-wide chromosome architecture prediction reveals biophysical principles underlying gene structure. Cell Genomics, 4(12):100698, December 2024. doi: 10.1016/j.xgen.2024.100698

[33] Hossein Salari, Genevieve Fourel, and Daniel Jost. Transcription regulates the spatio-temporal dynamics of genes through micro-compartmentalization. Nat Commun, 15(1):5393, June 2024. doi: 10.1038/s41467-024-49727-7

[34] Luca Fiorillo, Francesco Musella, Mattia Conte, Rieke Kempfer, Andrea M. Chiariello, Simona Bianco, Alexander Kukalev, Ibai Irastorza-Azcarate, Andrea Esposito, Alex Abraham, Antonella Prisco, Ana Pombo, and Mario Nicodemi. Comparison of the Hi-C, GAM and SPRITE methods using polymer models of chromatin. Nature Methods, 18(5): 482–490, 2021. doi: 10.1038/s41592-021-01135-1

[35] Maria Victoria Neguembor, Juan Pablo Arcon, Diana Buitrago, Rafael Lema, Jurgen Walther, Ximena Garate, Laura Martin, Pablo Romero, Jumana AlHaj Abed, Marta Gut, Julie Blanc, Melike Lakadamyali, Chao-ting Wu, Isabelle Brun Heath, Modesto Orozco, Pablo D. Dans, and Maria Pia Cosma. MiOS, an integrated imaging and computational strategy to model gene folding with nucleosome resolution. Nat Struct Mol Biol, 29(10):1011–1023, October 2022. doi: 10.1038/s41594-022-00839-y

[36] Joseph G. Wakim and Andrew J. Spakowitz. Physical modeling of nucleosome clustering in euchromatin resulting from interactions between epigenetic reader proteins. Proc. Natl. Acad. Sci. U.S.A., 121(26), June 2024. doi: 10.1073/pnas.2317911121

[37] Sumitabha Brahmachari, Vinícius G Contessoto, Michele Di Pierro, and José N Onuchic. Shaping the genome via lengthwise compaction, phase separation, and lamina adhesion. Nucleic Acids Research, 50(8):1–14, May 2022. doi: 10.1093/nar/gkac231

[38] Andriy Goychuk, Deepti Kannan, Arup K. Chakraborty, and Mehran Kardar. Polymer folding through active processes recreates features of genome organization. Proceedings of the National Academy of Sciences of the United States of America, 120(20): 1–12, 2023. doi: 10.1073/pnas.2221726120

[39] Xiakun Chu and Jin Wang. Deciphering the molecular mechanism of the cancer formation by chromosome structural dynamics. PLOS Computational Biology, 17(11):e1009596, November 2021. doi: 10.1371/journal.pcbi.1009596

[40] Stephanie Portillo-Ledesma, Zilong Li, and Tamar Schlick. Genome modeling: From chromatin fibers to genes. Current Opinion in Structural Biology, 78:102506, February 2023. doi: 10.1016/j.sbi.2022.102506

[41] Giovanni B Brandani, Chenyang Gu, Soundhararajan Gopi, and Shoji Takada. Multiscale Bayesian simulations reveal functional chromatin condensation of gene loci. PNAS Nexus, 3 (6), May 2024. doi: 10.1093/pnasnexus/pgae226

[42] Eric R. Schultz, Soren Kyhl, Rebecca Willett, and Juan J. de Pablo. Chromatin structures from integrated AI and polymer physics model. PLoS Comput Biol, 21(4):e1012912, April 2025. doi: 10.1371/journal.pcbi.1012912

[43] Bin Zhang and Peter G. Wolynes. Topology, structures, and energy landscapes of human chromosomes. Proceedings of the National Academy of Sciences, 112(19):6062–6067, May 2015. doi: 10.1073/pnas.1506257112. URL https://www.pnas.org/doi/10.1073/pnas.1506257112. Publisher: Proceedings of the National Academy of Sciences.

[44] Xingcheng Lin, Yifeng Qi, Andrew P. Latham, and Bin Zhang. Multiscale modeling of genome organization with maximum entropy optimization. The Journal of Chemical Physics, 155(1): 010901, July 2021. ISSN 0021-9606. doi: 10.1063/5.0044150. URL https://doi.org/10.1063/5.0044150.

[45] Sucheol Shin, Guang Shi, and D. Thirumalai. From Effective Interactions Extracted Using Hi-C Data to Chromosome Structures in Conventional and Inverted Nuclei. PRX Life, 1(1), August 2023. doi: 10.1103/prxlife.1.013010

[46] Greg Schuette, Xinqiang Ding, and Bin Zhang. Efficient Hi-C inversion facilitates chromatin folding mechanism discovery and structure prediction. Biophysical Journal, 122(17):3425–3438, September 2023. doi: 10.1016/j.bpj.2023.07.017

[47] Wen Jun Xie and Bin Zhang. Learning the Formation Mechanism of Domain-Level Chromatin States with Epigenomics Data. Biophysical Journal, 116(10):2047–2056, May 2019. doi: 10.1016/j.bpj.2019.04.006

[48] Viraat Y. Goel, Miles K. Huseyin, and Anders S. Hansen. Region Capture Micro-C reveals coalescence of enhancers and promoters into nested microcompartments. Nature Genetics, 55 (6):1048–1056, June 2023. ISSN 1061-4036, 1546-1718. doi: 10.1038/s41588-023-01391-1. URL https://www.nature.com/articles/s41588-023-01391-1.

[49] Clarice K. Y. Hong, Fan Feng, Varshini Ramanathan, Jie Liu, and Anders S. Hansen. Genome structure mapping with high-resolution 3D genomics and deep learning, May 2025. URL https://www.biorxiv.org/content/10.1101/2025.05.06.650874v1. Pages: 2025.05.06.650874 Section: New Results.

[50] Viraat Y. Goel, Nicholas G. Aboreden, James M. Jusuf, Haoyue Zhang, Luisa P. Mori, Leonid A. Mirny, Gerd A. Blobel, Edward J. Banigan, and Anders S. Hansen. Dynamics of microcompartment formation at the mitosis-to-G1 transition, September 2024. URL https://www.biorxiv.org/content/10.1101/2024.09.16.611917v1. Pages: 2024.09.16.611917 Section: New Results.

[51] Maxim Imakaev, Geoffrey Fudenberg, Rachel Patton McCord, Natalia Naumova, Anton Goloborodko, Bryan R. Lajoie, Job Dekker, and Leonid A. Mirny. Iterative correction of Hi-C data reveals hallmarks of chromosome organization. Nature Methods, 9(10):999–1003, October 2012. ISSN 1548-7105. doi: 10.1038/nmeth.2148. URL https://www.nature.com/articles/nmeth.2148. Publisher: Nature Publishing Group.

[52] Philip A. Knight and Daniel Ruiz. A fast algorithm for matrix balancing. IMA Journal of Numerical Analysis, 33(3):1029–1047, July 2013. ISSN 0272-4979. doi: 10.1093/imanum/drs019. URL https://doi.org/10.1093/imanum/drs019.

[53] Jin H. Yang and Anders S. Hansen. Enhancer selectivity in space and time: from enhancer–promoter interactions to promoter activation. Nature Reviews Molecular Cell Biology, 25(7):574–591, July 2024. ISSN 1471-0080. doi: 10.1038/s41580-024-00710-6. URL https://www.nature.com/articles/s41580-024-00710-6. Publisher: Nature Publishing Group.

[54] Sangwoo Park, Raquel Merino-Urteaga, Violetta Karwacki-Neisius, Gustavo Ezequiel Carrizo, Advait Athreya, Alberto Marin-Gonzalez, Nils A. Benning, Jonghan Park, Michelle M. Mitchener, Natarajan V. Bhanu, Benjamin A. Garcia, Bin Zhang, Tom W. Muir, Erika L. Pearce, and Taekjip Ha. Native nucleosomes intrinsically encode genome organization principles. Nature, 643(8071):572–581, July 2025. ISSN 1476-4687. doi: 10.1038/s41586-025-08971-7. URL https://www.nature.com/articles/s41586-025-08971-7. Publisher: Nature Publishing Group.

[55] Jianquan Xu, Hongqiang Ma, Jingyi Jin, Shikhar Uttam, Rao Fu, Yi Huang, and Yang Liu. Super-Resolution Imaging of Higher-Order Chromatin Structures at Different Epigenomic States in Single Mammalian Cells. Cell Reports, 24(4):873–882, July 2018. ISSN 2211-1247. doi: 10.1016/j.celrep.2018.06.085. URL https://www.sciencedirect.com/science/article/pii/S221112471831012X.

[56] Satoru Ishihara, Yohei Sasagawa, Takeru Kameda, Hayato Yamashita, Mana Umeda, Naoe Kotomura, Masayuki Abe, Yohei Shimono, and Itoshi Nikaido. Local states of chromatin compaction at transcription start sites control transcription levels. Nucleic Acids Research, 49(14):8007–8023, August 2021. ISSN 0305-1048. doi: 10.1093/nar/gkab587. URL https://doi.org/10.1093/nar/gkab587.

[57] Uwe Schwartz, Attila Németh, Sarah Diermeier, Josef H Exler, Stefan Hansch, Rodrigo Maldonado, Leonhard Heizinger, Rainer Merkl, and Gernot Langst. Characterizing the nuclease accessibility of DNA in human cells to map higher order structures of chromatin. Nucleic Acids Research, 47(3):1239–1254, February 2019. ISSN 0305-1048. doi: 10.1093/nar/gky1203. URL https://doi.org/10.1093/nar/gky1203.

[58] Yohsuke T. Fukai, Tomoya Kujirai, Masatoshi Wakamori, Setsuko Kanamura, Lisa Yamauchi, Somayeh Zeraati, Chiharu Tanegashima, Mitsutaka Kadota, Hitoshi Kurumizaka, Takashi Umehara, and Kyogo Kawaguchi. Engineered acetylation patterns drive large-scale chromatin organization in vitro, November 2024. URL https://www.biorxiv.org/content/10.1101/2024.11.08.622658v1. Pages: 2024.11.08.622658 Section: New Results.

[59] Martin Falk, Yana Feodorova, Natalia Naumova, Maxim Imakaev, Bryan R. Lajoie, Heinrich Leonhardt, Boris Joffe, Job Dekker, Geoffrey Fudenberg, Irina Solovei, and Leonid A. Mirny. Heterochromatin drives compartmentalization of inverted and conventional nuclei. Nature, 570 (7761):395–399, June 2019. ISSN 1476-4687. doi: 10.1038/s41586-019-1275-3. URL https://www.nature.com/articles/s41586-019-1275-3. Publisher: Nature Publishing Group.

[60] Taihei Fujimori, Carolina Rios-Martinez, Abby R. Thurm, Michaela M. Hinks, Benjamin R. Doughty, Joydeb Sinha, Derek Le, Antonina Hafner, William J. Greenleaf, Alistair N. Boettiger, and Lacramioara Bintu. Single-cell chromatin state transitions during epigenetic memory formation. bioRxiv, page 2023.10.03.560616, October 2023. doi: 10.1101/2023.10.03.560616. URL https://www.ncbi.nlm.nih.gov/pmc/articles/PMC10592931/.

[61] Antonina Hafner, Minhee Park, Scott E. Berger, Sedona E. Murphy, Elphege P. Nora, and Alistair N. Boettiger. Loop stacking organizes genome folding from TADs to chromosomes. Molecular Cell, 83(9):1377–1392.e6, May 2023. ISSN 1097-2765. doi: 10.1016/j.molcel.2023.04.008. URL https://www.sciencedirect.com/science/article/pii/S1097276523002526.

[62] Martin Ester, Hans-Peter Kriegel, Jorg Sander, and Xiaowei Xu. A density-based algorithm for discovering clusters in large spatial databases with noise. In Proceedings of the Second International Conference on Knowledge Discovery and Data Mining, KDD’96, pages 226–231, Portland, Oregon, August 1996. AAAI Press.

[63] Jennifer M. Luppino, Daniel S. Park, Son C. Nguyen, Yemin Lan, Zhuxuan Xu, Rebecca Yunker, and Eric F. Joyce. Cohesin promotes stochastic domain intermingling to ensure proper regulation of boundary-proximal genes. Nature Genetics, 52(8):840–848, August 2020. ISSN 1546-1718. doi: 10.1038/s41588-020-0647-9. URL https://www.nature.com/articles/s41588-020-0647-9. Publisher: Nature Publishing Group.

[64] Mattia Conte, Luca Fiorillo, Simona Bianco, Andrea M. Chiariello, Andrea Esposito, and Mario Nicodemi. Polymer physics indicates chromatin folding variability across single-cells results from state degeneracy in phase separation. Nature Communications, 11(1):3289, July 2020. ISSN 2041-1723. doi: 10.1038/s41467-020-17141-4. URL https://www.nature.com/articles/s41467-020-17141-4. Publisher: Nature Publishing Group.

[65] Meng Zhang, César Díaz-Celis, Bibiana Onoa, Cristhian Cañari-Chumpitaz, Katherinne I. Requejo, Jianfang Liu, Michael Vien, Eva Nogales, Gang Ren, and Carlos Bustamante. Molecular organization of the early stages of nucleosome phase separation visualized by cryo-electron tomography. Molecular Cell, 82(16):3000–3014.e9, August 2022. ISSN 1097-2765. doi: 10.1016/j.molcel.2022.06.032. URL https://www.sciencedirect.com/science/article/pii/S1097276522006505.

[66] Nathan Jentink, Carson Purnell, Brianna Kable, Matthew T. Swulius, and Sergei A. Grigoryev. Cryoelectron tomography reveals the multiplex anatomy of condensed native chromatin and its unfolding by histone citrullination. Molecular Cell, 83(18):3236–3252.e7, September 2023. ISSN 1097-2765. doi: 10.1016/j.molcel.2023.08.017. URL https://www.sciencedirect.com/science/article/pii/S1097276523006536.

[67] Jason Otterstrom, Alvaro Castells-Garcia, Chiara Vicario, Pablo A Gomez-Garcia, Maria Pia Cosma, and Melike Lakadamyali. Super-resolution microscopy reveals how histone tail acetylation affects DNA compaction within nucleosomes in vivo. Nucleic Acids Research, 47 (16):8470–8484, September 2019. ISSN 0305-1048. doi: 10.1093/nar/gkz593. URL https://doi.org/10.1093/nar/gkz593.

[68] Alvaro Castells-Garcia, Ilyas Ed-daoui, Esther González-Almela, Chiara Vicario, Jason Otte-strom, Melike Lakadamyali, Maria Victoria Neguembor, and Maria Pia Cosma. Super resolution microscopy reveals how elongating RNA polymerase II and nascent RNA interact with nucleosome clutches. Nucleic Acids Research, 50(1):175–190, January 2022. ISSN 0305-1048. doi: 10.1093/nar/gkab1215. URL https://doi.org/10.1093/nar/gkab1215.

[69] Hannah L. Harris, Huiya Gu, Moshe Olshansky, Ailun Wang, Irene Farabella, Yossi Eliaz, Achyuth Kalluchi, Akshay Krishna, Mozes Jacobs, Gesine Cauer, Melanie Pham, Suhas S. P. Rao, Olga Dudchenko, Arina Omer, Kiana Mohajeri, Sungjae Kim, Michael H. Nichols, Eric S. Davis, Dimos Gkountaroulis, Devika Udupa, Aviva Presser Aiden, Victor G. Corces, Douglas H. Phanstiel, William Stafford Noble, Guy Nir, Michele Di Pierro, Jeong-Sun Seo, Michael E. Talkowski, Erez Lieberman Aiden, and M. Jordan Rowley. Chromatin alternates between A and B compartments at kilobase scale for subgenic organization. Nature Communications, 14(1):3303, June 2023. ISSN 2041-1723. doi: 10.1038/s41467-023-38429-1. URL https://www.nature.com/articles/s41467-023-38429-1. Publisher: Nature Publishing Group.

[70] Sedona Eve Murphy and Alistair Nicol Boettiger. Polycomb repression of Hox genes involves spatial feedback but not domain compaction or phase transition. Nature Genetics, 56(3):493–504, March 2024. ISSN 1546-1718. doi: 10.1038/s41588-024-01661-6. URL https://www.nature.com/articles/s41588-024-01661-6. Publisher: Nature Publishing Group.

[71] Yodai Takei, Jina Yun, Shiwei Zheng, Noah Ollikainen, Nico Pierson, Jonathan White, Sheel Shah, Julian Thomassie, Shengbao Suo, Chee-Huat Linus Eng, Mitchell Guttman, Guo-Cheng Yuan, and Long Cai. Integrated spatial genomics reveals global architecture of single nuclei. Nature, 590(7845):344–350, February 2021. ISSN 1476-4687. doi: 10.1038/s41586-020-03126-2 URL https://www.nature.com/articles/s41586-020-03126-2. Number: 7845 Publisher: Nature Publishing Group.

[72] Alexandre Gaspar-Maia, Adi Alajem, Eran Meshorer, and Miguel Ramalho-Santos. Open chromatin in pluripotency and reprogramming. Nature Reviews Molecular Cell Biology, 12(1): 36–47, January 2011. ISSN 1471-0080. doi: 10.1038/nrm3036. URL https://www.nature.com/articles/nrm3036. Publisher: Nature Publishing Group.

[73] Timothy A. Daugird, Yu Shi, Katie L. Holland, Hosein Rostamian, Zhe Liu, Luke D. Lavis, Joseph Rodriguez, Brian D. Strahl, and Wesley R. Legant. Correlative single molecule lattice light sheet imaging reveals the dynamic relationship between nucleosomes and the local chromatin environment. Nature Communications, 15(1):4178, May 2024. ISSN 2041-1723. doi: 10.1038/s41467-024-48562-0. URL https://www.nature.com/articles/s41467-024-48562-0. Publisher: Nature Publishing Group.

[74] Marion Cremer, Volker J. Schmid, Felix Kraus, Yolanda Markaki, Ines Hellmann, Andreas Maiser, Heinrich Leonhardt, Sam John, John Stamatoyannopoulos, and Thomas Cremer. Initial high-resolution microscopic mapping of active and inactive regulatory sequences proves non-random 3D arrangements in chromatin domain clusters. Epigenetics & Chromatin, 10(1):39, August 2017. ISSN 1756-8935. doi: 10.1186/s13072-017-0146-0. URL https://doi.org/10.1186/s13072-017-0146-0.

[75] Yue Li, Vasundhara Agrawal, Ranya K. A. Virk, Eric Roth, Wing Shun Li, Adam Eshein, Jane Frederick, Kai Huang, Luay Almassalha, Reiner Bleher, Marcelo A. Carignano, Igal Szleifer, Vinayak P. Dravid, and Vadim Backman. Analysis of three-dimensional chromatin packing domains by chromatin scanning transmission electron microscopy (ChromSTEM). Scientific Reports, 12(1):12198, July 2022. ISSN 2045-2322. doi: 10.1038/s41598-022-16028-2. URL https://www.nature.com/articles/s41598-022-16028-2. Publisher: Nature Publishing Group.

[76] Michael Rubinstein and Ralph H. Colby. Polymer physics. Oxford university press, 2003. URL https://academic.oup.com/book/54754.

[77] Elisa Oberbeckmann, Kimberly Quililan, Patrick Cramer, and A. Marieke Oudelaar. In vitro reconstitution of chromatin domains shows a role for nucleosome positioning in 3D genome organization. Nature Genetics, pages 1–10, January 2024. ISSN 1546-1718. doi: 10.1038/s41588-023-01649-8. URL https://www.nature.com/articles/s41588-023-01649-8. Publisher: Nature Publishing Group.

[78] Oliver Wiese, Davide Marenduzzo, and Chris A. Brackley. Nucleosome positions alone can be used to predict domains in yeast chromosomes. Proceedings of the National Academy of Sciences, 116(35):17307–17315, August 2019. doi: 10.1073/pnas.1817829116. URL https://www.pnas.org/doi/full/10.1073/pnas.1817829116. Publisher: Proceedings of the National Academy of Sciences.

[79] Stephanie Portillo-Ledesma, Lucille H. Tsao, Meghna Wagley, Melike Lakadamyali, Maria Pia Cosma, and Tamar Schlick. Nucleosome Clutches are Regulated by Chromatin Internal Parameters. Journal of Molecular Biology, 433(6):166701, March 2021. ISSN 0022-2836. doi: 10.1016/j.jmb.2020.11.001. URL https://www.sciencedirect.com/science/article/pii/S0022283620306197.

[80] K. S. Beckwith, Ø Ødegård Fougner, N. R. Morero, C. Barton, F. Schueder, W. Tang, S. Alexander, J.-M. Peters, R. Jungmann, E. Birney, and J. Ellenberg. Nanoscale 3D DNA tracing in non-denatured cells resolves the Cohesin-dependent loop architecture of the genome in situ. Nature Communications, 16(1):6673, July 2025. ISSN 2041-1723. doi: 10.1038/s41467-025-61689-y. URL https://www.nature.com/articles/s41467-025-61689-y. Publisher: Nature Publishing Group.

[81] Brian Chan and Michael Rubinstein. Activity-driven chromatin organization during interphase: Compaction, segregation, and entanglement suppression. Proceedings of the National Academy of Sciences, 121(21):e2401494121, May 2024. doi: 10.1073/pnas.2401494121. URL https://www.pnas.org/doi/abs/10.1073/pnas.2401494121. Publisher: Proceedings of the National Academy of Sciences.

[82] Suhas S. P. Rao, Miriam H. Huntley, Neva C. Durand, Elena K. Stamenova, Ivan D. Bochkov, James T. Robinson, Adrian L. Sanborn, Ido Machol, Arina D. Omer, Eric S. Lander, and Erez Lieberman Aiden. A 3D Map of the Human Genome at Kilobase Resolution Reveals Principles of Chromatin Looping. Cell, 159(7):1665–1680, December 2014. ISSN 0092-8674, 1097-4172. doi: 10.1016/j.cell.2014.11.021. URL https://www.cell.com/cell/abstract/S0092-8674(14)01497-4. Publisher: Elsevier.

[83] Betul Akgol Oksuz, Liyan Yang, Sameer Abraham, Sergey V. Venev, Nils Krietenstein, Krishna Mohan Parsi, Hakan Ozadam, Marlies E. Oomen, Ankita Nand, Hui Mao, Ryan MJ Genga, Rene Maehr, Oliver J. Rando, Leonid A. Mirny, Johan Harmen Gibcus, and Job Dekker. Systematic evaluation of chromosome conformation capture assays, December 2020. URL https://www.biorxiv.org/content/10.1101/2020.12.26.424448v2. Pages: 2020.12.26.424448 Section: New Results.

[84] Kartik Kamat, Zhuohan Lao, Yifeng Qi, Yuchuan Wang, Jian Ma, and Bin Zhang. Compartmentalization with nuclear landmarks yields random, yet precise, genome organization. Biophysical Journal, 122(7): 1376–1389, 2023. Publisher: Elsevier.

[85] Zhuohan Lao, Kartik Kamat, Zhongling Jiang, and Bin Zhang. OpenNucleome for high resolution nuclear structural and dynamical modeling. eLife, 13, February 2024. doi: 10.7554/eLife.93223.1. URL https://elifesciences.org/reviewed-preprints/93223. Publisher: eLife Sciences Publications Limited.

[86] Benot Roux and Jonathan Weare. On the statistical equivalence of restrained-ensemble simulations with the maximum entropy method. The Journal of Chemical Physics, 138(8):084107, February 2013. ISSN 0021-9606. doi: 10.1063/1.4792208. URL https://www.ncbi.nlm.nih.gov/pmc/articles/PMC3598863/.

[87] Marco Dombrowski, Maik Engeholm, Christian Dienemann, Svetlana Dodonova, and Patrick Cramer. Histone H1 binding to nucleosome arrays depends on linker DNA length and trajectory. Nature Structural & Molecular Biology, 29(5):493–501, May 2022. ISSN 1545-9985. doi: 10.1038/s41594-022-00768-w. URL https://www.nature.com/articles/s41594-022-00768-w. Number: 5 Publisher: Nature Publishing Group.

[88] Lin Xingcheng and Zhang Bin. Explicit Ion Modeling Predicts Physicochemical Interactions for Chromatin Organization. eLife, 12, August 2023. doi: 10.7554/eLife.90073. URL https://elifesciences.org/reviewed-preprints/90073. Publisher: eLife Sciences Publications Limited.

[89] Shuming Liu, Advait Athreya, Zhuohan Lao, and Bin Zhang. From Nucleosomes to Compartments: Physicochemical Interactions Underlying Chromatin Organization. Annu. Rev. Biophys., 53(1):annurev–biophys–030822–032650, May 2024. doi: 10.1146/annurev-biophys-030822-032650

[90] Erica M. Hildebrand, Kirill Polovnikov, Bastiaan Dekker, Yu Liu, Denis L. Lafontaine, A. Nicole Fox, Ying Li, Sergey V. Venev, Leonid A. Mirny, and Job Dekker. Mitotic chromosomes are self-entangled and disentangle through a topoisomerase-II-dependent two-stage exit from mitosis. Molecular Cell, 84(8):1422–1441.e14, April 2024. ISSN 1097-2765. doi: 10.1016/j.molcel.2024.02.025. URL https://www.cell.com/molecular-cell/abstract/S1097-2765(24)00144-8. Publisher: Elsevier.

[91] Peter Eastman, Raimondas Galvelis, Rauí. P. Peláez, Charlles R. A. Abreu, Stephen E. Farr, Emilio Gallicchio, Anton Gorenko, Michael M. Henry, Frank Hu, Jing Huang, Andreas Kramer, Julien Michel, Joshua A. Mitchell, Vijay S. Pande, João PGLM Rodrigues, Jaime Rodriguez-Guerra, Andrew C. Simmonett, Sukrit Singh, Jason Swails, Philip Turner, Yuanqing Wang, Ivy Zhang, John D. Chodera, Gianni De Fabritiis, and Thomas E. Markland. OpenMM 8: Molecular Dynamics Simulation with Machine Learning Potentials. The Journal of Physical Chemistry B, 128(1):109–116, January 2024. ISSN 1520-6106. doi: 10.1021/acs.jpcb.3c06662. URL https://doi.org/10.1021/acs.jpcb.3c06662. Publisher: American Chemical Society

[92] Zhijun Zhang, Xinzijian Liu, Kangyu Yan, Mark E. Tuckerman, and Jian Liu. Unified Efficient Thermostat Scheme for the Canonical Ensemble with Holonomic or Isokinetic Constraints via Molecular Dynamics. The Journal of Physical Chemistry A, 123(28):6056–6079, July 2019. ISSN 1089-5639. doi: 10.1021/acs.jpca.9b02771. URL https://doi.org/10.1021/acs.jpca.9b02771. Publisher: American Chemical Society

