## Supplementary Information for "Euchromatin forms condensed domains with short active regions on the surface"

Joseph M. Paggi, Bin Zhang*


Department of Chemistry, Massachusetts Institute of Technology, Cambridge, MA, USA

**2 View of neighbor balancing as correction for competition between ligation events** **4**

**3 Details of maximum entropy inversion** **4**

**4 Comparison to chromatin tracing experiments** **4**

**5 Analysis of 3D structures** **5**

**6 Epigenetics data** **6**

**7 Supplementary figures** **10**

#### Micro-C data processing

##### Alignment and filtering

Reads were aligned to the mm39 genome using bowtie2 with options "–reorder –local –very-sensitive-local –maxins 1 –minins 1000000" [1]. These constraints on the insert size are designed to be unsatisfiable, resulting in no "concordantly" aligning reads. This removes a bias towards short-range inward-oriented alignments [2]. The read alignments are then transferred to pairs format using "pairtools parse" with options "–walks-policy mask –add-mapq –min-mapq 2" [3]. Duplicate reads are then removed using "pairtools dedup" with option -"-max-mismatch 1". Only reads where both ends align uniquely to a capture region are retained.

#### Mapping reads to expected nucleosome center positions

A custom analysis script is used to shift the assigned positions to the most probable nucleosome center positions according to the read pair orientation, i.e. the strand to which each read end aligns, as described below and illustrated in Figure S1a. We acknowledge that not every nucleosome will have the same length protected DNA and that factors other than nucleosomes, such as transcription factors, which might protect different lengths of DNA are likely the source of some reads. In any case, the shifted position is a better guess for the center position than the 5' of the read end.

- Upstream +/ Downstream -: "inward", result of ligation between the 3' end of an upstream nucleosome and the 5' end of a downstream nucleosome, shift first + protected / 2, shift second - protected / 2
- Upstream - / Downstream +: "outward", result of ligation between the 5' end of an upstream nucleosome and the 3' end of a downstream nucleosome, shift first - protected / 2, shift second + protected / 2
- Upstream - / Downstream +: "tandem entry", result of ligation between the 5' end of an upstream nucleosome and the 5' end of a downstream nucleosome, shift first - protected / 2, shift second - protected / 2
- Upstream +/ Downstream +: "tandem exit", result of ligation between the 3' end of an upstream nucleosome and the 3' end of a downstream nucleosome, shift first + protected / 2, shift second + protected / 2

The number of base pairs protected by a nucleosome was set to 131 as this value maximizes the concordance between the P(s) curve for the different read orientations (Figure S1b). This is inconsistent with the textbook view that 147 bps are protected by each nucleosome, however, a shift of 131 maximizing agreement between read orientations is in agreement with previous work [4].

The genomic region we plan to simulate is divided into 200 bp windows. We assume that a nucleosome is positioned in the center of each of these bins. For each read, we assign each end to a nucleosome corresponding to its aligned genomic position, after applying the shift described above. Read pairs are converted to a contact map using cooler [5].

#### Removal of likely unligated reads

After shifting, we found that the abundances and distance-dependence of all read directions was concordant, except for an overabundance of "inward" reads between adjacent nucleosomes (Figure S1b). These "inward" reads are in the expected orientation for unligated DNA fragments, and previous work suggests that these reads arise from unligated DNA fragments that escape exonuclease mediated degradation [6]. To correct for the overabundance of close-range "inward" reads, we discarded "inward" reads assigned to i+1 or i+2 nucleosomes and multiplied the remaining counts by 4/3. This operation assumes that true inward ligation events occur at approximately the same rate as other types of junctions. This step, along with shifting the reads according to their read direction, dramatically alters the P(s) curve at short range (Figure S1d).

#### ICE balancing and interpolation of regions with low coverage and low capture probe coverage

ICE balancing is used to account for variations in coverage across the genome [7]. We made several modifications to the approach to make it more amenable for use with region-capture data.

First, we use modified filtering rules to explicitly mask pixels with low capture probe coverage. The entirety of the region of interest is not covered by capture probes to avoid including probes with sequences that occur multiple places in the genome, as this would lower the enrichment rate for the region of interest. This poses a problem for ICE balancing. ICE balancing assumes that biases in read counts are multiplicative, e.g. if a region contains twice as many free DNA ends after MNase digestion, two times as many contacts are observed for all pairs including the region. However, low capture probe coverage is instead additive as a ligated section of DNA can be captured by either end. If one end of a pair has no coverage, it can still be captured efficiently if the other end has capture probe coverage, but if the other end does not have capture probe coverage, then it will rarely be captured. This results in problematic gaps in coverage for pairs with no coverage in either end. These are particularly common for i+1 contacts as uncovered regions often extend greater than 200 bps, introducing artifacts in the neighbor contact frequencies used in neighbor balancing. These regions are not always filtered out with standard masking protocols as they can still contain a large number of reads. To deal with this we explicitly mask pixels where the total capture probe coverage is below 20%. This removes the problematic gaps in coverage and stabilizes the estimated neighbor contact probabilities (Figure S2).

Second, we modify the ICE balancing algorithm to account for the fact that we are only considering a subset of the genome. Only considering a subset of the genome introduces a boundary effect to the contact maps. About half of the contacts for nucleosomes at the very ends of the region of interest are to out of region nucleosomes and thereby not considered in ICE balancing. Applying ICE balancing thereby artificially doubles the contact frequency for these nucleosomes. To account for this, we modify the protocol for computing marginal contacts in the ICE balancing protocol. For each position, we infer the number of missing contacts by extending the contact map assuming that the missing values have the average contact frequency for their corresponding genomic separation.

Finally, we integrate an interpolation step to prevent masking from perturbing the contact density in the simulations. The standard practice is to treat masked values as zeros during ICE balancing. However, in our simulations (and in reality) these masked contacts occur with non-zero frequency. Thereby nucleosomes with a high number of masked contacts end up having inflated contact densities when these masked contacts are added on top of the unmasked values. These artificial variations in contact density are of about the same magnitude as the true variations, so treating this issue is essential. To address this issue, we employ a three phase ICE balancing protocol. The first phase is performed as usual. In the second phase, between each iteration, masked values are interpolated. Interpolation is performed across each diagonal of the contact map to avoid perturbing the relationship between genomic separation and contact frequency. The third stage is standard ICE balancing with the interpolated values included but not re-computed between each stage. The first phase is necessary as we found that interpolating from the beginning can result in the algorithm not converging. The third step is necessary to ensure that all rows sum to the same value as interpolating between each iteration can result in small deviations.

#### Neighbor balancing protocol

The neighbor balanced contact frequencies $C^{(\text{neighbor})}$ are computed from the ICE balanced contact frequencies $C$ as

$$C_{ij}^{(\text{neighbor})} = \frac{C_{ij}}{\sqrt{\frac{1}{2}(C_{i,j-1} + C_{i,j+1})\frac{1}{2}(C_{i-1,j} + C_{i+1,j})}}$$

The ICE balanced contact frequencies are smoothed using a Gaussian kernel smoother with a standard deviation of 1 bin when computing the denominator of this expression.

Note that if ICE balanced neighbor contacts were constant over a region, then this procedure would exactly update the neighbor contact frequencies to be one. In practice, the neighbor contacts are not constant and this procedure does not perfectly set the neighbor contact frequencies to one. We considered performing iterative updates to bring the neighbor contact frequencies closer to one, but we found that this often results in rapidly oscillating contact densities. In the future, it may be possible to develop a procedure that explicitly optimizes for smooth contact densities and neighbor contact frequencies close to 1, combining the ICE and neighbor balancing objectives.

#### Contact map smoothing

The contact map is smoothed prior to being used in simulations (Fig. S2). Despite the fact that RCMC produces much higher-resolution contact maps than typical micro-C experiments, we found that at 200 bp resolution some smoothing is required to obtain maps that are not visibly pixelated. Pixels far from the diagonal have particularly high variance as they generally have lower contact frequencies and thereby fewer reads to estimate the contact frequency. To maintain higher-resolution where possible, we smooth the matrix more aggressively far from the diagonal and not at all near the diagonal. We smooth the map by grouping squares of pixels and assigning all pixels the average value. The squares are doubled in size at farther distances from the diagonal to maintain a consistent average number of reads per group.

#### Neighbor balancing on genome-wide micro-C maps

Genome-wide micro-C data are generally too sparse to view at nucleosome resolution (200 bps), so some alterations needed to be made to the neighbor balancing protocol. To perform neighbor balancing on lower resolution maps, e.g. 25.6 kb resolution, we first compute ICE balanced contact maps at both 200 bp resolution and the desired resolution. Second, we compute the neighbor contact frequencies in the nucleosome-resolution map, as described above. Third, for each lower resolution bin, we average the respective nucleosome-resolution neighbor contact frequencies. Finally, the neighbor balancing update is performed as described above using these averaged neighbor contact frequencies.

### View of neighbor balancing as correction for competition between ligation events

In this work, we assume that the probability of a ligation occurring between two nucleosomes (or, more accurately, two fragments of nucleosome protected DNA) is proportional only to the distance between the two nucleosomes. However, in a single cell, each nucleosome can only ligate productively to one other nucleosome. Therefore, a more accurate model would be to assume that nucleosomes are in competition with one another to be ligated. In effect, nucleosomes surrounded by many nucleosomes will have a lower probability of ligating to any given nucleosome than a nucleosome in a more dilute environment.

Neighbor balancing approximately acts to convert a contact map generated under this competitive process to one generated purely based on pairwise distances assumed in the rest of this study. Intuitively, a nucleosome in a dense environment will have lower neighbor contact frequencies as these interactions are competed for by many other nucleosomes. Consistent with this, neighbor balancing increases the contact density for nucleosomes with low neighbor contacts.

### Details of maximum entropy inversion

In MEI, the energy coefficients are typically solved for using Newton's method [8]. However, considering large regions at 200 bp resolution makes the Hessian matrix too large to fit into memory as it's size scales with the number of monomers to the fourth. To enable simulations, we instead use gradient descent:

$$\alpha_{ij}^{t+1} = \alpha_{ij}^{t} + \gamma(T_{ij} - \hat{S}_{ij}^{t})$$

where $\hat{S}^t$ is an iteration averaged matrix of simulated contact frequencies, $T$ is the matrix of target contact frequencies, and $\gamma$ is the learning rate.

$\hat{S}^t$ is defined as an exponential moving average of the contact maps for each iteration $\hat{S}^t$

$$\hat{S}_{ij}^{(t)} = (1-\beta)S_{ij}^{(t)} + \beta\hat{S}_{ij}^{(t-1)}$$

This moving average is not used in the first five rounds of optimization as the contact frequencies change rapidly, so the contact frequencies from previous rounds are not informative. In all simulations, $\gamma = 0.3$ and $\beta = 0.75$.

For each iteration, 16 simulations are performed for 11,000,000 steps each, a frame is stored every 10,000 steps, and the first 1,000,000 steps are excluded when computing contact maps. This procedure is run until the simulated P(s) curve closely matches the reference. The simulated contact maps were visually assessed for key features, such as matching contact densities, bulk insulation scores, and presence of micro-compartment dots.

In the first iteration, the initial conformation is created by placing each nucleosome on a Hilbert curve, which provides a dense conformation with no knots. Subsequent iterations are initiated from the final frame of the previous iteration to reduce the required equilibration time.

### Comparison to chromatin tracing experiments

The coordinates determined by chromatin tracing contain measurement noise. Adding noise to coordinates tends to inflate the distances between them, especially if the true distances are of comparable magnitude to the measurement noise. We therefore needed to account for measurement noise to accurately assess whether our simulations reproduce the distance scaling observed in experiments.

To estimate the measurement noise distribution in ORCA data for the *Nanog* locus [9], we used control data in which one region was imaged twice per chromatid. Assuming that the imaged region does not move, the distances between these subsequent measurements can be modeled as the difference in position between two independent samples from the noise distribution. Therefore, we searched for an underlying noise distribution for which the difference between two samples matches the empirical distribution of distances between images. Initially, we expected the noise distribution to be normally distributed, but the empirical distribution has too heavy of a tail to be modeled with a normal distribution. We expect this reflects some of the measurement noise corresponding to mis-identification of the region in the image. We found that a scaled t-distribution, which provides an additional shape parameter controlling the weight of the tails, provided a much better fit. We performed a grid search to find the scale and shape parameters maximizing the likelihood of the empirical distribution (Figure S7a). We fit parameters separately using only the x and y coordinates, as these appeared to have less measurement noise than the z coordinates (Figure S7b).

For the *Sox2* region [10], no control data is available. Instead, we solved for a noise distribution that, when added to the simulated coordinates, brought the distribution of distances between loci separated by 60 kb into agreement with the ORCA measurements (Figure S7c). This approach effectively assumes that the simulated distances at this scale are on average correct. Thereby, we cannot assess if the absolute distance values are correct. However, this approach allows meaningful assessment of the overall distance scaling trend and relative distances between specific loci. Comparison to live cell imaging data from Li et al. [11] provides support for the absolute scale of distances (Figure S7d).

### Analysis of 3D structures

#### Assignment of packing domains using DBSCAN

DBSCAN [12], as implemented in scikit-learn [13], was used to decompose chromatin conformations into a set of packing domains and protrusions. DBSCAN has two tunable parameters: epsilon, which controls the maximum distance at which two points are considered neighboring, and min_samples, which controls the number of neighbors needed for a point to be considered a "core" point. Epsilon was set to 30 nm and min_samples was set to 14. This combination of parameters results in a nucleosome concentration of at least 205 µM being required at the core of a domain. The clusters identified by DBSCAN were considered packing domains and the remaining "noise" points are considered protrusions. Protrusions less than 3 nucleosomes long that exit and return to the same packing domain were relabeled as part of the packing domain. A two nucleosome protrusion was added between packing domains with no protrusion linking between them.

This procedure yields packing domains that visually correspond to densely grouped nucleosomes with continuously high density. Naturally, changing the parameters or clustering algorithm would yield quantitatively different results. In particular, substantially decreasing min_samples would result in more nucleosomes being identified as in domains, whereas increasing it would result in fewer nucleosomes being identified as in domains. Importantly, increasing min_samples to 20, corresponding to an core density of at least 300 µM, increases the protrusion probability homogeneously for the 5 regions studied Table 1, suggesting that there is not an excess of nucleosomes just on the edge of being called a protrusion in gene rich regions when using our chosen parameters. Requiring this higher density resulted in nucleosomes that visually appeared as part of a condensed domain being called a protrusion, motivating our decision to require a lower density at the core of domains.

| Region | 200 µM | 300 µM | Ratio |
|---|---|---|---|
| *Nanog* | 0.166 | 0.445 | 2.68 |
| *Sox2* | 0.049 | 0.194 | 3.96 |
| *Fbn2* | 0.063 | 0.250 | 3.97 |
| *Klf1* | 0.135 | 0.386 | 2.86 |
| *Ppm1g* | 0.050 | 0.189 | 3.78 |

Table 1: **Protrusion probabilities increase by roughly the same factor when the required core nucleosome density is increased.**

#### Assignment of contiguous nucleosome clutches using insulation scores

Clutches were assigned by considering single frame insulation scores. First, a contact map was computed using a capture radius of 40 nm. Then, insulation scores were computed through cooltools using a window size of ten nucleosomes and ignoring the first two diagonals [14]. Finally, positions with an insulation score less than half the median value across all systems were called as boundaries and the remaining contiguous blocks were defined as contiguous nucleosome clutches (Figure S13a). This choice of window size and boundary cutoff yielded the largest clutches that are largely unmixed with each other (Figure S14c,e).

##### Estimating the density of domains

Estimating the density of domains was surprisingly challenging. Their irregular shape and internal cavities makes it unclear how to define their surface and different approaches resulted in different density estimates (Figure S9a-d). Here we summarize the methods we considered and comment on their results. First, we considered using the probe excluded volume. This method gave inflated estimates for small domains because the surface was drawn precisely around the nucleosomes, whereas it gave deflated estimates for large domains because the packing domains often contain unfilled cavities (Figure S9d,g). Second, we augmented this approach by padding the nucleosome diameters to prevent inflated estimates for small domains. Specifically, we padded the nucleosome radii to 11 nm, which is roughly half the average distance between nucleosomes inside of packing domains (Figure S9h). Third, we considered the maximum density inside of spheres centered on each nucleosome in a domain (Figure S9c). This yielded substantially higher densities than the other methods (Figure S9c,f). Fourth, we used the radius of gyration to characterize the size of packing domains. This approach does not provide a density, but was useful as it contains no free parameters and does not appear to display any significant size dependent bias. Finally, We considered the relationship between the time a nucleosome spends on the surface and it's average local density. These quantities are linearly related and if we extrapolate the trend to no time spent on the surface, we obtain an average local density of about 300 µM.

While these different approaches yield different absolute densities, the values are very similar between systems, after accounting for size dependent biases. The farthest deviation from this trend is that large packing domains in the *Ppm1g* are modestly denser than those in the other systems.

#### Epigenetics data

The pybigwig [15] to load epigenetics tracks and UCSC liftover tool [16] to mm39 where appropriate. Where available, we used signal p-value tracks from ENCODE [17, 18].

| | **Source** | **Accession** | **Citation** |
|---|---|---|---|
| **Condensability** | GEO | GSE252941 | Park et al. [19] |
| **ATAC** | GEO | GSE98390 | King et al. [20] |
| **H3K27ac** | ENCODE | ENCFF230RNU | Sethi et al. [21] |
| **H3K4me1** | ENCODE | ENCFF410CGG | The ENCODE Project Consortium [17] |
| **H3K4me3** | ENCODE | ENCFF523UIR | [17] |
| **H3K27me3** | ENCODE | ENCFF160FEV | He et al. [22] |
| **H3K9me3** | ENCODE | ENCFF293DGT | The ENCODE Project Consortium [17] |
| **H1** | GEO | GSM1124783 | Cao et al. [23] |
| **RNA-seq** | GEO | GSE123636 | Hansen et al. [24] |
| **PolII** | GEO | GSM6809981 | Goel et al. [25] |

Table 2: **Sources of epigenetics data.** For condensability, the average of GSE252941_E14_NCP_sp_1rep_10kb_score.gtab.txt.gz and GSE252941_E14_NCP_sp_2rep_10kb_score.gtab.txt.gz at Sample ID 8 was used. For, H1 GSM1124783 H1d-1 IP-IN.bw was used. For RNA-seq, GSE123636_C59_1_2_RNAseq_coverage.bw was used.

#### References

[1] Ben Langmead and Steven L. Salzberg. Fast gapped-read alignment with Bowtie 2. *Nature Methods*, 9(4): 357–359, April 2012. ISSN 1548-7105. doi: 10.1038/nmeth.1923. URL https://www.nature.com/articles/nmeth.1923. Publisher: Nature Publishing Group.

[2] Houda Belaghzal, Job Dekker, and Johan H. Gibcus. Hi-C 2.0: An optimized Hi-C procedure for high-resolution genome-wide mapping of chromosome conformation. *Methods*, 123:56–65, July 2017. ISSN 1046-2023. doi: 10.1016/j.ymeth.2017.04.004. URL https://www.sciencedirect.com/science/article/pii/S1046202316304662.

[3] Open2C, Nezar Abdennur, Geoffrey Fudenberg, Ilya M. Flyamer, Aleksandra A. Galitsyna, Anton Goloborodko, Maxim Imakaev, and Sergey V. Venev. Pairtools: From sequencing data to chromosome con-

tacts. *PLOS Computational Biology*, 20(5):e1012164, May 2024. ISSN 1553-7358. doi: 10.1371/journal.pcbi.1012164. URL https://journals.plos.org/ploscompbiol/article?id=10.1371/journal.pcbi.1012164. Publisher: Public Library of Science.

[4] Masae Ohno, Tadashi Ando, David G. Priest, Vipin Kumar, Yamato Yoshida, and Yuichi Taniguchi. Sub-nucleosomal Genome Structure Reveals Distinct Nucleosome Folding Motifs. *Cell*, 176(3):520–534.e25, January 2019. ISSN 0092-8674. doi: 10.1016/j.cell.2018.12.014. URL https://www.sciencedirect.com/science/article/pii/S0092867418316283.

[5] Nezar Abdennur and Leonid A Mirny. Cooler: scalable storage for Hi-C data and other genomically labeled arrays. *Bioinformatics*, 36(1):311–316, January 2020. ISSN 1367-4803. doi: 10.1093/bioinformatics/btz540. URL https://doi.org/10.1093/bioinformatics/btz540.

[6] Tsung-Han S. Hsieh, Assaf Weiner, Bryan Lajoie, Job Dekker, Nir Friedman, and Oliver J. Rando. Mapping Nucleosome Resolution Chromosome Folding in Yeast by Micro-C. *Cell*, 162(1):108–119, July 2015. ISSN 0092-8674. doi: 10.1016/j.cell.2015.05.048. URL https://www.sciencedirect.com/science/article/pii/S0092867415006388.

[7] Maxim Imakaev, Geoffrey Fudenberg, Rachel Patton McCord, Natalia Naumova, Anton Goloborodko, Bryan R. Lajoie, Job Dekker, and Leonid A. Mirny. Iterative correction of Hi-C data reveals hallmarks of chromosome organization. *Nature Methods*, 9(10):999–1003, October 2012. ISSN 1548-7105. doi: 10.1038/nmeth.2148. URL https://www.nature.com/articles/nmeth.2148. Publisher: Nature Publishing Group.

[8] Benoît Roux and Jonathan Weare. On the statistical equivalence of restrained-ensemble simulations with the maximum entropy method. *The Journal of Chemical Physics*, 138(8):084107, February 2013. ISSN 0021-9606. doi: 10.1063/1.4792208. URL https://www.ncbi.nlm.nih.gov/pmc/articles/PMC3598863/.

[9] Taihei Fujimori, Carolina Rios-Martinez, Abby R. Thurm, Michaela M. Hinks, Benjamin R. Doughty, Joydeb Sinha, Derek Le, Antonina Hafner, William J. Greenleaf, Alistair N. Boettiger, and Lacramioara Bintu. Single-cell chromatin state transitions during epigenetic memory formation. *bioRxiv*, page 2023.10.03.560616, October 2023. doi: 10.1101/2023.10.03.560616. URL https://www.ncbi.nlm.nih.gov/pmc/articles/PMC10592931/.

[10] Antonina Hafner, Minhee Park, Scott E. Berger, Sedona E. Murphy, Elphège P. Nora, and Alistair N. Boettiger. Loop stacking organizes genome folding from TADs to chromosomes. *Molecular Cell*, 83(9):1377–1392.e6, May 2023. ISSN 1097-2765. doi: 10.1016/j.molcel.2023.04.008. URL https://www.sciencedirect.com/science/article/pii/S1097276523002526.

[11] Jieru Li, Angela Hsu, Yujing Hua, Guanshi Wang, Lingling Cheng, Hiroshi Ochiai, Takashi Yamamoto, and Alexandros Pertsinidis. Single-gene imaging links genome topology, promoter–enhancer communication and transcription control. *Nature Structural & Molecular Biology*, 27(11):1032–1040, November 2020. ISSN 1545-9985. doi: 10.1038/s41594-020-0493-6. URL https://www.nature.com/articles/s41594-020-0493-6. Publisher: Nature Publishing Group.

[12] Martin Ester, Hans-Peter Kriegel, Jörg Sander, and Xiaowei Xu. A density-based algorithm for discovering clusters in large spatial databases with noise. In *Proceedings of the Second International Conference on Knowledge Discovery and Data Mining*, KDD'96, pages 226–231, Portland, Oregon, August 1996. AAAI Press.

[13] Fabian Pedregosa, Gaël Varoquaux, Alexandre Gramfort, Vincent Michel, Bertrand Thirion, Olivier Grisel, Mathieu Blondel, Peter Prettenhofer, Ron Weiss, Vincent Dubourg, Jake Vanderplas, Alexandre Passos, David Cournapeau, Matthieu Brucher, Matthieu Perrot, and Édouard Duchesnay. Scikit-learn: Machine Learning in Python. *Journal of Machine Learning Research*, 12(85):2825–2830, 2011. ISSN 1533-7928. URL http://jmlr.org/papers/v12/pedregosa11a.html.

[14] Open2C, Nezar Abdennur, Sameer Abraham, Geoffrey Fudenberg, Ilya M. Flyamer, Aleksandra A. Galitsyna, Anton Goloborodko, Maxim Imakaev, Betul A. Oksuz, Sergey V. Venev, and Yao Xiao. Cooltools: Enabling high-resolution Hi-C analysis in Python. *PLOS Computational Biology*, 20(5):e1012067, May 2024. ISSN 1553-7358. doi: 10.1371/journal.pcbi.1012067. URL https://journals.plos.org/ploscompbiol/article?id=10.1371/journal.pcbi.1012067. Publisher: Public Library of Science.

[15] Fidel Ramírez, Devon P Ryan, Björn Grüning, Vivek Bhardwaj, Fabian Kilpert, Andreas S Richter, Steffen Heyne, Friederike Dündar, and Thomas Manke. deepTools2: a next generation web server for deep-sequencing data analysis. *Nucleic Acids Research*, 44(W1):W160–W165, July 2016. ISSN 0305-1048. doi: 10.1093/nar/gkw257. URL https://doi.org/10.1093/nar/gkw257.

[16] A. S. Hinrichs, D. Karolchik, R. Baertsch, G. P. Barber, G. Bejerano, H. Clawson, M. Diekhans, T. S. Furey, R. A. Harte, F. Hsu, J. Hillman-Jackson, R. M. Kuhn, J. S. Pedersen, A. Pohl, B. J. Raney, K. R. Rosenbloom, A. Siepel, K. E. Smith, C. W. Sugnet, A. Sultan-Qurraie, D. J. Thomas, H. Trumbower, R. J. Weber, M. Weirauch, A. S. Zweig, D. Haussler, and W. J. Kent. The UCSC Genome Browser Database: update 2006. *Nucleic Acids Research*, 34(suppl_1):D590–D598, January 2006. ISSN 0305-1048. doi: 10.1093/nar/gkj144. URL https://doi.org/10.1093/nar/gkj144.

[17] The ENCODE Project Consortium. An integrated encyclopedia of DNA elements in the human genome. *Nature*, 489(7414):57–74, September 2012. ISSN 1476-4687. doi: 10.1038/nature11247. URL https://www.nature.com/articles/nature11247. Publisher: Nature Publishing Group.

[18] Benjamin C. Hitz, Jin-Wook Lee, Otto Jolanki, Meenakshi S. Kagda, Keenan Graham, Paul Sud, Idan Gabdank, J. Seth Strattan, Cricket A. Sloan, Timothy Dreszer, Laurence D. Rowe, Nikhil R. Podduturi, Venkat S. Malladi, Esther T. Chan, Jean M. Davidson, Marcus Ho, Stuart Miyasato, Matt Simison, Forrest Tanaka, Yunhai Luo, Ian Whaling, Eurie L. Hong, Brian T. Lee, Richard Sandstrom, Eric Rynes, Jemma Nelson, Andrew Nishida, Alyssa Ingersoll, Michael Buckley, Mark Frerker, Daniel S. Kim, Nathan Boley, Diane Trout, Alex Dobin, Sorena Rahmanian, Dana Wyman, Gabriela Balderrama-Gutierrez, Fairlie Reese, Neva C. Durand, Olga Dudchenko, David Weisz, Suhas S. P. Rao, Alyssa Blackburn, Dimos Gkountaroulis, Mahdi Sadr, Moshe Olshansky, Yossi Eliaz, Dat Nguyen, Ivan Bochkov, Muhammad Saad Shamim, Ragini Mahajan, Erez Aiden, Tom Gingeras, Simon Heath, Martin Hirst, W. James Kent, Anshul Kundaje, Ali Mortazavi, Barbara Wold, and J. Michael Cherry. The ENCODE Uniform Analysis Pipelines, April 2023. URL https://www.biorxiv.org/content/10.1101/2023.04.04.535623v1. Pages: 2023.04.04.535623 Section: New Results.

[19] Sangwoo Park, Raquel Merino-Urteaga, Violetta Karwacki-Neisius, Gustavo Ezequiel Carrizo, Advait Athreya, Alberto Marin-Gonzalez, Nils A. Benning, Jonghan Park, Michelle M. Mitchener, Natarajan V. Bhanu, Benjamin A. Garcia, Bin Zhang, Tom W. Muir, Erika L. Pearce, and Taekjip Ha. Native nucleosomes intrinsically encode genome organization principles. *Nature*, 643(8071):572–581, July 2025. ISSN 1476-4687. doi: 10.1038/s41586-025-08971-7. URL https://www.nature.com/articles/s41586-025-08971-7. Publisher: Nature Publishing Group.

[20] Hamish W. King, Nadezda A. Fursova, Neil P. Blackledge, and Robert J. Klose. Polycomb repressive complex 1 shapes the nucleosome landscape but not accessibility at target genes. *Genome Research*, 28(10):1494–1507, October 2018. ISSN 1088-9051, 1549-5469. doi: 10.1101/gr.237180.118. URL https://genome.cshlp.org/content/28/10/1494. Company: Cold Spring Harbor Laboratory Press Distributor: Cold Spring Harbor Laboratory Press Institution: Cold Spring Harbor Laboratory Press Label: Cold Spring Harbor Laboratory Press Publisher: Cold Spring Harbor Lab.

[21] Anurag Sethi, Mengting Gu, Emrah Gumusgoz, Landon Chan, Koon-Kiu Yan, Joel Rozowsky, Iros Barozzi, Veena Afzal, Jennifer A. Akiyama, Ingrid Plajzer-Frick, Chengfei Yan, Catherine S. Novak, Momoe Kato, Tyler H. Garvin, Quan Pham, Anne Harrington, Brandon J. Mannion, Elizabeth A. Lee, Yoko Fukuda-Yuzawa, Axel Visel, Diane E. Dickel, Kevin Y. Yip, Richard Sutton, Len A. Pennacchio, and Mark Gerstein. Supervised enhancer prediction with epigenetic pattern recognition and targeted validation. *Nature Methods*, 17(8):807–814, August 2020. ISSN 1548-7105. doi: 10.1038/s41592-020-0907-8. URL https://www.nature.com/articles/s41592-020-0907-8. Publisher: Nature Publishing Group.

[22] Yupeng He, Manoj Hariharan, David U. Gorkin, Diane E. Dickel, Chongyuan Luo, Rosa G. Castanon, Joseph R. Nery, Ah Young Lee, Yuan Zhao, Hui Huang, Brian A. Williams, Diane Trout, Henry Amrhein, Rongxin Fang, Huaming Chen, Bin Li, Axel Visel, Len A. Pennacchio, Bing Ren, and Joseph R. Ecker. Spatiotemporal DNA methylome dynamics of the developing mouse fetus. *Nature*, 583(7818):752–759, July 2020. ISSN 1476-4687. doi: 10.1038/s41586-020-2119-x. URL https://www.nature.com/articles/s41586-020-2119-x. Publisher: Nature Publishing Group.

[23] Kaixiang Cao, Nathalie Lailler, Yunzhe Zhang, Ashwath Kumar, Karan Uppal, Zheng Liu, Eva K. Lee, Hongwei Wu, Magdalena Medrzycki, Chenyi Pan, Po-Yi Ho, Guy P. Cooper Jr, Xiao Dong, Christoph Bock,

Eric E. Bouhassira, and Yuhong Fan. High-Resolution Mapping of H1 Linker Histone Variants in Embryonic Stem Cells. *PLOS Genetics*, 9(4):e1003417, April 2013. ISSN 1553-7404. doi: 10.1371/journal.pgen.1003417. URL https://journals.plos.org/plosgenetics/article?id=10.1371/journal.pgen.1003417. Publisher: Public Library of Science.

[24] Anders S. Hansen, Tsung-Han S. Hsieh, Claudia Cattoglio, Iryna Pustova, Ricardo Saldaña-Meyer, Danny Reinberg, Xavier Darzacq, and Robert Tjian. Distinct Classes of Chromatin Loops Revealed by Deletion of an RNA-Binding Region in CTCF. *Molecular Cell*, 76(3):395–411.e13, November 2019. ISSN 1097-2765. doi: 10.1016/j.molcel.2019.07.039. URL https://www.sciencedirect.com/science/article/pii/S1097276519305945.

[25] Viraat Y. Goel, Nicholas G. Aboreden, James M. Jusuf, Haoyue Zhang, Luisa P. Mori, Leonid A. Mirny, Gerd A. Blobel, Edward J. Banigan, and Anders S. Hansen. Dynamics of microcompartment formation at the mitosis-to-G1 transition, September 2024. URL https://www.biorxiv.org/content/10.1101/2024.09.16.611917v1. Pages: 2024.09.16.611917 Section: New Results.

[26] Tsung-Han S. Hsieh, Claudia Cattoglio, Elena Slobodyanyuk, Anders S. Hansen, Oliver J. Rando, Robert Tjian, and Xavier Darzacq. Resolving the 3D Landscape of Transcription-Linked Mammalian Chromatin Folding. *Molecular Cell*, 78(3):539–553.e8, May 2020. ISSN 1097-2765. doi: 10.1016/j.molcel.2020.03.002. URL https://www.cell.com/molecular-cell/abstract/S1097-2765(20)30150-7. Publisher: Elsevier.

[27] Longzhi Tan, Dong Xing, Chi-Han Chang, Heng Li, and X. Sunney Xie. Three-dimensional genome structures of single diploid human cells. *Science*, 361(6405):924–928, August 2018. doi: 10.1126/science.aat5641. URL https://www.science.org/doi/10.1126/science.aat5641. Publisher: American Association for the Advancement of Science.

[28] Leonid A. Mirny. The fractal globule as a model of chromatin architecture in the cell. *Chromosome Research*, 19(1):37–51, January 2011. ISSN 1573-6849. doi: 10.1007/s10577-010-9177-0. URL https://doi.org/10.1007/s10577-010-9177-0.

[29] Viraat Y. Goel, Miles K. Huseyin, and Anders S. Hansen. Region Capture Micro-C reveals coalescence of enhancers and promoters into nested microcompartments. *Nature Genetics*, 55(6):1048–1056, June 2023. ISSN 1061-4036, 1546-1718. doi: 10.1038/s41588-023-01391-1. URL https://www.nature.com/articles/s41588-023-01391-1.

#### Supplementary figures

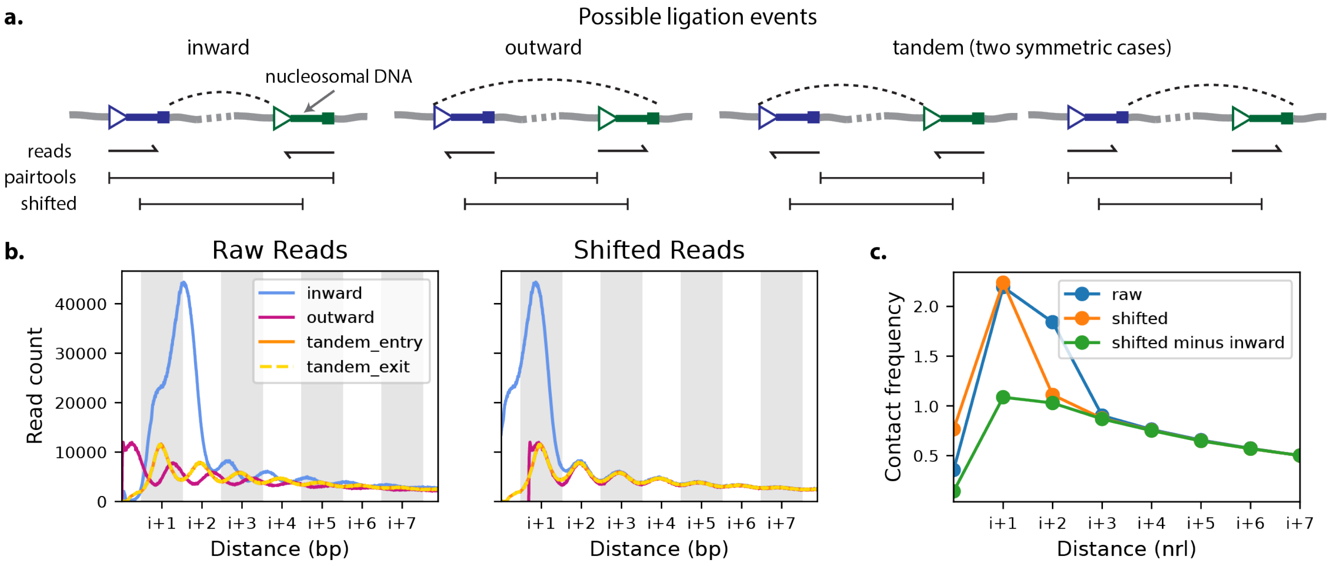

Fig. S1: **Impact of micro-C data processing on local interaction patterns.** (a) A cartoon depicting micro-C read orientations and the strategy for estimating nucleosome center positions (shifted). (b) Base resolution P(s) curves separated by read orientation. Using the 5'-ends of the reads the curves are out of phase with one another (raw reads), but after shifting to estimated nucleosome center positions they are mostly concordant. However, a disproportionate number of short range inwards oriented reads are present. (c) Nucleosome resolution P(s) curves with different processing protocols. For the raw data, the large peak in i+1 and i+2 reads stems from the large number of short range inwards reads. Shifting the reads to estimated nucleosome center positions moves this peak to the i+1 position. Removing inwards reads from the i+1 and i+2 bins and rescaling the remaining counts by 4/3 leaves a smooth curve (shifted minus inward).

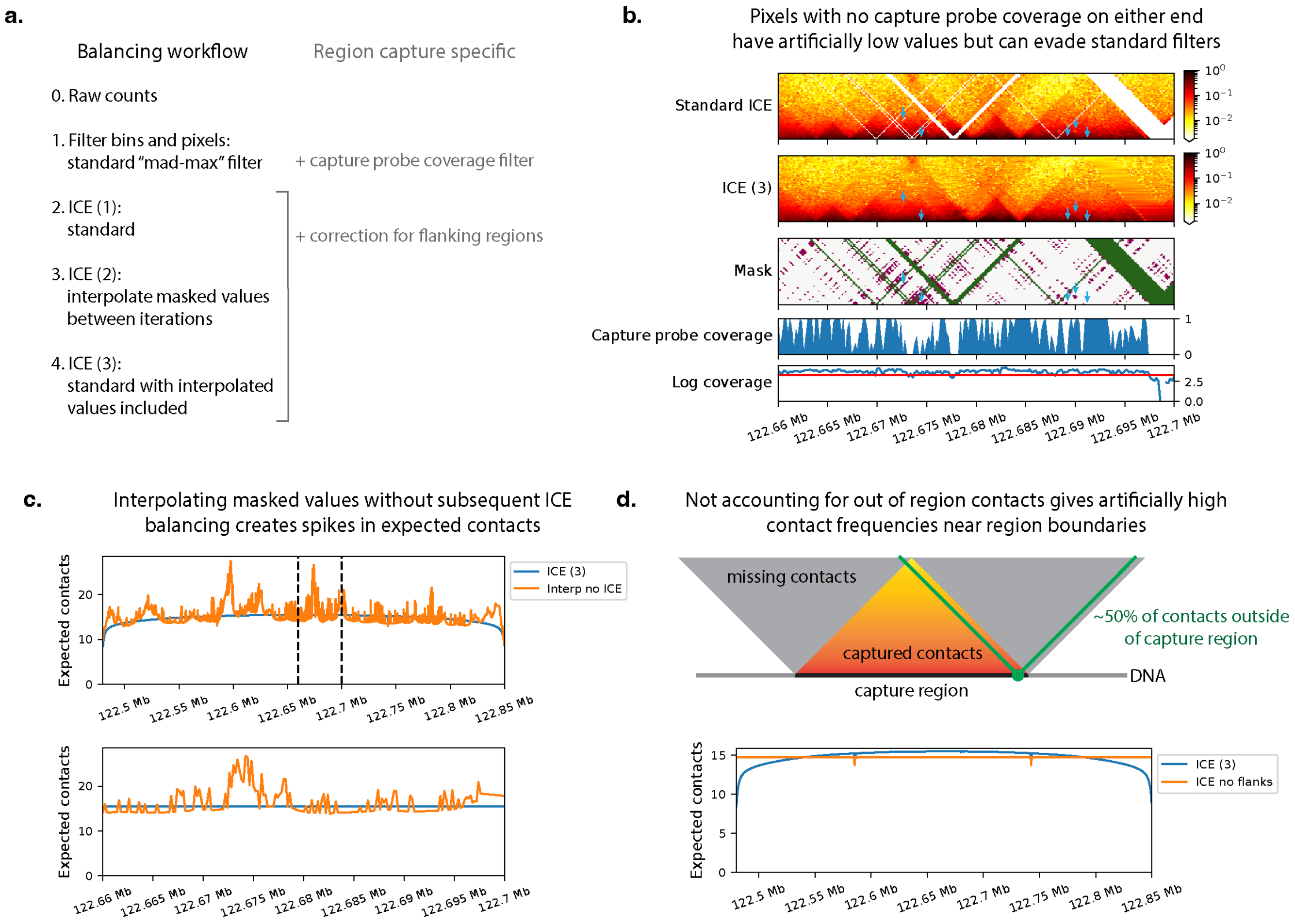

Fig. S2: **An augmented iterative correction balancing workflow.** (a) Outline of our workflow for micro-C data processing. (b) Lack of capture probe coverage can result in artifacts not accounted for using standard processing approaches. The capture probe coverage track shows the fraction of 200-bp windows that are covered by capture probes. The mask image shows pixels filtered based on read coverage in green and additional pixels filtered based on capture probe coverage in magenta. These pixels without capture probe coverage, highlighted by blue arrows, can result in holes in the contact map using standard ICE balancing. These holes are removed using our procedure. (c) Typically, masked pixels of the contact map are treated as zeros in ICE balancing. However, this results in artificial variation in contact density if these masked values are subsequently interpolated because some windows interact more closely with masked pixels. This is problematic as our simulations effectively interpolate missing values. Our augmented ICE balancing procedure overcomes this issue. (d) Enforcing that the capture region has uniform coverage creates artificially high interaction frequencies near the boundaries. We account for this by filling in the missing contacts (gray) with the average value for the respective genomic separation.

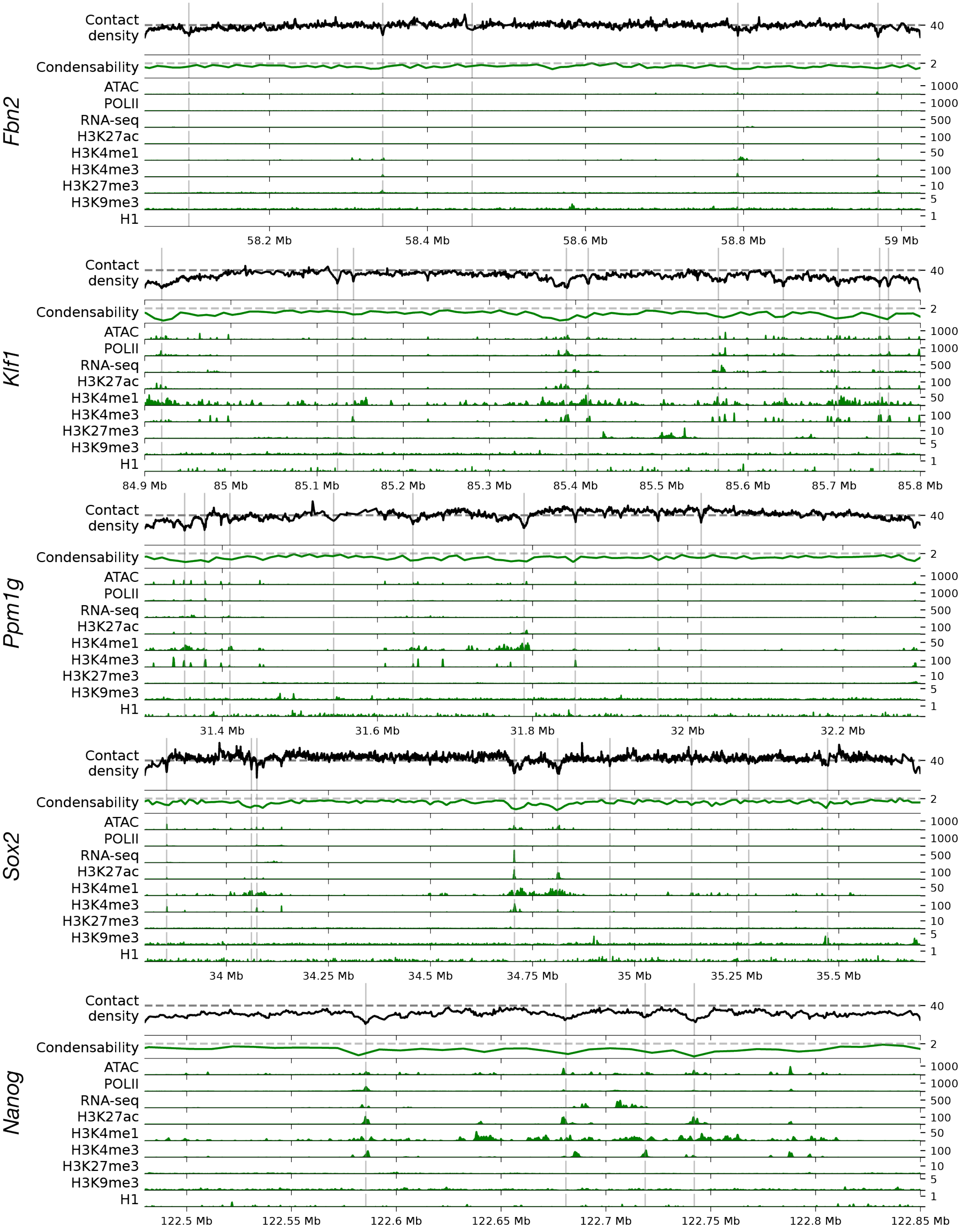

Fig. S3: **Contact densities predicted by neighbor balancing are correlated with epigenetics marks** Equivalent to Fig. 1c but showing all five regions and a larger repertoire of epigenetics tracks.

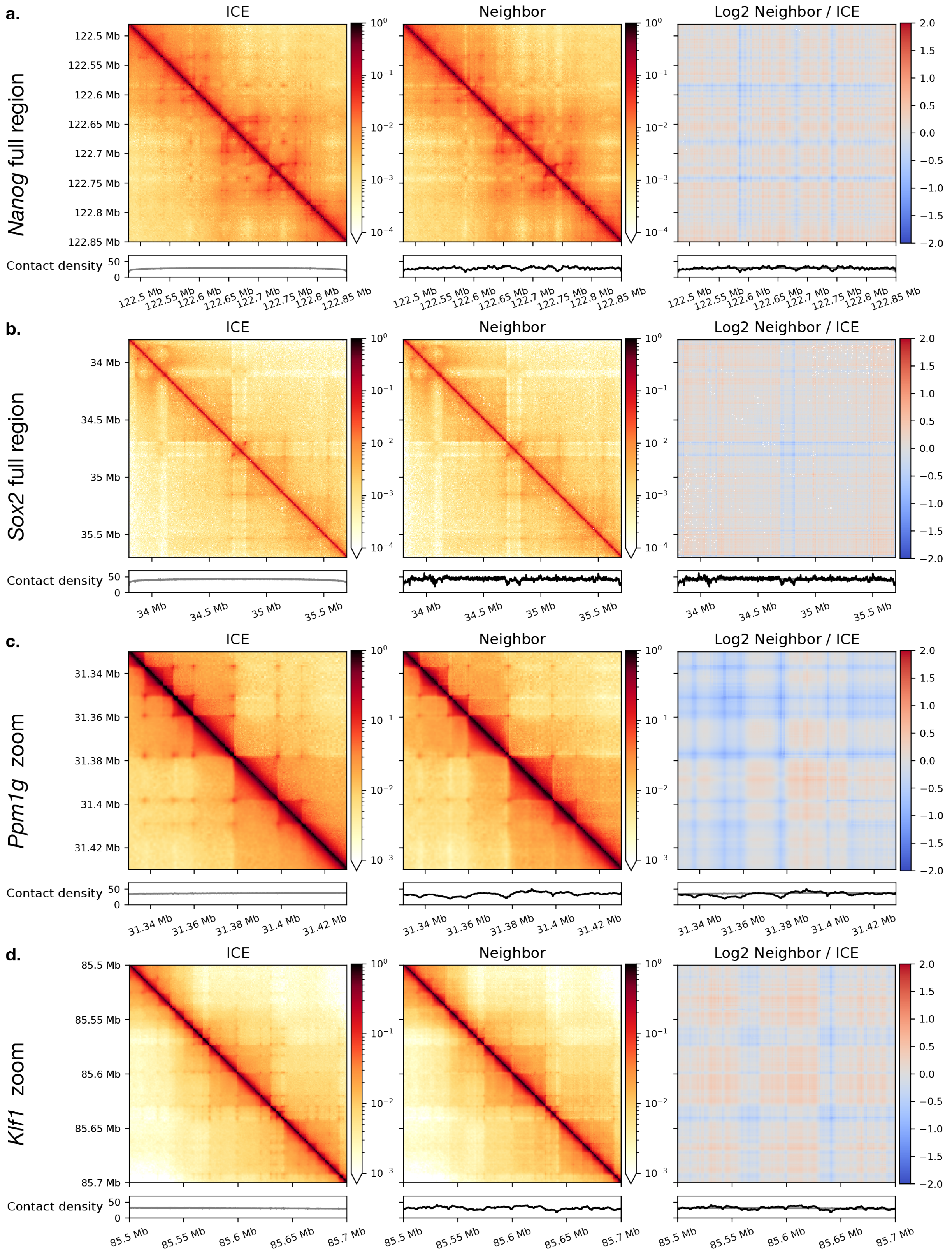

Fig. S4: **Enhancer–promoter interactions are reduced by neighbor balancing** (a, b) Contact maps for the full regions show that neighbor balancing generally reduces the strength of enhancer–promoter interactions. (c, d) Zoom ins of microcompartment containing regions show that neighbor balancing quantitatively decreases the strength of the interactions, but they remain clearly visible in the contact map.

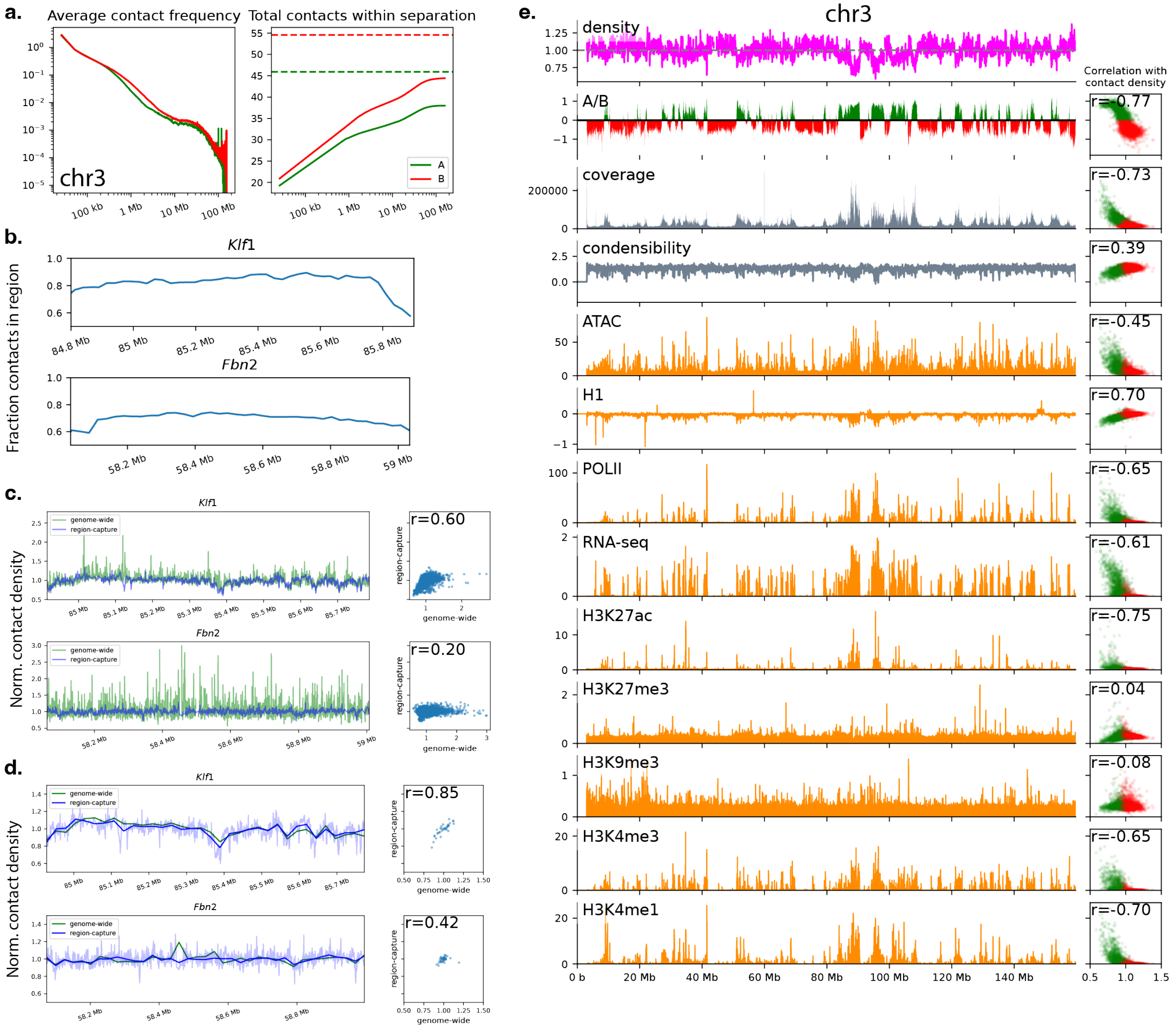

Fig. S5: **Neighbor balancing captures chromosome-scale variations in contact density.** Application of neighbor balancing to genome-wide micro-C for mouse embryonic stem cells from Hsieh et al. [26]. (a, left) P(s) curve for chromosome 3 for regions with an A/B compartment score greater than 0.2 (green) and less than -0.2 (red). (a, right) The contact density considering interactions up to the indicated genomic separation. The dotted line shows the contact densities including trans contacts. (b) The fraction of total micro-C contacts inside of the simulated regions. Simulating 1 MB regions generally results in approximately 75% of contacts being included. (c) Normalized nucleosome-resolution contact densities computed using region-capture micro-C data and genome-wide data are largely consistent. However, those computed with genome-wide data are substantially noisier, highlighting that RCMC is required to achieve nucleosome resolution. (d) When binned at 25.6 kb resolution, the normalized contact densities become smooth and are in close agreement. However, viewing the contact densities at this coarse-resolution smooths away the sharp dips in contact density at active regulatory elements, e.g. the dip near 85.4 MB in the *Klf1* region. (e) Comparison of normalized contact densities to a variety of epigenetics tracks.

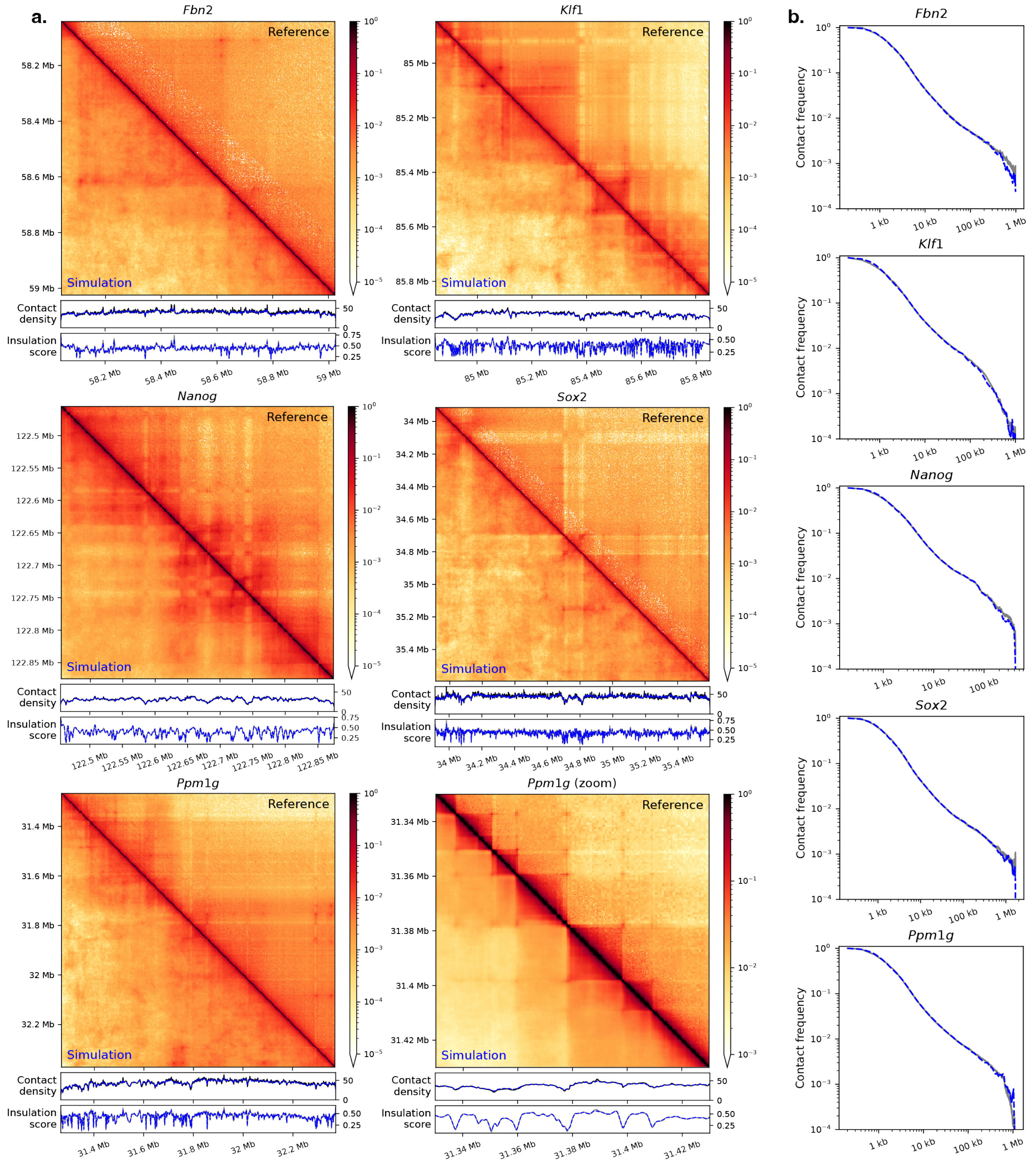

Fig. S6: **Simulations successfully reproduce experimental contact maps.** (a) Simulated contact maps are shown in the lower triangle and reference RCMC contact maps are shown in the upper triangle. Below, the contact densities and insulation scores using a window size of 10 nucleosomes (as used in our definition of clutches) are shown for simulation (blue) and reference (black). The bottom right plot shows a zoom in of the *Ppm1g* region highlighting that our simulations capture fine-scale microdomains and microcompartments. (b) P(s) curves for simulation (blue) and reference RCMC maps (gray).

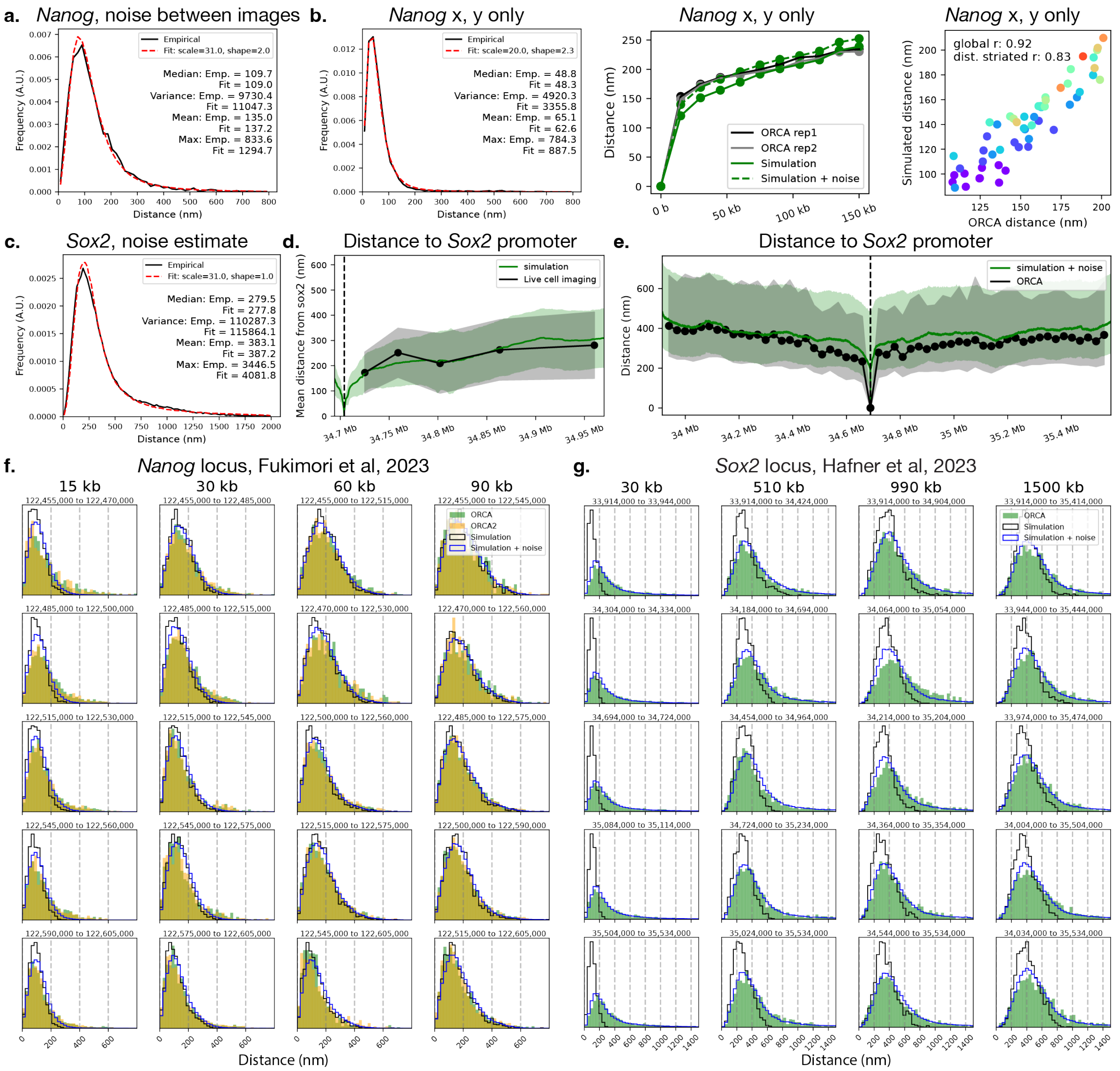

Fig. S7: **Simulations match fluctuations in pairwise distances.** (a) Estimated measurement noise fit to data where a single region is imaged twice. The black line shows the empirical distribution of distances between images; if there was no measurement noise, these distances would be zero. The red line shows the distance between images expected under the fit noise distribution. (b) The x, y coordinates of the *Nanog* ORCA data contain substantially less measurement uncertainty. From left to right, here we show equivalents to (a), Fig. 3a, and Fig. 3b, using only the x, y coordinates. Distances are multiplied by $\sqrt{3/2}$ to approximate 3D distances. (c) No control data was available for *Sox2*, so we fit the measurement uncertainty to match the experimental distribution of distances between regions separated by 60 kb. The empirical distribution of 60 kb distances from ORCA are shown in black and distribution of 60 kb distances from simulations with noise added are shown in red. (d) Comparison of distances to *Sox2* from simulations to live cell imaging from Li et al. [11]. The solid lines show the mean and the shaded region encloses the standard deviation. (e) Comparison of distances to *Sox2* from simulations to ORCA [10]. The solid lines show the median and the shaded region encloses the 20th to 80th percentile. (f) Example pairwise distance distributions for the *Nanog* locus using only x, y coordinates. ORCA and ORCA2 represent two replicates of the ORCA experiment. (g) Same for the *Sox2* locus but using all coordinates.

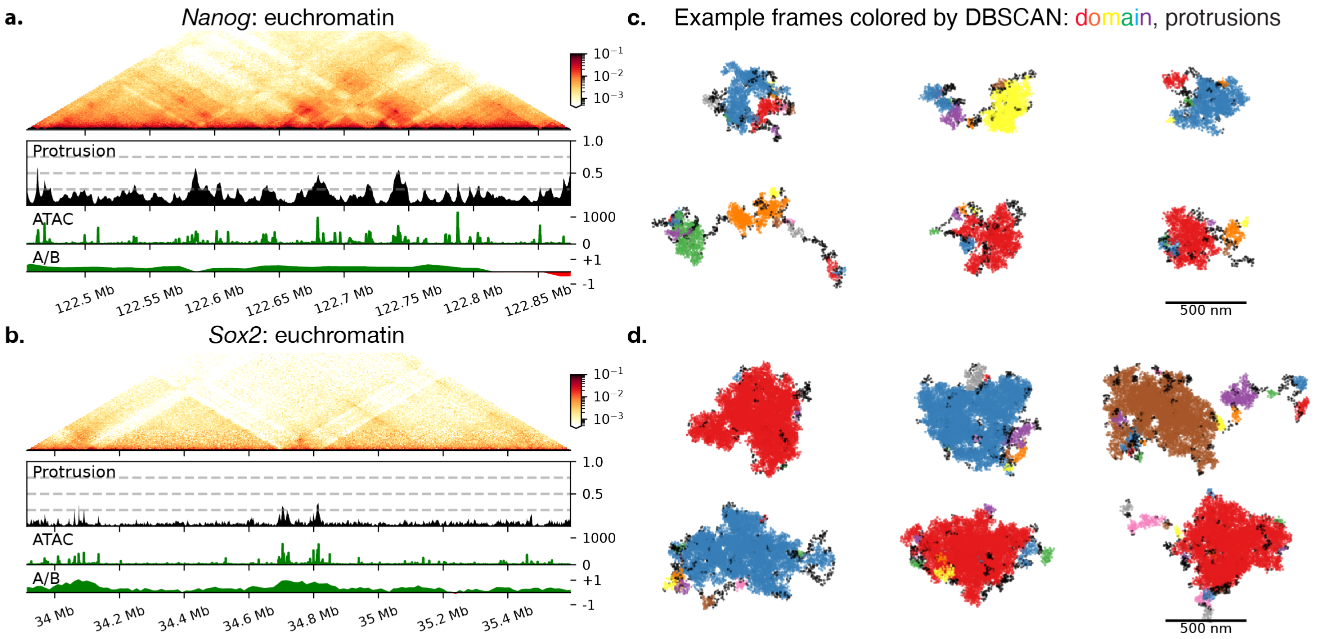

Fig. S8: **Supporting both heterochromatin and euchromatin are largely comprised of condensed domains.** Equivalent to Fig. 3a-f for the remaining two systems. Note that the *Sox2* system is 1.6 MB and the *Nanog* system is 0.37 MB, whereas the remaining systems are approximately 1 MB.

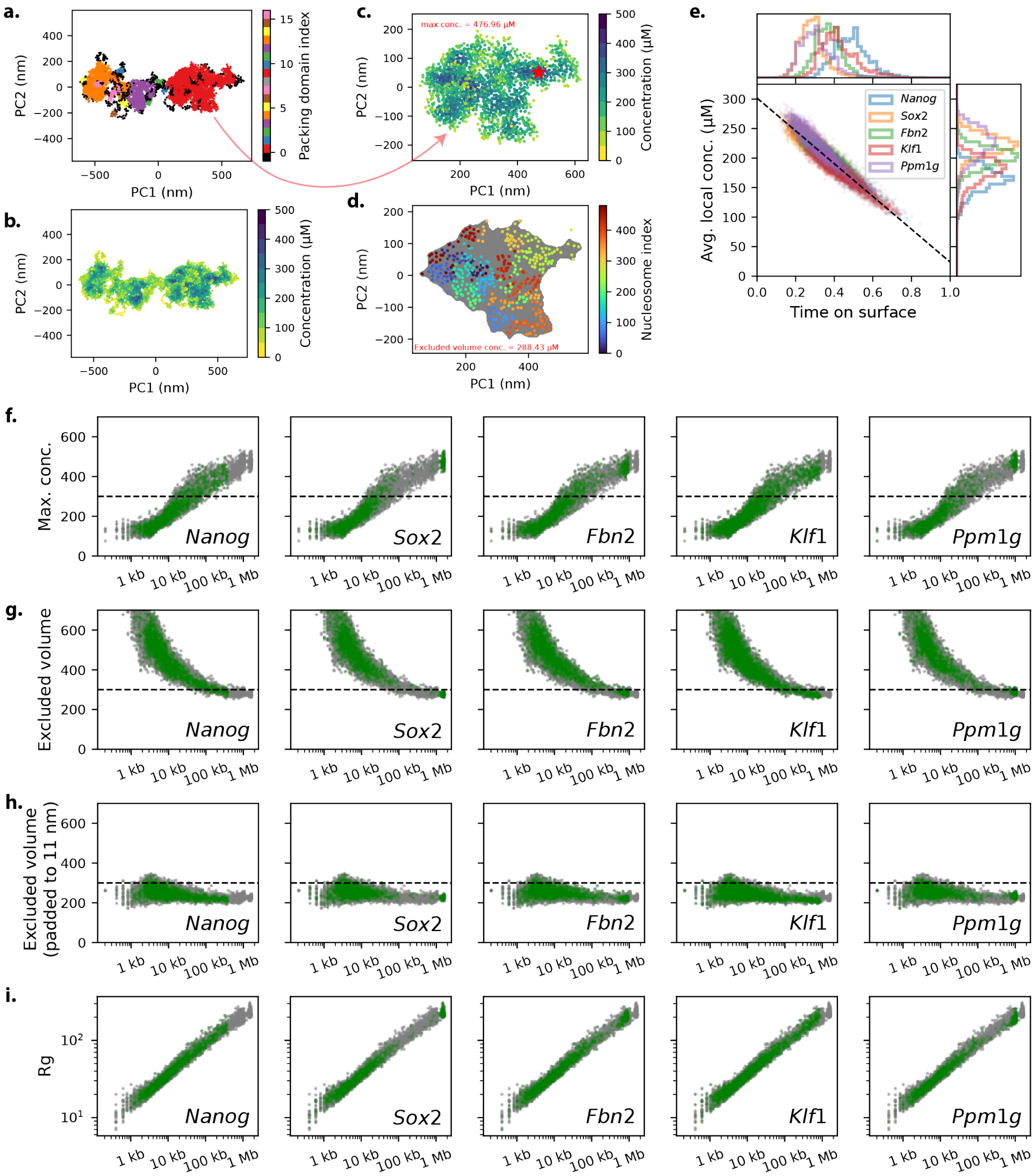

Fig. S9: **Packing domain densities are consistent across regions.** Example of density computation using the maximum local density and probe excluded volume. (a) An example simulation frame colored by packing domain assignment. (b) The same simulation frame colored by local density, i.e. the density inside a sphere with a radius of 40 nm centered on each nucleosome. (c) A zoom in to a single packing domain. Nucleosomes in the interior often have local densities in excess of 400 µM. (d) Illustration of probe excluded volume. The probe excluded volume of a 20 nm slice is shown in gray, and nucleosomes within a 31 nm slice are shown in rainbow colors. (e) The time a nucleosome spends on the surface explains much of the variation in average local density. A nucleosome is considered on the surface if it can be contacted by a 20 nm radius probe. This includes most nucleosomes in protrusions and those on the surface of packing domains. While no such cases are present, extrapolating this relationship to a nucleosome that spends no time on the surface suggests an average nucleosome concentration of 300 µM. (f) Packing domain densities computed using the maximum local density. Densities for all systems are plotted in gray and densities for the indicated system are shown in green. 300 µM is shown as a dotted black line for reference. (g) Same for the probe excluded volume. (h) Same for the probe excluded volume with nucleosome radii padded to 11 nm. (i) Same for the radius of gyration. Overall

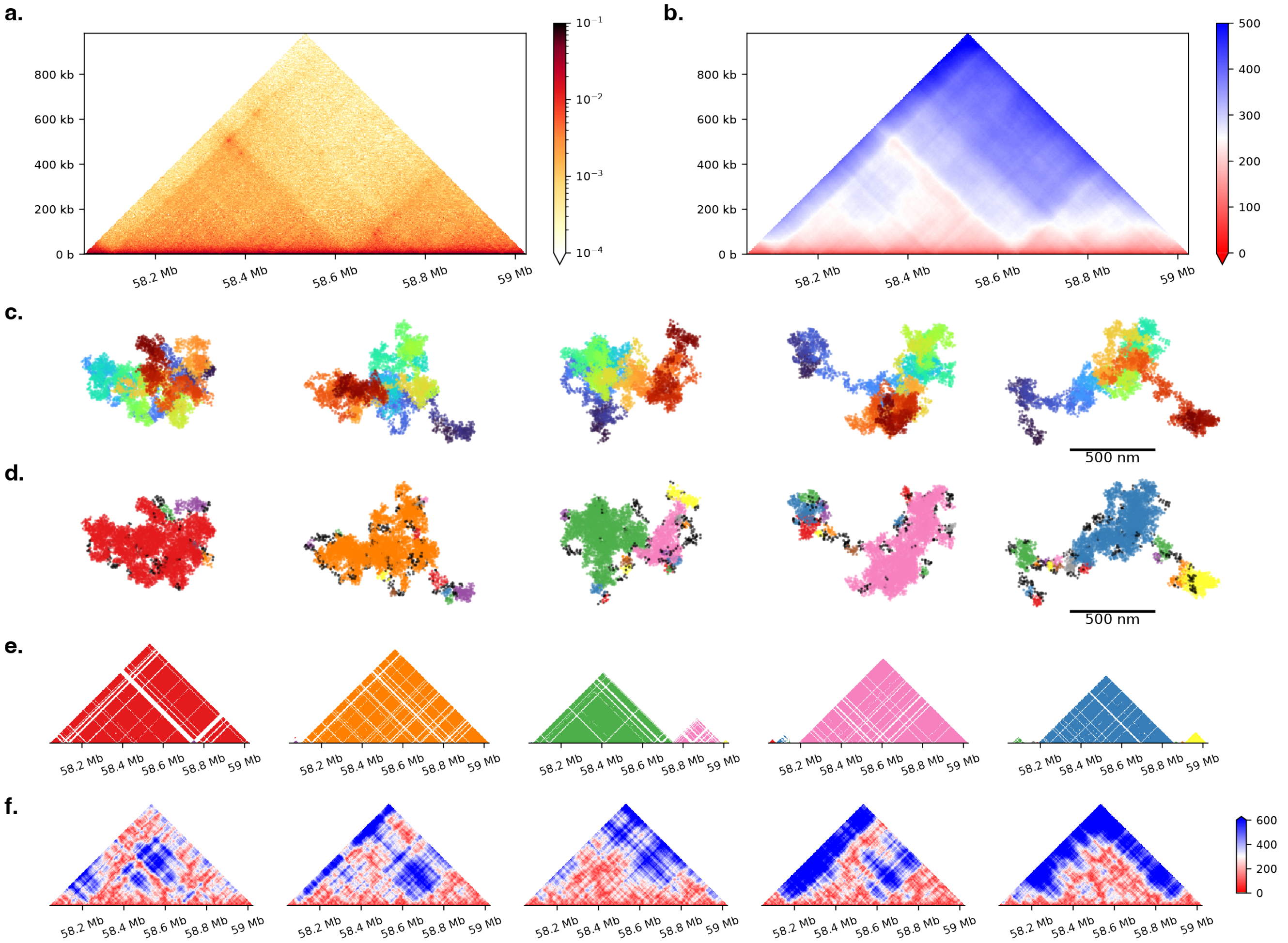

Fig. S10: **Simulations capture the stochastic character of packing domains.** (a) The reference contact map for the *Fbn2* region, shows that the region is decomposed into three TADs: one complete and two partial. (b) Median simulated distances show that nucleosomes within TADs are indeed on average closer to each other than nucleosomes in different TADs. However, in individual simulation frames, the allotment of this region into packing domains is highly varied. Representative simulation frames colored by chain index (c) and packing domain assignments (d). (e) An illustration of the genomic content of each packing domain. (f) Per-frame distance maps.

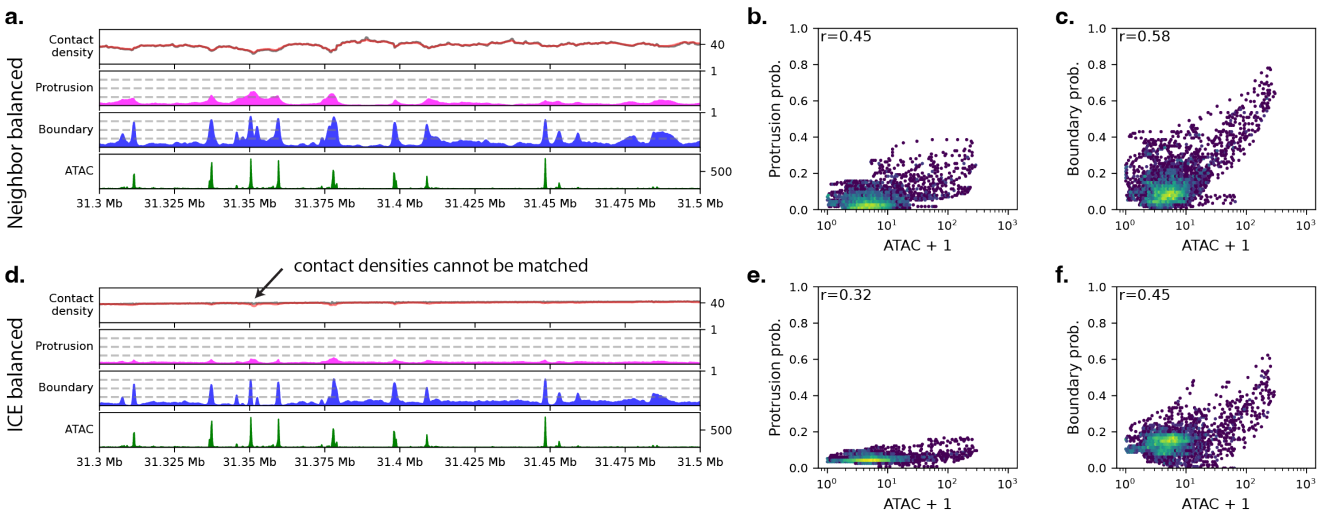

Fig. S11: **Simulations using ICE balanced contact maps.** (a, d) Reference (gray) and simulated contact densities, protrusion probabilities, clutch boundary probabilities, and ATAC-seq coverage for a portion of the *Ppm1g* region from simulations using a neighbor balanced and ICE balanced contact map, respectively. (b, e) Correlations between ATAC-coverage and protrusion probability. (c, f) Correlations between ATAC-coverage and clutch boundary probability. When using ICE balanced contact maps, the contact densities are largely constant and accordingly, there is little variation in the protrusion probability. There is still some variation because neighbor contact frequencies cannot be optimized (as the bond length is largely fixed) and ICE balanced data contains substantial variation in the neighbor contact frequencies. In particular, ATAC peaks generally have higher values in the first off-diagonal, leading to deviations between the reference contact density and the simulated local density.

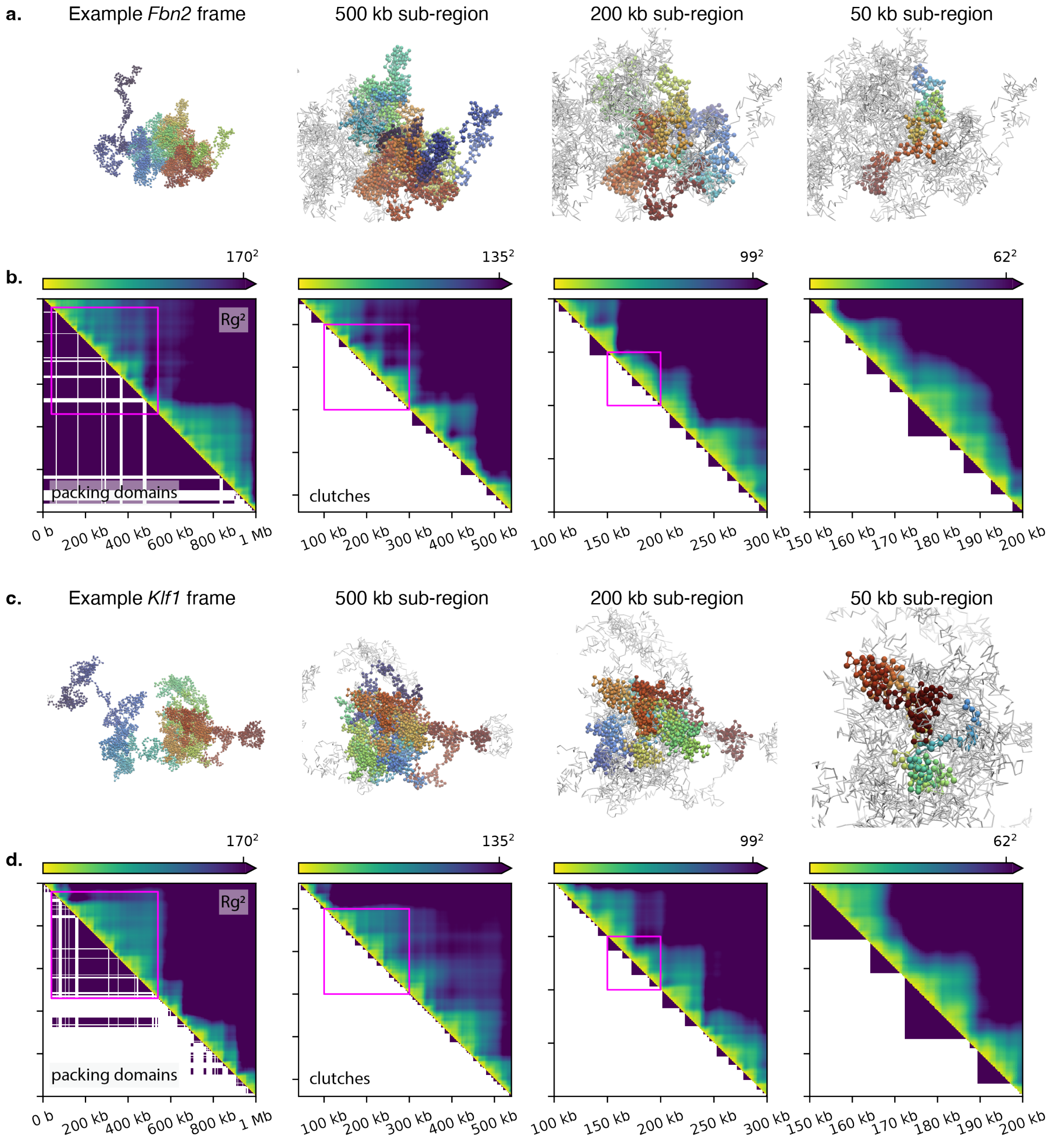

Fig. S12: **Chromatin has hierarchical structure across a broad range of scales.** (a) Example simulation frame for the optimized *Fbn2* system. Each panel shows increasingly smaller sub-regions in rainbow colors with the beginning of the region in red and end in purple and the remainder of the system in white. (b) Comparison of the squared radius of gyration ($Rg^2$) to packing domain and clutch assignments for the simulation frame shown above. Each pixel of the upper triangle of each panel represents the $Rg^2$ for the corresponding segment of the system, i.e. the ijth pixel is the $Rg^2$ for nucleosomes i through j. The lower triangle shows the packing domain assignments (left panel) or clutch assignments (remaining panels). Self-interacting domains are present at all scales. Clutches, though defined using insulation scores, not $Rg$, represent a slice of this hierarchical structure. (c, d) Equivalent for an example simulation frame for the *Klf1* system. This figure was inspired by Figure 2F from Tan et al. [27]. They observe a fractal globule at greater than MB scales [28]. Our model similarly shows self-interacting domains down to the kilobase scale. However, at small scales, sub-regions expand rapidly within packing domains.

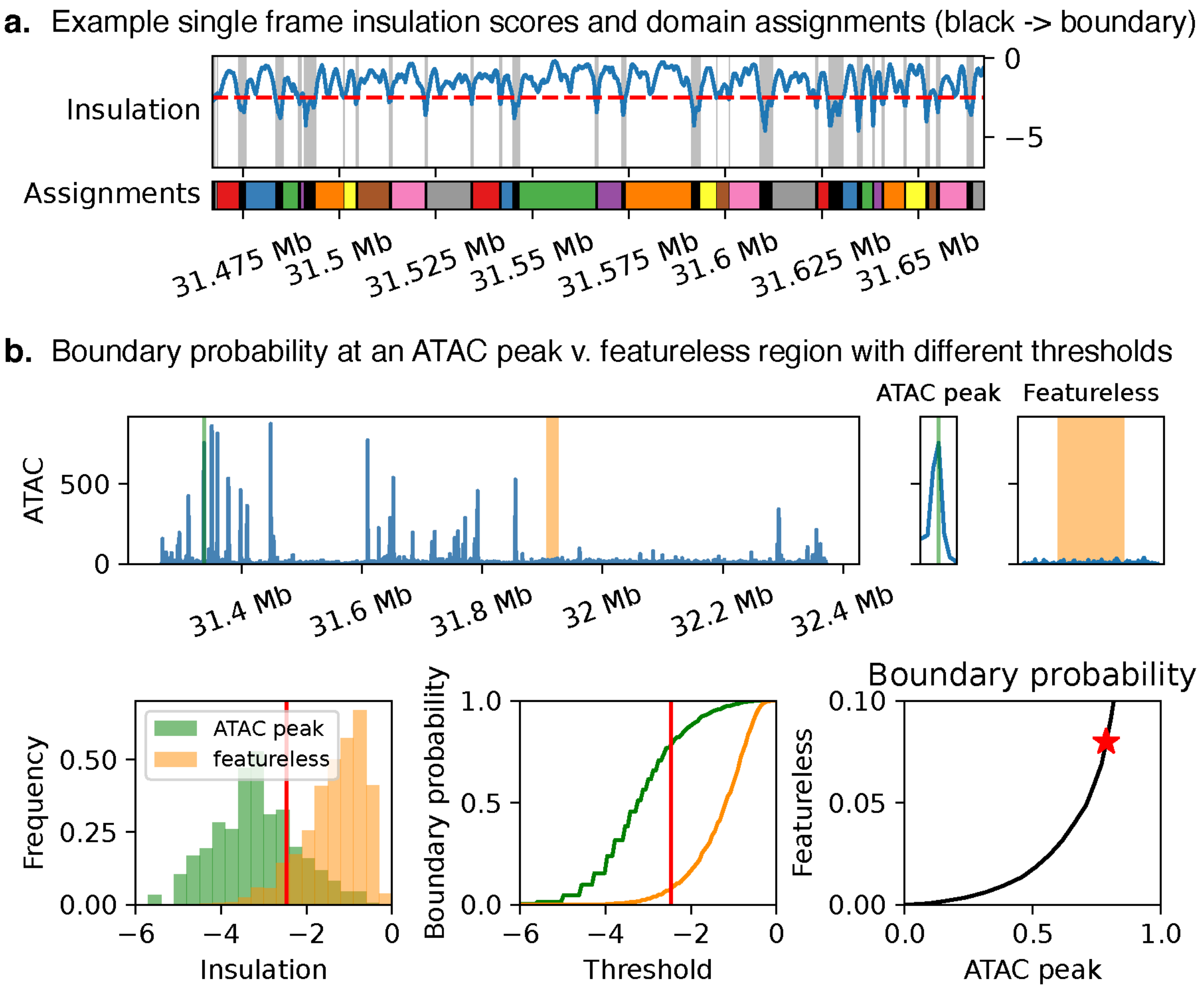

Fig. S13: **Impact of threshold choice in insulation analysis.** (a) Illustration of clutch assignments. (b) Here we compare the boundary probabilities at an ATAC peak to those in a largely featureless region. We find that the insulation scores at the ATAC peak are generally lower than in the featureless region–consistent with this region being more locally extended. However, there is significant overlap between the distributions, supporting that meaningful clutch boundaries also occur spontaneously in featureless regions.

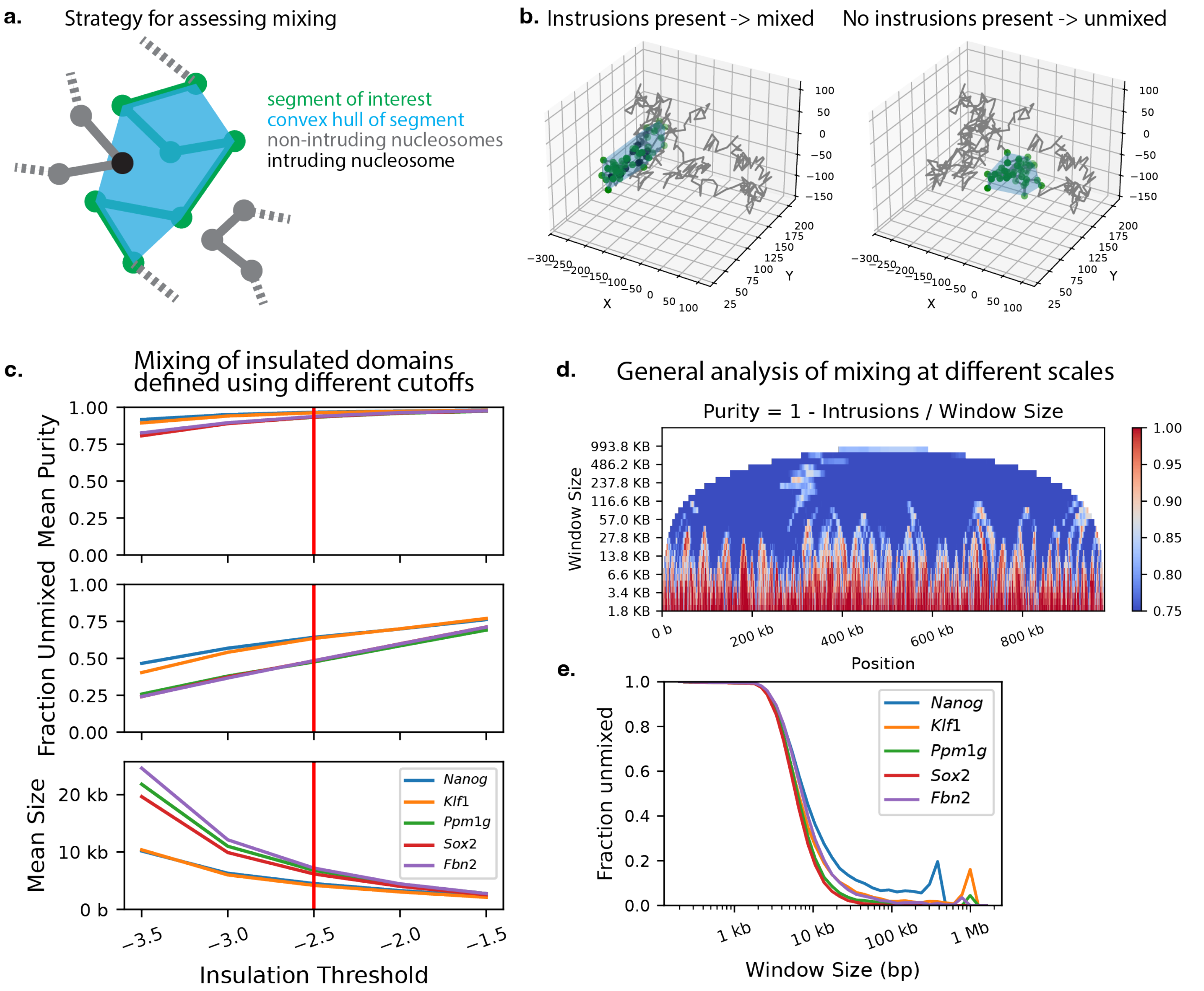

Fig. S14: **Clutches are largely unmixed.** (a) The extent to which a segment of interest is mixed with the rest of the system was assessed by counting the number of other nucleosomes inside the convex hull of the segment of interest. (b) A segment is considered mixed if any such intrusions are present and unmixed if no intrusions are present. We additionally define the "purity" of a segment as one minus the number of intrusions divided by the size of the segment of interest. (c) Clutches have a high purity (top) and are unmixed roughly half of the time (middle). Choosing a stricter insulation score threshold would have resulted in bigger (bottom) and more mixed clutches (middle). (d) The purities for segments of various sizes centered on each nucleosome. (e) The fraction of unmixed segments decays with increasing segment size. The window size at which the fraction unmixed is 0.5 is about the size of clutches.

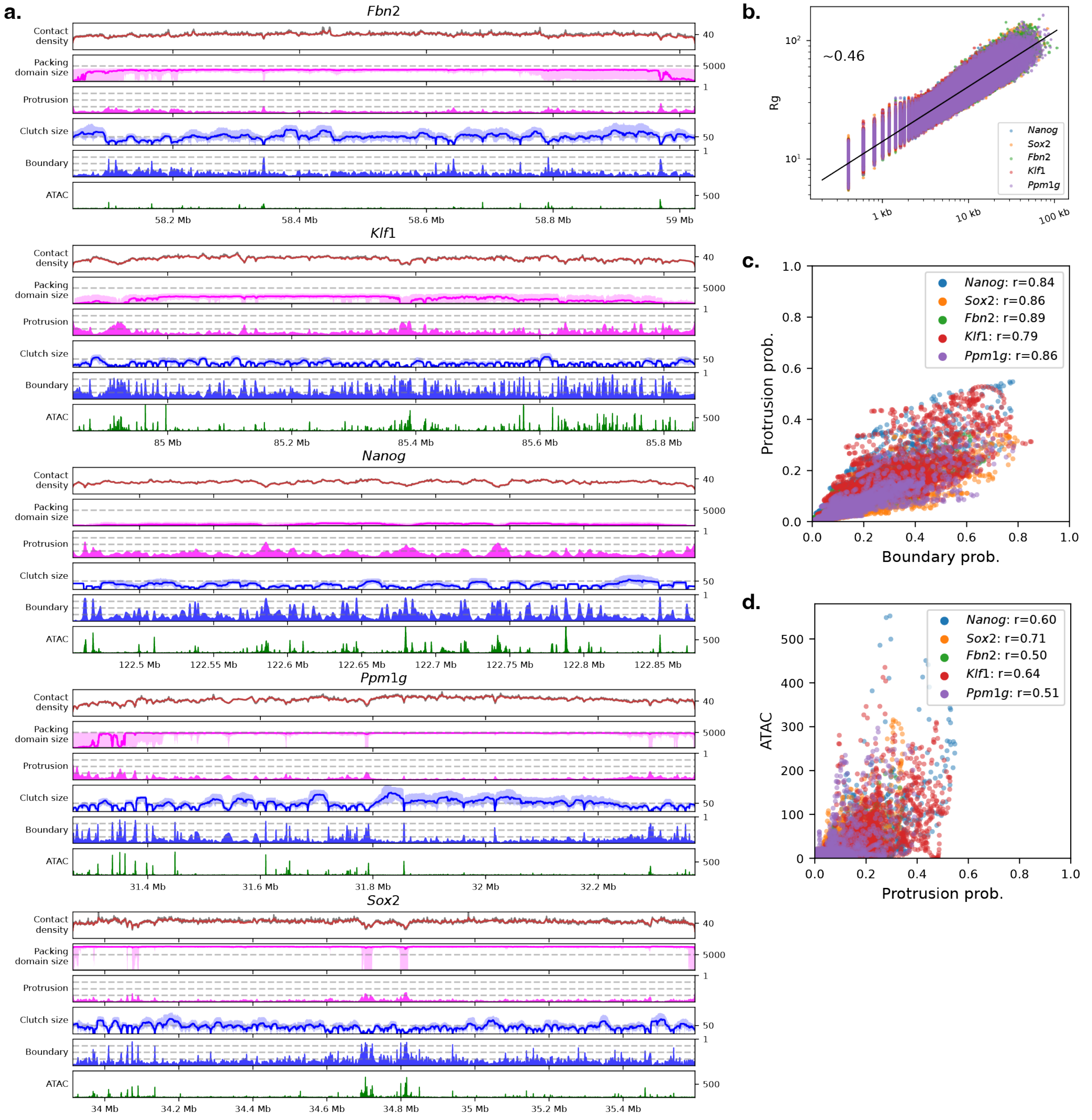

Fig. S15: **Relationship between clutches, packing domains, and chromatin accessibility.** (a) Contact densities, protrusions, and clutch boundaries are highly correlated with chromatin accessibility, as assessed by ATAC-seq coverage. Moreover, both packing domains and clutches tend to be smaller around ATAC-seq peaks. The size of domains are plotted with the median as a solid line and the 25th to 75th percentile shaded. (b) The $Rg$ of clutches scales with an exponent of $\sim 0.46$, reflecting an elongated, fiber-like structure. (c) The probability of a position occurring as a protrusion or clutch boundary are highly correlated. (d) Protrusions are more likely at positions with high ATAC coverage.

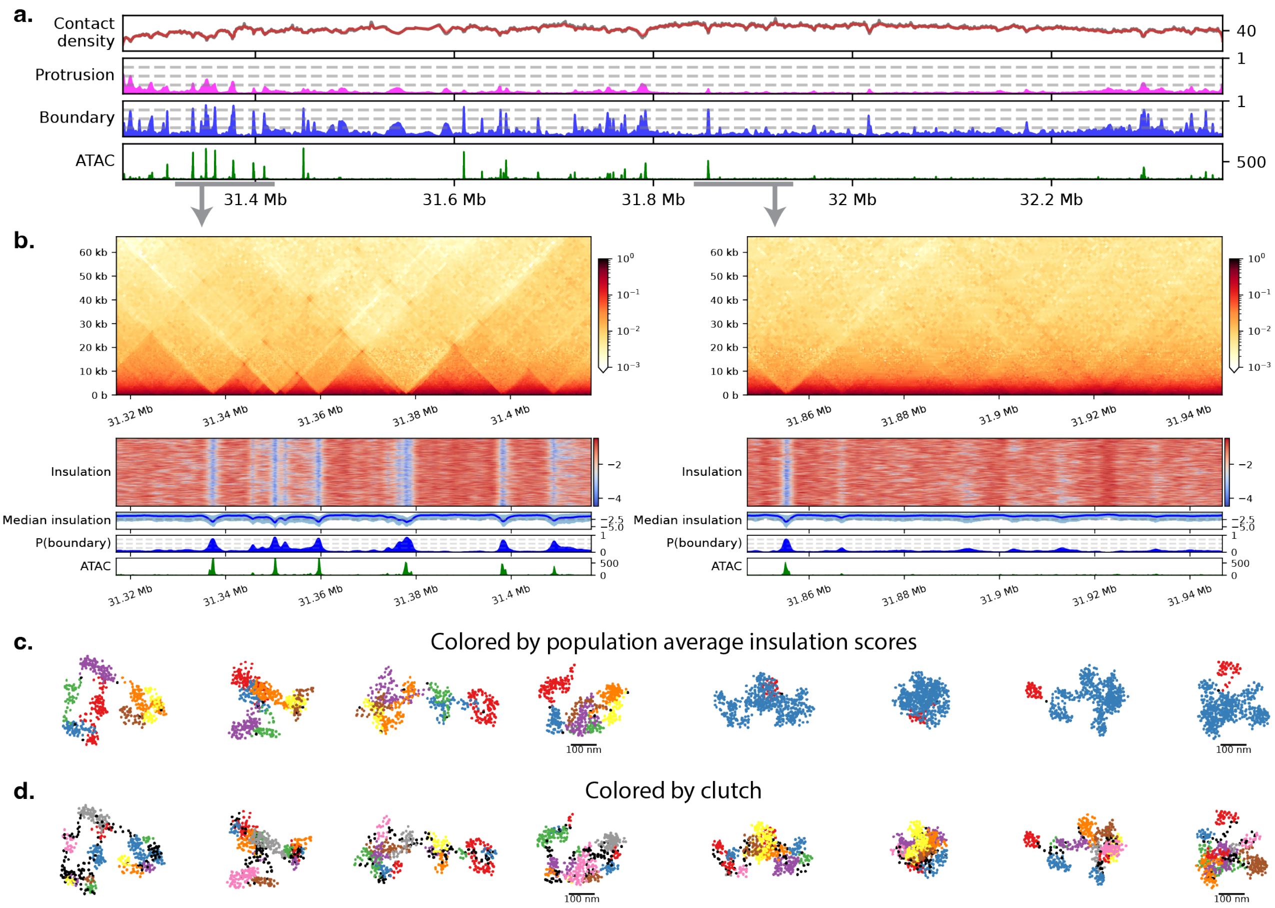

Fig. S16: **Well-phased clutches emerge as microdomains.** (a) Reproduction of Figure S15a for *Ppm1g* region to show context of highlighted regions. (b) Some regions show well phased clutch boundaries, typically at ATAC-seq peaks, and these boundaries result in microdomains (left). Each row of the heatmap shows insulation scores for a single simulation frame. In other regions, boundaries still occur, albeit at a lower rate, but they are positioned randomly, so do not appear in the contact map. (c) Representative simulation frames for the two regions colored by domains assigned as described for clutches but using the population average contact map. The microdomains largely remain unmixed, but often group together, contact each other on their surfaces. (d) The same frames colored by clutch (using the single-frame contact maps).

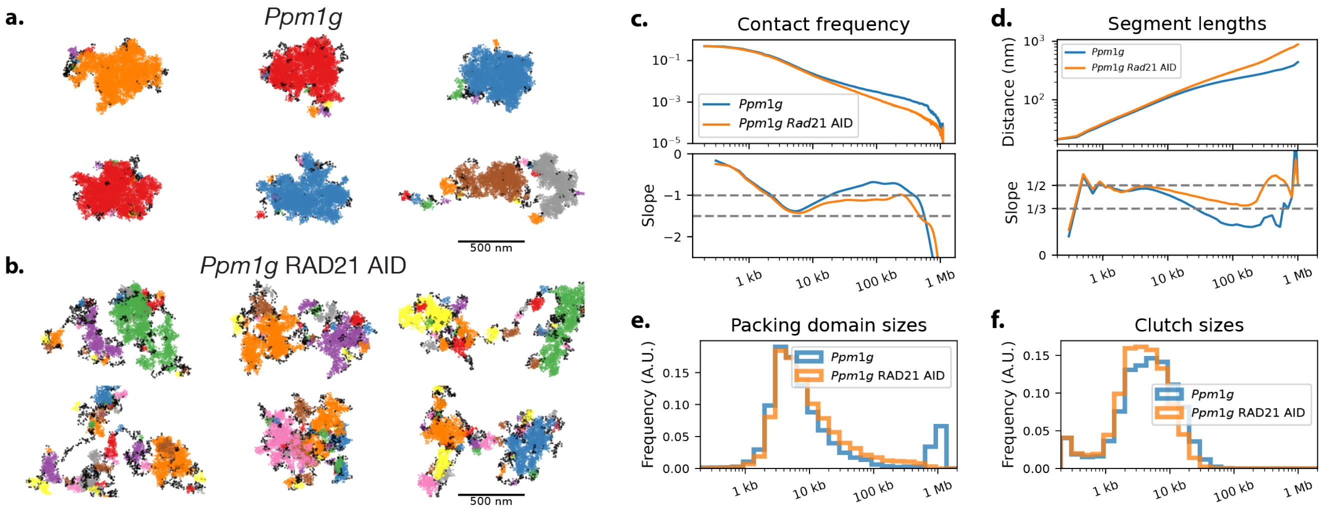

Fig. S17: **Upon cohesin depletion chromatin expands at large scales but local structure is maintained.** Representative simulation frames for the *Ppm1g* system using (a) WT RCMC data and (b) RCMC data in which RAD21 was depleted using an auxin inducible degron (AID) [29]. (c) Neighbor balanced RCMC contact frequency scaling, (d) segment length scaling, (e) packing domain sizes, and (f) clutch sizes for the two conditions.

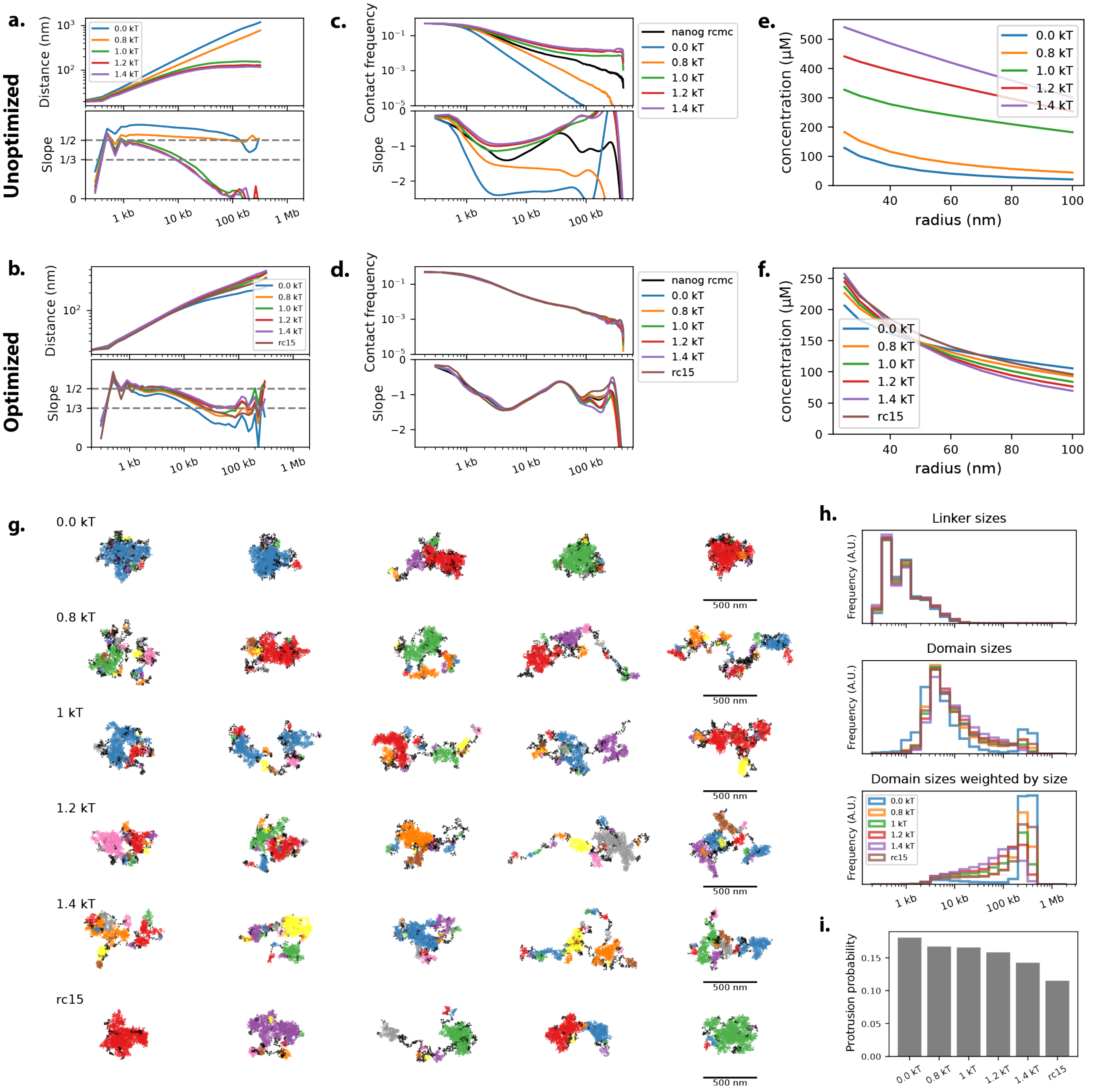

Fig. S18: **Altering the base nucleosome interaction energy changes scaling properties.** (a) Segment end-to-end distances for the base model with a range of interaction energies. The theta condition is achieved with an interaction energy between 0.8 kT and 1.0 kT. (b) Equivalent for the optimized *Nanog* system. The distances scale faster with higher interaction energies, but this trend begins to plateau with an interaction energy of 1.0 kT or higher. In the "rc15" simulations, we directly optimize the short range nucleosome interactions instead of the interaction potentials based on the contact indicator function. It proved challenging to completely reproduce the reference data with this strategy, but nonetheless is provides comparable values for our observables of interest as our standard simulations. (c, d) Simulated P(s) curves before and after optimization. (e, f) The average local nucleosome concentration, defined as the average concentration within the indicated radius of each nucleosome. Strong nucleosome interaction energies favor short range interactions, whereas weak interactions favor long range interactions. (g) Representative simulation frames colored by packing domain assignments for different nucleosome interaction energies. Protrusions are shown in black. To our eyes, conformations are qualitatively similar in all but the 0.0 kT condition. (h) Size of domains and protrusions for different interaction energies. (i) Nucleosome interactions have minor impacts on protrusion probabilities.
